# Cross-Species Integration of Transcriptomic Effects of Tobacco and Nicotine Exposure Helps to Prioritize Genetic Effects on Human Tobacco Consumption

**DOI:** 10.1101/2019.12.23.887083

**Authors:** Rohan H C Palmer, Chelsie E. Benca-Bachman, Jason A. Bubier, John E McGeary, Nikhil Ramgiri, Jenani Srijeyanthan, Spencer Huggett, Jingjing Yang, Peter Visscher, Jian Yang, Valerie Knopik, Elissa J. Chesler

## Abstract

Computational advances have fostered the development of new methods and tools to integrate gene expression and functional evidence into human-genetic association analyses. Integrative functional genomics analysis for altered response to alcohol in mice provided the first evidence that multi-species analysis tools, such as GeneWeaver, can identify or confirm novel alcohol-related loci. The present study describes an integrative framework to investigate how highly-connected genes linked by their association to tobacco-related behaviors, contribute to individual differences in tobacco consumption. Data from individuals of European ancestry in the UKBiobank (N=139,043) were used to examine the relative contribution of orthologs of a set of genes that are transcriptionally co-regulated by tobacco or nicotine exposure in model organism experiments to human tobacco consumption. Multi-component mixed linear models using genotyped and imputed single nucleotide variants indicated that: (1) variation within human orthologs of these genes accounted for 2-5% of the observed heritability (meta h^2^_SNP-Total_=0.08 [95% CI: 0.07, 0.09]) of tobacco/nicotine consumption across three independent folds of unrelated individuals (enrichment ranging from 0.85 - 2.98), and (2) variation around (5, 10, 15, 25, and 50 Kb regions) the set of co-transcriptionally regulated genes accounted for 5-36% of the observed SNP-heritability (enrichment ranging from 1.60 – 31.45). Notably, the effects of variants in co-transcriptionally regulated genes were enriched in tobacco GWAS. These findings highlight the advantages of using multiple species evidence to isolate genetic factors to better understand the etiological complexity of tobacco and other nicotine consumption.

## INTRODUCTION

Contemporary thought on genetic research of complex traits in humans is that large scale genome-wide association studies (GWAS) are required to identify reproducible single nucleotide polymorphism (SNP) associations that can lead to insights into biological systems that underpin a particular phenotype. The agnostic nature of GWAS, i.e., all SNPs being tested without bias, is a strength that allows for the identification of previously unrecognized biological underpinnings. However, the GWAS approach is not without limitations. For example, examination of genome-wide variation requires a stiff penalty for multiple comparisons leading to the need for increasingly large sample sizes. The requirement of sample sizes in the 100’s of thousands to millions (i.e., mega-GWAS) exerts pressure on the depth of phenotyping that may be done (i.e., more intensive and costly phenotypes are untenable for Mega-GWAS studies). Additionally, SNPs implicated by GWAS are not always readily associated with gene function. In fact, a majority of GWAS hits fall in non-coding or intergenic regions^1^. Linkage disequilibrium allows for a relatively sparse coverage of the genome to be maximally informative, but simultaneously limits the immediate “translatability” of the signals (i.e., a SNP identified by GWAS may be a proxy for a causative SNP some genomic distance away). In sum, while GWAS findings have become increasingly reproducible as sample sizes increase, it has become increasingly evident that additional sources of data (e.g., gene regulatory and epigenetic data^2^) are needed to understand how subtle SNP effects increase risk for pathology or can be utilized in identifying critical biological mechanisms.

Genetic studies of tobacco consumption assume that genetic variation in the biological sample collected (e.g., blood and saliva) reflects the genetic influences in brain that mediate the psychoactive properties of nicotine and other chemicals found in tobacco products. Nicotine has been shown to cause changes in neural organization, particularly in the brain’s reward systems, psychomotor and cognitive processes via its ability to interact with nicotinic acetylcholine receptors (nAChRs).^3; 4^ By altering neural circuits, especially those comprising the dopaminergic systems of the midbrain, nicotine elicits a high potential for addiction, regardless of the form in which it is marketed.^5^ Altogether, these properties of tobacco products highlight putative genetic mechanisms that may mediate consumption. The largest tobacco consumption meta-GWAS, to date, has identified 566 genetic variants in 406 loci associated with various phenotypes related to tobacco consumption (i.e., initiation, cessation, and heaviness of use).^6^ While the individual effects of these loci are limited, their application in the form of polygenic risk scores (PRS; i.e., the sum weighted effect of genome-wide variants that have been shown to predict individual differences on a trait) has been shown to have some utility in predicting consumption in similarly ascertained samples.^6^ Moreover, the variation in predictive utility of a PRS based on how the polymorphisms included are selected (e.g., p-value thresholds versus Best Linear Unbiased Predictors) underscores the need for additional lines of evidence to prioritize a subset of genome-wide signals contributing to consumption. However, short of increasing sample sizes to realize shared cumulative variant effects across subgroups of tobacco users in a GWAS, there are few methods to increase power to realize other genetic variants.

One approach to increase power in GWAS is the use of prioritized subsets of genomic variants while correcting for the overall genome-wide false discovery rate (FDR) using a multivariate mixed linear modeling framework. Indeed, the use of mixed models and prioritized subset approaches that fit multiple single nucleotide polymorphisms (SNPs) simultaneously have been shown to account for variation in a trait and improve power in association analyses.^7^ The recent development and application of genomic-relatedness-matrix restricted maximum likelihood (GREML^8; 9^) to addiction phenotypes and other complex traits, provides a multivariate framework so that the joint effects of loci can be determined. Moreover, GREML enhances power to localize the source of genetic variance for complex traits by aggregating the effects across *a priori* defined regions or categories of SNPs while accounting for LD.^10^ For instance, we applied GREML to Heroin Dependence and showed that SNPs in the 1-10% MAF range largely contribute to the known additive genetic variance even while controlling for LD.^11^ Similarly, Brazel et al., demonstrated that exonic rare variants in and around common variants are capable of indexing upwards phenotypic and genetic variance of alcohol and nicotine consumption, respectively, albeit with varied effects across phenotypes.^12^

While there have been several advances in application of genome-wide addiction genetics, overcoming the limitation of how to integrate prior knowledge and prioritize genomic variants, outside of broad functional categories (e.g., 3’ UTR, Intergenic, Rare coding, etc.), remains a critical limitation. Furthermore, lack of ready access to brain tissue in a living intact human precludes a direct understanding of tissue-specific epigenetic and/or expression differences that arise from continued exposure, which would aid in localizing expression quantitative trait loci sensitive to drug processes. In light of these concerns, the intuitive appeal of human-only genetic analysis is diminished, and suggests that another compelling approach is the use of complementary genomic data from model organism systems.

In this study, we evaluate the possibility of bridging between human GWAS and model organism genomics using a novel and integrative framework to answer the empirical question as to whether or not findings from model system studies may be leveraged with the human GWAS approach to speed advancements in this area. We used transcriptome-informed exposure models of tobacco/nicotine to parse genome-wide SNP-heritability estimates to test this hypothesis directly. This was achieved using the GeneWeaver heterogeneous functional genomics repository and analysis system as the primary platform for integration of evidence^13^ across existing studies.

## MATERIALS & METHODS

### Building an a priori network of genes co-transcriptionally regulated by nicotine

A gene set for nicotine consumption was identified using GeneWeaver ^14; 15^, a genomics data repository and analysis system. GeneWeaver integrates data from numerous databases, such as NCBI and ENSEMBL, various model organism databases (e.g., the Mouse and Rat Genome Databases, and the Zebrafish Model Organism Database) and genomic experimental results from the literature to produce curated sets of genes that can be analyzed using a suite of analytical tools.^13^ GeneWeaver was specifically designed for integration of genomic evidence and comprises over 199,664 gene sets spanning studies across 10 species. Using GeneWeaver we identified genes of interest from several *Mus musculus, Rattus norvegicus*, and *Danio rerio* functional genomics (typically microarray) experiments. As of October 2019, relevant data from no other species were identified upon review of the current literature and archived experimental sets available in GeneWeaver. Figure 1 outlines the protocol for establishing separate lines of evidence for each species.

**Figure 1.**
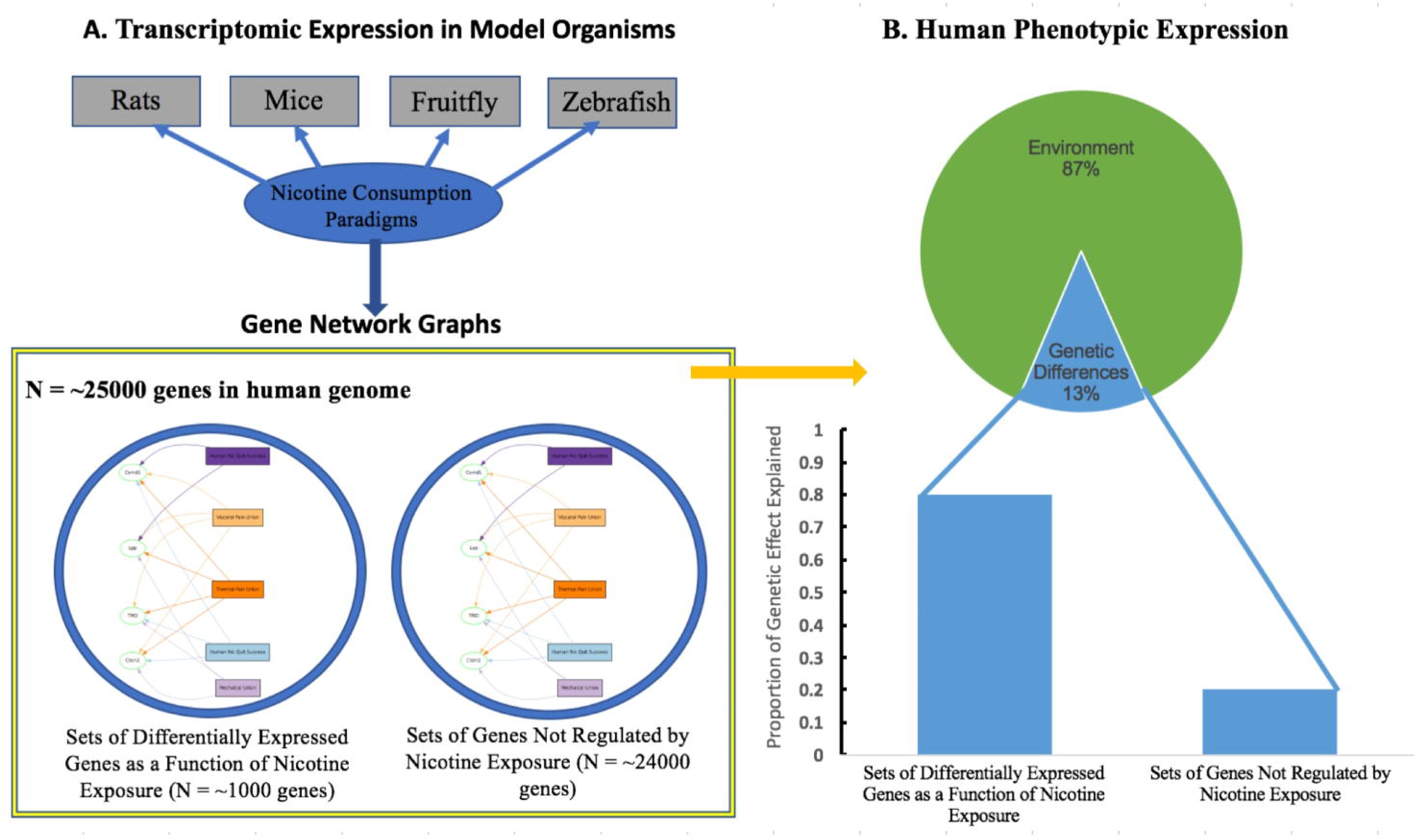
Theoretical integrative genomics approach to characterizing genetic underpinnings of nicotine consumption using model organisms.

We first identified studies by literature review or by shared summary statistics archived in the GeneWeaver system. Experimental studies were included if they provided differential expression or whole genome co-expression network analyses along with accessible summary statistics. Literature searches focused on exposure studies utilizing nicotine-specific model organism paradigms, including subcutaneous nicotine treatment, IVSA, nicotine delivered to the animal’s drinking water, and nicotine-induced conditioned place preference (see Table 1; no studies involving *Drosophila melanogaster* were identified which was most likely due to the fact that nicotine is a natural insecticide). Priority was given to weighted gene co-expression network analysis (WGCNA) studies to minimize inflation of the Type I error rate typically seen in QTL studies. Next, we merged studies with multiple reported gene sets (i.e., either by region, up/down regulated, or across time) to avoid inflating the replication threshold of individual genes. We then identified orthologous genes using GeneWeaver’s “Combine Gene sets” function which merges multiple gene sets into a single matrix while accounting for orthology across species; none of the identified studies were conducted in human samples.^13^ Lastly, identified gene sets were compared to the current human genome build (hg19) to localize relevant variants that were conserved across species; 712 orthologous genes were identified. Given the lack of a proof of principle for prospectively integrating model organism evidence in human studies we integrated the limited evidence across studies, especially given the minimal overlap between gene sets (Jaccard similarity ∼0.00-0.01; supplemental Table S1). Of these genes, 201 were replicated twice across GeneWeaver gene lists. For instance, ABL1 and GRIK2 were only observed in five brain regions from the Wang et al. study, but not observed in other studies. Supplementary Figure 1 provides a bipartite graph visualization of the 51 genes that were present in at least three gene lists. When collapsing across study and removing duplicates, 21 genes were observed across studies (see Supplementary Table S2). None of them overlapped across more than two studies. The analyses described below focused on SNPs in and around the 712 orthologous genes (GeneWeaver Gene Set ID: GS357552).

**Table 1.** Identified Geneweaver Gene Sets Related to Tobacco/Nicotine Exposure.

| Author(s) | GeneWeaver ID | Model Organism | Nicotine Consumption/Exposure Paradigm | Experimental Design | Brain Region | Number of Genes Contributed |
| --- | --- | --- | --- | --- | --- | --- |
| Chen et al.<sup>45</sup> | GS87128 | *Mus musculus* | Subcutaneous acute nicotine treatment (expression changes measured at time-points of 1, 2, 4, and 6 hrs) | Microarray Analysis, WGCNA | VTA | 184 |
| Polesskaya et al.<sup>46</sup> | GS14885 | *Rattus norvegicus* | Subcutaneous chronic nicotine treatment (at ages p25, p35, p45, and p55) | Microarray Analysis, qRT-PCR, Principle Cluster Analysis | PFC, Ventral Striatum, Hippo. | 66 |
| Wang et al.<sup>38</sup> | GS14888, GS14889, GS14890, GS14891, GS14892, GS14893 | *Mus musculus* | Nicotine administration in drinking water in two selectively bred mouse strains | Microarray Analysis, qRT-PCR, WGCNA | Amygdala, Hippo., nAcc, PFC, VTA | 651 |
| Kily et al.<sup>47</sup> | GS14902, GS14903 | *Danio rerio* | Nicotine-induced conditioned place preference | Microarray Analysis, qRT-PCR | Whole Brain | 158 |
| Sharp et al.<sup>34</sup> | GS128167 | *Rattus norvegicus* | Chronic nicotine self-administration | Microarray Analysis, RT-PCR | nAcc | 188 |

Table showing identified publications and GeneWeaver gene sets. Note: Gene set IDs can be used to review the full complement of genes supplied by each study.

### Fold creation and power calculation for the UK Biobank tobacco consumption sample

Hypotheses were tested using multiple subsets (i.e., folds) of the UKB data for computational efficiency and to demonstrate the robustness of the findings via replication as each dataset contained unrelated individuals. Analyses focus on the reported number of cigarettes by each participant (i.e., for prior and current smokers; nonsmokers were excluded). We identified 139,043 individuals of European ancestry as identified by principal components analysis and multidimensional scaling^16; 17^, who were no more related than second cousins and who also provided smoking data. The number of folds were determined *a priori* in order to maximize statistical power. The GCTA-GREML Power Calculator was used to estimate *a priori* power for sample sizes that provided at least 70% power to detect SNP-heritability estimates as small as one-third of 1% (0.333%).^18^ Power was based on the previously reported SNP-heritability and observed variance of the off-diagonal elements (∼6.68×10^−4^) in each fold.^6^ Consequently, the total sample was divided into three approximately equal folds (n_nic1_=41,263, n_nic2_=41,368, n_nic3_=41,213), each of which was made constitutionally equivalent by randomly sampling individuals from each quartile of the nicotine consumption distribution.

### Genotype quality control

Analyses focused on raw and imputed genotypes obtained using the Affymetrix UK BiLEVE Axiom and UK Biobank Axiom® arrays, which genotyped ∼850,000 variants (details available here: https://www.ukbiobank.ac.uk/scientists-3/genetic-data/). Quality control and imputation (to over 90 million SNPs, indels, and large structural variants) was performed by a collaborative group headed by the Wellcome Trust Centre for Human Genetics. Analyses focused on genotyped and imputed SNPs with good quality scores (r^2^ > 0.3). PLINK (version 1.9) was used to filter markers using the following criteria: genotyping rate >99%, minor allele frequency > 0.01, Hardy-Weinberg equilibrium p-value > 0.0001, and missing genotype rate < 0.10.^19^

### Regions-of-Interest Heritability Mapping

Given evidence for the import of intergenic variants in complex traits/disease, we partitioned the genetic variance of nicotine consumption into three regions-of-interest based on the list of genes acquired from the GeneWeaver database.^20^ As illustrated in Figure 2, the “gene” region was demarcated by the start and stop positions of each of the consumption genes. The flanking “buffer” regions of the genome were set to encompass the base pairs directly up-/down-stream of the 5’ and 3’ ends of each gene, respectively. We considered six buffer lengths in order to capture the effects of transcription factor (TF) binding sites whose exact position is unknown (0 kilo-base pairs (kb), 5kb, 10kb, 25kb, 35kb, and 50kb). Following marker extraction, the 5’ and 3’ variants for each buffer length were aggregated into a single buffer marker variant list for a given length. In addition, we examined the effects of all unselected variants (referred to as “other variants”), which belonged to regions of the genome that comprised SNP markers that were not within the parameters specified for the gene or buffer regions defined by the consumption gene set (Table 2 provides a count of the number of SNPs assigned to each component of the model). Consequently, the number of SNPs that comprised the “other variants” category varied depending on the length of the buffer regions.

**Table 2.** Estimated SNP-heritability for each component of the ROI model.

| *Model component* | *F1 $h^2_{SNP}$* | *F1 SE* | *F2 $h^2_{SNP}$* | *F2 SE* | *F3 $h^2_{SNP}$* | *F3 SE* | *$h^2_{meta}$ (95% CI)* | *% total $h^2_{meta}$* |
| --- | --- | --- | --- | --- | --- | --- | --- | --- |
| *ROI - Genes* |  |  |  |  |  |  |  |  |
| Gene (0kb buffer model) | 3.820E-03<sup>b</sup> | 1.64E-03 | 3.296E-03<sup>a</sup> | 1.63E-03 | 5.459E-03<sup>aaa</sup> | 1.79E-03 | 4.11E-3 [2.20E-3,6.00E-3] | 4.96% |
| Gene (5kb buffer model) | 2.865E-03<sup>a</sup> | 1.68E-03 | 1.540E-03 | 1.62E-03 | 4.184E-03<sup>aa</sup> | 1.86E-03 | 2.74E-3 [0.80E-3,4.70E-3] | 3.26% |
| Gene (10kb buffer model) | 1.720E-03 | 1.60E-03 | 8.890E-04 | 1.56E-03 | 2.933E-03<sup>a</sup> | 1.78E-03 | 1.76E-3 [-0.10E-3,3.60E-3] | 2.16% |
| Gene (25kb buffer model) | 1.260E-03 | 1.57E-03 | 1.070E-03 | 1.58E-03 | 2.773E-03<sup>a</sup> | 1.76E-03 | 1.60E-3 [-0.20E-3,3.50E-3] | 1.92% |
| Gene (35kb buffer model) | 1.340E-03 | 1.58E-03 | 1.310E-03 | 1.60E-03 | 2.995E-03<sup>a</sup> | 1.77E-03 | 1.81E-3 [<-0.01E-3,3.70E-3] | 2.16% |
| Gene (50kb buffer model) | 3.117E-03<sup>a</sup> | 1.61E-03 | 2.600E-03<sup>a</sup> | 1.59E-03 | 4.947E-03<sup>aa</sup> | 1.78E-03 | 3.50E-3 [1.60E-3,5.30E-3] | 4.17% |
| *ROI - Buffer* |  |  |  |  |  |  |  |  |
| Buffer 0kb | N/A | N/A | N/A | N/A | N/A | N/A | N/A | N/A |
| Buffer 5kb | 3.253E-03<sup>a</sup> | 1.78E-03 | 5.822E-03<sup>c</sup> | 1.88E-03 | 3.431E-03<sup>a</sup> | 1.86E-03 | 4.13E-3 [2.00E-3,6.20E-3] | 4.95% |
| Buffer 10kb | 7.880E-03<sup>c</sup> | 2.08E-03 | 8.964E-03<sup>c</sup> | 2.10E-03 | 7.717E-03<sup>c</sup> | 2.13E-03 | 8.19E-3 [5.80E-3,1.06E-2] | 9.84% |
| Buffer 25kb | 1.101E-02<sup>c</sup> | 2.35E-03 | 9.625E-03<sup>c</sup> | 2.27E-03 | 1.016E-02<sup>c</sup> | 2.39E-03 | 1.02E-2 [7.60E-3,1.29E-2] | 12.36% |
| Buffer 35kb | 1.135E-02<sup>c</sup> | 2.44E-03 | 9.542E-03<sup>c</sup> | 2.35E-03 | 1.048E-02<sup>c</sup> | 2.48E-03 | 1.04E-2 [7.70E-3,1.32E-2] | 12.50% |
| Buffer 50kb | 3.141E-02<sup>c</sup> | 5.29E-03 | 3.303E-02<sup>c</sup> | 5.28E-03 | 2.743E-02<sup>c</sup> | 5.32E-03 | 3.06E-2 [2.46E-2,3.66E-2] | 36.47% |
| *ROI - Other genomewide variants* |  |  |  |  |  |  |  |  |
| All Other Variants (0kb buffer model) | 7.145E-02<sup>b</sup> | 7.82E-03 | 7.586E-02<sup>c</sup> | 7.76E-03 | 8.853E-02<sup>c</sup> | 8.04E-03 | 7.84E-2 [6.95E-2,8.73E-2] | 94.92% |
| All Other Variants (5kb buffer model) | 6.923E-02<sup>c</sup> | 7.85E-03 | 7.235E-02<sup>c</sup> | 7.76E-03 | 8.658E-02<sup>c</sup> | 8.06E-03 | 7.58E-2 [6.69E-2,8.48E-2] | 91.44% |
| All Other Variants (10kb buffer model) | 6.607E-02<sup>c</sup> | 7.80E-03 | 7.042E-02<sup>c</sup> | 7.73E-03 | 8.408E-02<sup>c</sup> | 8.03E-03 | 7.33E-2 [6.44E-2,8.22E-2] | 88.00% |
| All Other Variants (25kb buffer model) | 6.359E-02<sup>c</sup> | 7.75E-03 | 6.933E-02<sup>c</sup> | 7.70E-03 | 8.184E-02<sup>c</sup> | 7.99E-03 | 7.14E-2 [6.25E-2,8.02E-2] | 85.71% |
| All Other Variants (35kb buffer model) | 6.319E-02<sup>c</sup> | 7.73E-03 | 6.898E-02<sup>c</sup> | 7.68E-03 | 8.105E-02<sup>c</sup> | 7.97E-03 | 7.09E-2 [6.20E-2,7.97E-2] | 85.22% |
| All Other Variants (50kb buffer model) | 4.216E-02<sup>c</sup> | 7.40E-03 | 4.524E-02<sup>c</sup> | 7.34E-03 | 6.242E-02<sup>c</sup> | 7.72E-03 | 4.96E-2 [4.11E-2,5.80E-2] | 59.12% |
| *Total* |  |  |  |  |  |  |  |  |
| Total heritability (0kb buffer model) | 7.530E-02 | 7.86E-03 | 7.920E-02 | 7.79E-03 | 9.400E-02 | 8.08E-03 | 8.26E-2 [7.36E-2,9.15E-2] | N/A |
| Total heritability (5kb buffer model) | 7.530E-02 | 7.85E-03 | 7.970E-02 | 7.78E-03 | 9.420E-02 | 8.08E-03 | 8.29E-2 [7.39E-2,9.18E-2] | N/A |
| Total heritability (10kb buffer model) | 7.570E-02 | 7.84E-03 | 8.030E-02 | 7.78E-03 | 9.470E-02 | 8.07E-03 | 8.33E-2 [7.44E-2,9.23E-2] | N/A |
| Total heritability (25kb buffer model) | 7.590E-02 | 7.84E-03 | 8.000E-02 | 7.77E-03 | 9.480E-02 | 8.07E-03 | 8.33E-2 [7.44E-2,9.22E-2] | N/A |
| Total heritability (35kb buffer model) | 7.590E-02 | 7.84E-03 | 7.980E-02 | 7.78E-03 | 9.450E-02 | 8.07E-03 | 8.32E-2 [7.43E-2,9.21E-2] | N/A |
| Total heritability (50kb buffer model) | 7.670E-02 | 7.85E-03 | 8.090E-02 | 7.79E-03 | 9.480E-02 | 8.09E-03 | 8.39E-2 [7.49E-2,9.28E-2] | N/A |

Table shows the estimated heritability for each fold and the meta-heritability estimated across folds. Note that components are labelled according to the observed effects used across the models with varied buffer lengths. Consequently, there are no effects for the 0Kb buffer length model. Abbreviations: F1-F3 indicate analysis folds 1 thru 3, N/A - not applicable, SE - standard error. Notations: <sup>a</sup> - $p < 0.05$ , <sup>b</sup> $p < 0.01$ , <sup>c</sup> $p < 0.001$ (van Dam et al., 2019)

**Figure 2.**
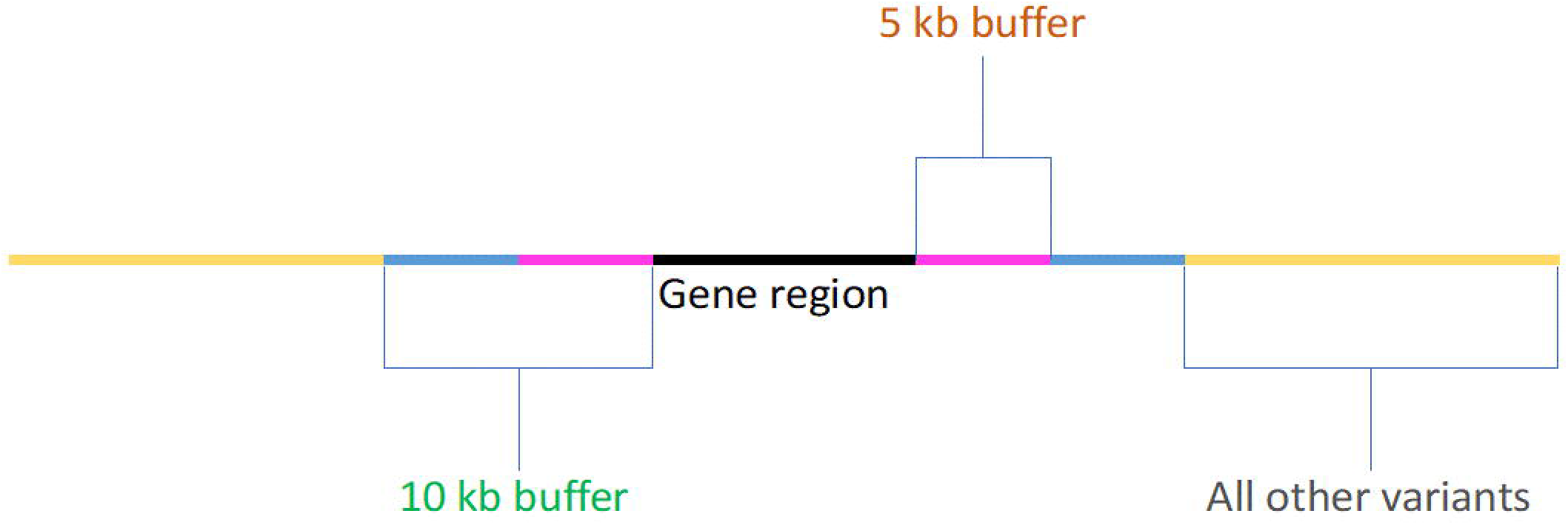
Visualization of each model-component utilized within statistical analyses.

The relative contribution of variants within the gene, buffer, and ‘all other’ components was evaluated under a polygenic model. Regions-of-Interest heritability mapping was achieved using multiple genetic components in GREML analyses implemented in GCTA [version 1.92] using the set of SNPs from each ROI to define the components of the model.^21; 22^ Analyses employed a set of three genetic relatedness matrices (GRMs) for a given fold. Variance component ROI-G reflected variation across SNPs in the transcriptionally regulated gene set depicted in Table 1. ROI-Buffers, of varying lengths, was used to reflect the effect of loci around the ROI-G. ROI-All_Others, reflected aggregate variant effects from the remainder of the genome, given the corresponding size of ROI-G and ROI-Buffer. The significance of each variance component was assessed using a likelihood ratio test while accounting for age and sex. Population stratification effects were controlled using strict selection for individuals of European Ancestry using genomic principal components and multidimensional scaling.^11^ Enrichment (E) values were calculated to determine whether the observed component-heritability estimates were greater than what would be expected by chance given the observed total genetic variance and the 4.6 million SNPs used in the analysis (i.e., the variance explained we would expect via a random selection of loci of the same size from the genome). As such, the statistical significance of an enrichment was evaluated on the basis of whether the expected h^2^_SNP_ fell within the 95% confidence interval of the observed h^2^_SNP_ (i.e., E > 1.96).

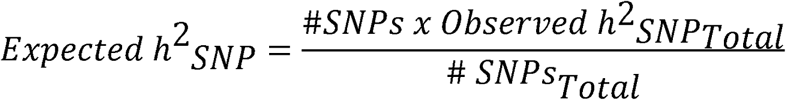

Meta-analyzed SNP-heritability estimates were obtained by pooling results across folds and meta-analyzing using a weighted fixed-effect model. Heritability estimates across UKB-folds were combined using fixed-effects inverse-variance meta-analysis implemented in R using the “rmeta” package. Mixed linear model association analyses were performed in GCTA and gene-based testing were done using MAGMA (version 1.06) implemented in FUMA (v.1.3.5e).^23^ Gene-level p-values were used to conduct gene set tests against “Curated Gene Sets” and “GO terms” pathways identified in Msigdb v5.2.^24^ We considered all SNP and gene-based signals below 5×10^−8^ and 2.89×10^−6^ as genome-wide and gene-wide (i.e., based on 17287 genes tested) significant, respectively; further, we also implemented a less conservative threshold using a False Discovery Rate (FDR) of q<0.05.^25^ All analyses minimized the effects of confounders by including sex, testing site location, age, and age^2^ as covariates.

## RESULTS

### Co-expressed Genes in Model Organisms Explain Variation in Human Tobacco Consumption

The estimated total additive genetic effect (i.e., SNP-heritability) of tobacco consumption ranged from 7.6% to 9.5% across the three folds (see Table 2 reported meta-h^2^ _SNP-Total_ values). Variants across the ROI-genes component of the model (ROI-G) accounted for approximately 0.2-0.4% of the variation in tobacco consumption across folds (see Table 2) while those in the buffer (ROI-Buffer) and remainder of the genome (ROI-All_Others) accounted for 0.4-3% and 5-8%, respectively. There was significant enrichment (E) in almost all instances where the variants in or surrounding the genes of interest were examined (Table 3); no enrichment was observed in the ROI-All_Others category.

**Table 3:**
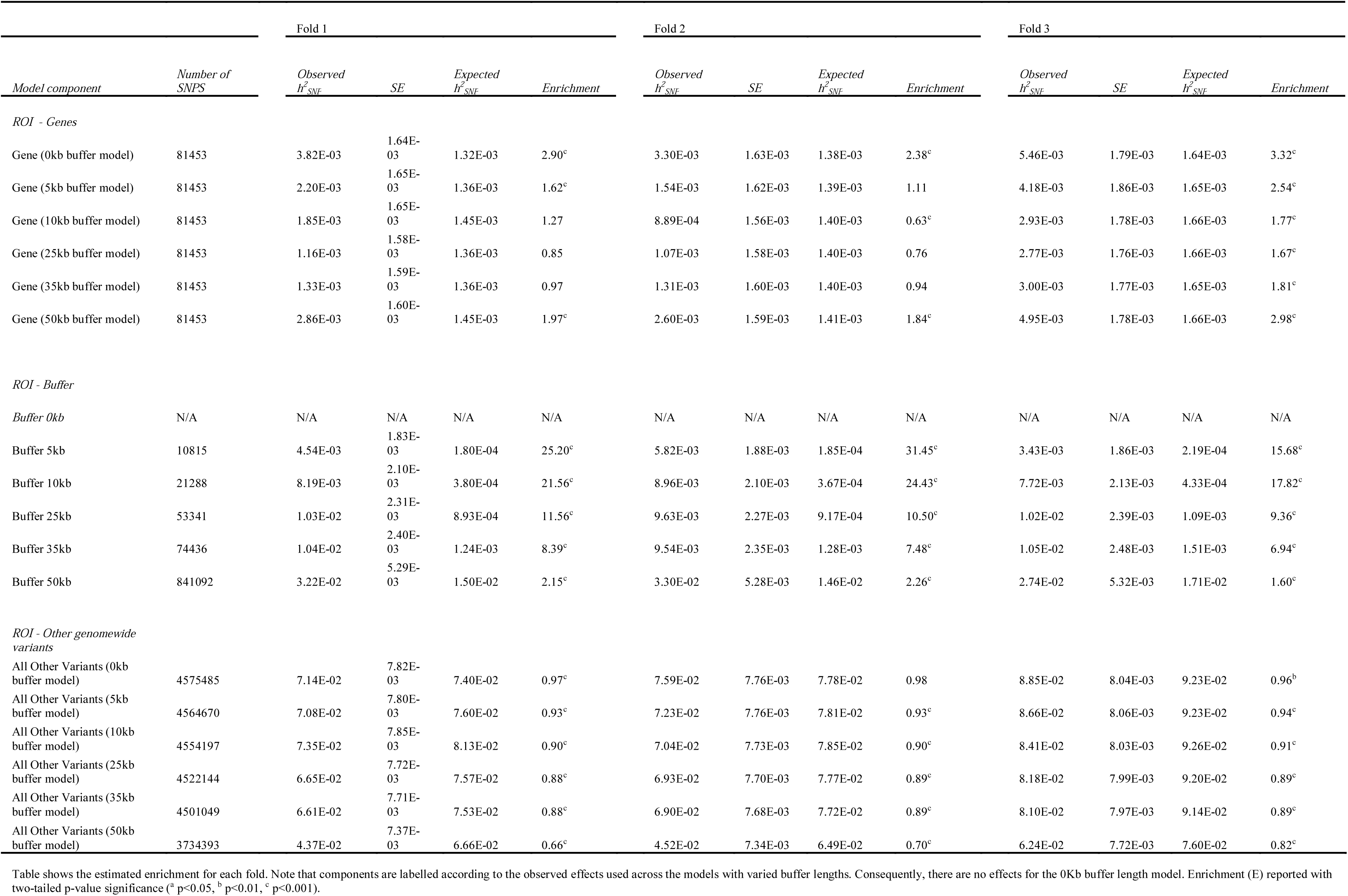
Calculated Enrichment Values for Each Component of the ROI Model.

There was limited association between variance explained by ROI-G and buffer length (model R^2^ across folds ranging 0.003-0.09; see Figure 3 panel A). On the contrary, the variance explained by SNPs in and around the genes of interest (i.e., ROI-Buffer) that were modeled using buffers of various length (ROI-buffer-#Kb) increased over buffer size (model R^2^ across folds ranging 0.75-0.86; see Figure 3 panel B), whereas the variance explained decreased for ROI-All_Others as buffer size increased (model R^2^ across folds ranging 0.73-0.81; see Figure 3 panel C). This result is in line with the observation from previous work that variance explained is proportional to DNA length^22^, consistent with a polygenic model. Notably, variance explained by variants located around genes of interest were positively associated with buffer size, but the enrichment decreased with buffer size (see Figure 4), suggesting that that the trait-associated variants are more enriched near genes.

**Figure 3.**
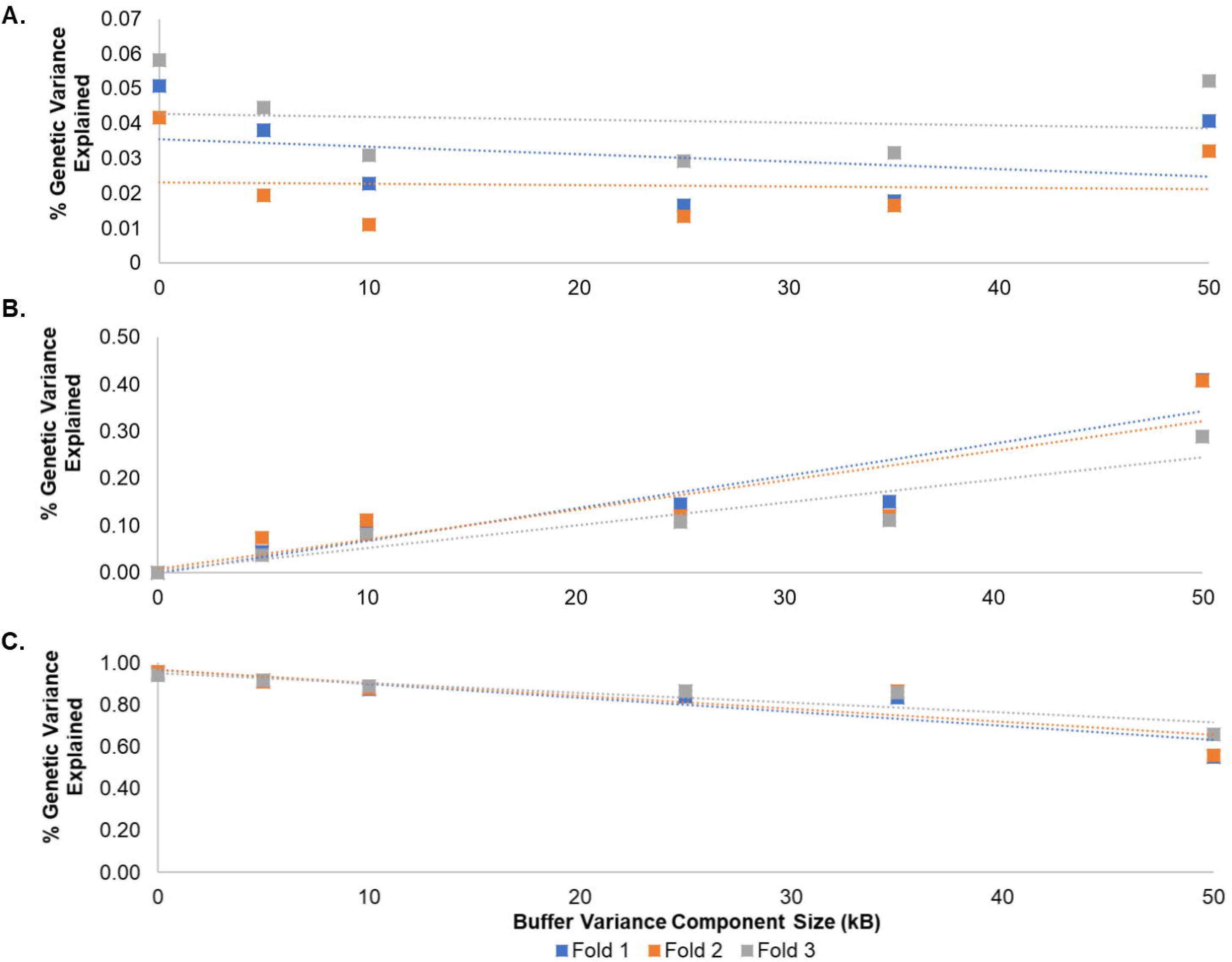
Plots of the relationship between buffer length and percent genetic variance explained by each model component. Citation: Lines shown reflect inferred trends for buffer lengths not assessed. Panel A shows the percent genetic variance explained by the gene region model component. Observed relationships between length and variance explained are reflected by the regression equation and model fit (r-squared; R^2^) by the following equations for Fold 1: - 0.002(Buffer length) + 0.0355 [model R^2^=0.0869]; Fold 2: -4E-05(Buffer length) + 0.0231 [model R^2^=0.0034]; Fold 3: -8E-05(Buffer length) + 0.0428 [model R^2^=0.0869]. Panel B shows the percent genetic variance explained by the buffer region model component. For this component, the observed relationships between length and variance explained are reflected by the regression equation and model fit (r-squared) by the following equations for Fold 1: 0.007(Buffer length) - 0.001 [model R^2^=0.858]; Fold 2: 0.006(Buffer length) + 0.008 [model R^2^=0.754]; Fold 3: 0.005(Buffer length) + 0.004 [model R^2^=0.86]. Lastly, Panel C describes the percent genetic variance explained by the “all-other variants” model component. The observed relationships between length and variance explained for panel C are reflected by the regression equation and model fit (r-squared) by the following equations for Fold 1: -0.007(Buffer length) + 0.966 [model R^2^=0.809]; Fold 2: -0.006(Buffer length) + 0.969 [model R^2^=0.729]; Fold 3: - 0.005(Buffer length) + 0.953 [model R^2^=0.812].

**Figure 4.**
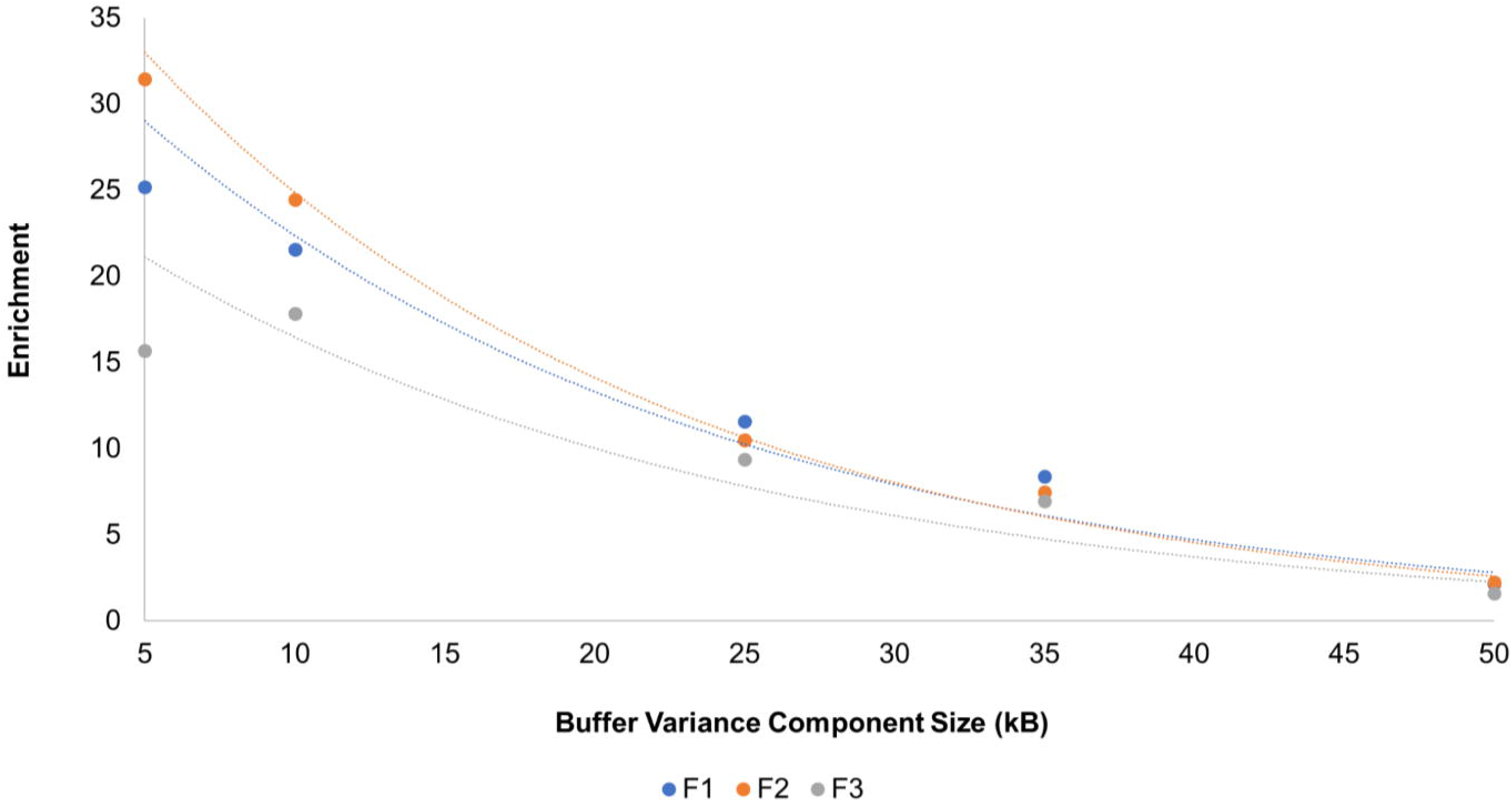
Scatterplot illustrating change in enrichment (E) of the set of ROI-buffer variants as a function of models of with varied buffer size. Abbreviations F1, F2, and F3, correspond to folds 1, 2, and 3, respectively.

### Genome-wide Association, Gene-based, and Gene set effects

Association analyses using all 139,043 smokers confirmed previously associated regions identified in a larger meta-analysis that included these data.^6^ We identified 594 signals that were genomewide significant, and a larger set of 938 signals with q<0.05 (see supplementary Table S3 for complete summary statistics and Supplementary Figures S2 and S3 for the Manhattan and Q-Q plot, respectively). The top signals resided on chromosomes 15, 19, 8, 7, 4, 3, and 1 (see Supplementary Figures S4 thru S7 for regional association plots for associations across nicotinic acetylcholine receptor genes CHRNA4/A5/A6 and CYP2A6, respectively). Most of the associated SNPs are functionally annotated as intronic, intergenic, and intronic non-coding RNA (see Supplementary Figure S8). Gene-based analyses identified 20 genes that surpassed the Bonferroni significance threshold and 31 with q<0.05 (see Supplementary Table S4 and Supplementary Figures S9 and S10 for the gene-based test Manhattan and Q-Q plots, respectively). Of the gene-wide significant genes, four were differentially expressed across the model organism experiments and this overlap was more than we would expect by chance, OR = 7.20, empirical *p* = 4.41E-3. Post-hoc examination of the test statistics (i.e., using 10,000 permutations of 500 gene sets from non-GeneWeaver genes) indicated that the majority of the signals originated from genes largely captured by the *a priori Mus musculus* studies (two sample t-test: t = 2.2813, df= 664.87, empirical p = 0.023; Supplementary Figure S11). Gene set analyses, which focused on curated gene sets and GO term annotations from MsigDB, identified 745 significant gene sets (p<0.05), but only one gene set, REACTOME: Presynaptic Nicotinic Acetylcholine Receptors (R-HSA-622323; https://reactome.org/PathwayBrowser/#/R-HSA-622323) survived multiple-testing correction (Bonferroni-corrected p = 2.5×10^−8^).

## DISCUSSION

We integrated genomic and bioinformatic analyses which provided a rapid approach for filling the translational space between human and animal genetics research. Similar to other genetic studies of drug use^26-28^, these findings indicated a neuro-epigenetic component to the genetic inheritance of tobacco consumption, while also localizing genomic regions of interest. By using a genetic sample of over 100,000 humans and meta-analyzing across three species from seven gene expression studies, we found that approximately 4.2%-39.5% of the heritability for the frequency of human tobacco use can be attributed to mRNA readout related to nicotine exposure/consumption in the brain. Given that the observed neuro-molecular associations observed with tobacco/nicotine use were inferred via model systems, irrespective of prior GWAS findings, it stands to reason that integrating knowledge across species will enhance genomic discoveries related to tobacco use. Importantly, most cross-species findings appeared to be buried under the conservative genome-wide significant threshold – demonstrating the strength of our approach, which incorporates significant and non-significant sources of genomic variations *a priori* and helps accommodate the numerous (relevant) genes with small effect sizes riddled across the human genome. Notably, these observations highlight an interesting perspective of polygenic effects, in so much as it provides support for a mixture of effects on tobacco consumption.

This study demonstrates the importance of transcriptionally regulated genes and is in accordance with broad human GWAS research, which detects most of its associations among intergenic regions.^1^ Our results suggest that the genetic proclivity to tobacco use is mediated, in part, by gene expression in relevant brain regions that relate to specific behavioral mechanisms. Similarly recent genome-wide research identified genome-wide significant loci in neurotransmission and reward learning genes for tobacco use and prioritized non-synonymous protein coding variants.^6^ By using just half of the sample size from Liu et al., our findings corroborated the importance of reward-related and neurotransmission genes and further disentangle the underlying genetic structure of tobacco consumption by highlighting transcriptionally relevant cis-eQTLs in hundreds of genomic regions. Overall, our study suggests that the genetic architecture of tobacco consumption feeds into the neuro-molecular landscape via modulation of gene expression.

These data also suggest that the use of model systems allows for the direct sampling of brain tissue, in the context of a trait relevant phenotype which models, in a simplified way, characteristics of human disease measured in an organism (e.g., *Mus musculus, Drosophilae Melanogaster, Rattus Norvegicus*, and *Caenorhabditis elegans*, to name a few) with a genome that has some similarities to humans, including mammals with high percentages of orthologous genes. It is important to note that when we refer to “modeling” here, we are not referring to the questionable practice of establishing a single gene perturbation in a model organism as a “model” of a person with a disease. Rather we are referring to the practice of evaluating the complex genomic basis of traits that are characteristic of various aspects of the disease state. Taken alone, model system work has a number of key advantages (over and above access to brain tissue) including, but not limited to, the use of neurogenetic methods (e.g., optogenetic, thermogenetic, etc.) which can introduce much larger biological effects in model systems than could be seen in typical GWAS studies. Additionally, controlled environmental exposures (e.g., pharmacological, behavioral, etc.) may be used in model systems in a fashion that would be impossible in humans. The strengths of model systems allow for smaller-sample studies to be maximally informative due to larger effect sizes and tighter experimental control, but the “translatability” of these findings to the human condition has limitations. While some more basic behavioral traits are convincingly modeled in animals, other complex phenotypes and disorders are represented only in part by these systems ^29^. Furthermore, the phylogenetic distance between the model organism and *Homo sapiens* can pose additional challenges as only a subset of genes will be conserved in an informative way; notably, studies have shown conservation of epigenetic marks across mice and humans.^30^ Attempts to leverage conserved evidence across mice and humans in alcohol dependence research have revealed networks of genes and loci, which had gone undetected in prior GWAS.^31^ In sum, model systems bring unique advantages and disadvantages to behavior genetics that may complement human GWAS studies of related traits.

These analyses identified various genes previously linked to nicotine consumption and cessation, including validated nicotinic acetylcholine receptor genes *CHRNA3/A4/A5/B4*, as well as nicotine metabolism genes (*CYP2A6/A7*), which provides a sanity-check for our genome-wide analyses. Mechanistic research in mice suggests that a mutation of the *CHRNA5* gene and concomitant habenular expression of *CHRNA5* robustly increases nicotine consumption, but not after experimentally restoring habenuala *CHRNA5* levels back to normal^32^. These results buttress our findings delineating the path from genetic predisposition to gene expression and eventually specific behavioral outcomes and may suggest a gene x drug interaction. That is, those at higher genetic risk for tobacco use may have an altered physiological response that increases susceptibility for augmented consumption.^33; 34^ Apart from the established nicotinic acetylcholine receptors, we also discovered significant genetic association of chromosome 19 genes: *RAB4B, EGLN2* and *CYP2A6* with tobacco consumption. *RAB4B* is involved in the breakdown of GTP for vesicular transport^35^ and was previously associated with PFC gene expression among those with major depression.^36^ While *RAB4B, EGLN2* and *CYP2A6* are in strong linkage disequilibrium, research suggests they correspond to largely independent mechanisms.^37^ Our study suggests that *RAB4B* is driven by a brain-dependent mechanism (identified in mice)^38^ and might underlie neuroplasticity processes related to nicotine reward^39^. On the other hand, *EGLN2* and *CYP2A6* were not associated with gene expression findings in animal models of nicotine use/exposure. *EGLN2* is a hypoxia inducible factor and plays a role in oxygen homeostasis^40^ and may be uniquely associated with humans because the vehicle for nicotine intake is via oxygen restricting smoke (i.e., carbon monoxide present in cigarette smoke preferentially binds to hemoglobin and thus reduces its ability to transport oxygen), whereas animal models typically study nicotine through injections or implementation in the drinking water. *CYP2A6* is an enzyme that accounts for ∼80% of nicotine clearance^41^ and is almost exclusively expressed in the liver, which is a likely reason that it was not included in our brain-mediated cross-species gene list. Therefore, our integrative approach better contextualizes the effects of genes associated with human complex traits and better determines how specific genetic associations relate to relevant model systems in particular tissues.

While novel, there are several considerations for interpreting the current findings. First, these analyses are limited by current understanding of the consequences of tobacco exposure using only microarray studies. We sought to overcome this limitation by integrating multiple sources of information using differences across brain and model organisms, but future studies are needed to determine whether these effects are invariant, as well as whether the experimental paradigm itself may alter this line of evidence, especially as the volume of literature increases. Second, our analyses did not examine genes that have been shown to be differentially methylated by tobacco exposure; we assumed that such processes would equate to direct differences in mRNA levels, constructed gene list utilized animal research, which focused primarily on orthologous genes.^42; 43^ As such, there was less emphasis on regulatory elements for said genes, which may also generalize across species. We attempted to capture said effects by using buffers of various lengths to approximate the relative import of *cis* and possibly *trans* acting effects. It should be noted however, that our results are in line with the *multiple enhancer variant hypothesis*, which purports a similar role of noncoding variants in common traits.^44^

Future research is warranted to determine whether our integrative framework generalizes across complex human traits. Traits with different genetic architectures, epigenetic landscapes and animal models may yield disparate findings. We found that the bulk of our cross-species signal stemmed from mouse models of nicotine use, but it will be important for future research to be conducted across multiple smoking phenotypes and include additional species/studies and incorporate findings from human tissues to benchmark findings with other model organisms. Ideally, integrative genomics comparisons would leverage equitable and minimally error prone outcomes or endophenotypes across studies. Given the array of animal models for human traits, an inviting avenue of research should clarify the utility of specific tissues, cell types and animal models in human genetics. With a large enough literature base, we may be able to better refine what tissues and specific mechanisms human genomic signals stem from and ultimately may better characterize the genetic make-up for complex traits. Future studies leveraging these approaches should consider strategies for reducing buffer size and examining heterogeneity across tissue/cell types, as well as whether the observed effects generalize across human populations (e.g., European, African, Asian, etc).

## Conclusions

In sum, this study represents a step forward for interspecies behavioral genetics and provides a proof of principle for bridging the gap between human and animal genetics in identifying polygenic risk variants. We show that enhancing human GWAS by incorporating *a priori* information on relevant traits (even across species) is a worthwhile path to unraveling the genetic basis for complex traits.

## Supporting information

supplementary collection

## Acknowledgements

We acknowledge the National Institute on Drug Abuse award DP1DA042103 (to RHCP) and the National Institute on Alcohol Abuse and Alcohol (R01AA018776) (to EJC). We acknowledge the Wellcome Trust medical charity, Medical Research Council, Department of Health, Scottish Government, Northwest Regional Development Agency, Welsh Government, British Heart Foundation, Cancer Research UK and Diabetes UK, and the National Health Service (NHS) for their part in supporting the UK Biobank without which this study would not have been possible. The contents of this paper do not represent the views of the U.S. Department of Veterans Affairs or the United States Government.

## Data Accessibility Information

The genetic and phenotype datasets from UK Biobank that were analyzed here are available via the UK Biobank data access process (see http://www.ukbiobank.ac.uk/register-apply/). Detailed information about the genetic data available from UK Biobank is available at http://www.ukbiobank.ac.uk/scientists-3/genetic-data/ and http://biobank.ctsu.ox.ac.uk/crystal/label.cgi?id=100314. Note that the exact number of samples with genetic data currently available in UK Biobank may differ slightly from those described in this paper as it is subject to the data use agreement at the time of each study.

