## supplementary collection for "Cross-Species Integration of Transcriptomic Effects of Tobacco and Nicotine Exposure Helps to Prioritize Genetic Effects on Human Tobacco Consumption"

### Supplemental Text

*Gene Set Extraction Protocol using GeneWeaver*

Once a list of outputs were generated from a search, the abstract of each gene set would be read in order to determine if the study followed a pre-set inclusion criteria. The inclusion criteria are as followed: (1) the study had to utilize a phenotype of nicotine consumption / exposure as its dependent variable, (2) the gene list extracted from the study had to have been the direct result of experimentation, (3) the study itself analyzed expression-Quantitative Trait Loci (eQTL), therefore consisted of a microarray or WGCNA design, and was not, for example, a GWAS. If a gene set was found to be relevant for subsequent analysis, then it was added to its appropriate project.

*Data-Screening and Gene list Extraction*

Once data-mining was completed, studies were double-checked to make sure they met inclusion criteria. For every gene set, its PubMed ID (provided in the gene set description) was used to download its publication, which was then read over in full to make sure the study design represented a dependent variable that would be relevant for subsequent analysis. Once passing this screening process, gene sets were then combined if they all belonged to the same study, only differing in neural region or time. This was done utilizing GeneWeaver's "Boolean gene set logic" tool, which takes several different gene sets and outputs a single gene set containing the union, intersection, or highest-replicated genes through all the studies. In the case of this study, gene sets that belonged to the same publication were submitted through the tool's "Union" function, a tool which allows a single export of all genes present within all inputted gene sets. "Homology" (a feature provided by GeneWeaver to prevent duplicating genes that differentiate only by species) selected to be "On"[18]. Upon the completion of data-screening, the "Boolean gene set tool" was used to create a final gene list that would be used for analysis.

**List of Supplementary Tables and Figures (with citations)**

Table S1: Jaccard Similarity estimates across identified gene sets

Table S2: Replicated genes across Gene Weaver Studies

Table S3: List of SNPs from MLMA analysis that reach genome-wide significance

Table S4: Top genes identified after correction for multiple testing

Figure S1: Gene set Graph showing representation of genes across gene sets identified in Table 1 from five studies. Citation: Plots of the relationship between buffer length and the calculated enrichment values explained by the buffer-length model components. Enrichment values are calculated across all 3 folds to be representative of the entire sample, not by-fold. Lines shown reflect inferred trends for buffer lengths not assessed.

Figure S2: Manhattan plot of tobacco consumption GWAS

Figure S3:  GWAS-summary QQ plot visualization based on the mixed-linear model-based association

Figure S4: GWAS regional association plot of CHRNA4 on chromosome 20

Figure S5: GWAS regional association plot of CHRNA5 on chromosome 15

Figure S6: GWAS regional association plot of CHRNA6 on chromosome 8

Figure S7: GWAS regional association plot of CYP2A6 on chromosome 19

Figure S8: GWAS SNP Annotation plot

Figure S9: Gene-based Manhattan plot

Figure S10: Gene-based QQ plot visualization based on MAGMA gene testing

Figure S11. Cross−Species comparison of human gene−based test on tobacco use. Citation: Comparison of the –log10 p-value across contributing gene sets was conducted using Welch Two Sample t-tests (Mus musculus:Danio rerio- t=0.9244, df=225.54, p=0.3563; Mus musculus:Rattus norvegicus-t=2.3719, df=833.63, p=0.01792). Note we re-sampled the average –log10 p-value (10,000 permutations) of a list of 500 genes outside of our Gene Weaver list (Non-Gene Weaver Genes) to determine whether the genes from our nicotine Gene Weaver lists were more significant in our human gene-based test than we would expect by chance. Only the gene list from mice was significantly greater than this random permuted list of non-Gene Weaver genes (Mus musculus:Non-Gene Weaver Genes- $t=2.4342$, $df=640.03$, $p=0.0152$).

Table S1. Jaccard Similarity Estimates Across Identified gene sets

| Gene set | Jaccard Similarity Estimate | | | | |
|---|---|---|---|---|---|
| Merged Kily et al.[30] | 1 | | | | |
| Merged Wang et al.[29] | 6.00E-03 | 1 | | | |
| Chen et al.[27] | 0.00E+00 | **1.59E-02** | 1 | | |
| Sharp et al.[31] | 3.30E-03 | 3.50E-03 | 5.40E-03 | 1 | |
| Polesskaya et al.[28] | 5.50E-03 | **8.80E-03** | **8.10E-03** | 0.00E+00 | 1 |

Note: Gene sets shown do not exactly align with the full complement of GeneWeaver set IDs shown in Table 1 as several sets (within studies) were merged to reduce overrepresentation of genes in a given study. Bolded values indicate $p < 0.05$.

Table S2. Replicated Genes Across Gene Weaver Studies

| GENE_NAME | STUDY_1 | SPECIES_1 | BEHAVIOR_1 | BRAIN_REGION_1 | STUDY_2 | SPECIES_2 | BEHAVIOR_2 | BRAIN_REGION_2 |
|---|---|---|---|---|---|---|---|---|
| ACTB | Wang | mouse | Nic_Self-Admin | Amyg_Hipp_nAcc_PFC_VTA | Chen | mouse | Acute_Nic | VTA |
| ALDH7A1 | Sharp | rat | Chronic_Nic_Self-Admin | nAcc | Kily | fish | Nic_CPP | Whole_Brain |
| CCT3 | Wang | mouse | Nic_Self-Admin | Amyg_Hipp_nAcc_PFC_VTA | Polesskaya | rat | Chronic_Nic_Treatment | PFC_vStriatum_Hipp |
| DSTN | Wang | mouse | Nic_Self-Admin | Amyg_Hipp_nAcc_PFC_VTA | Sharp | rat | Chronic_Nic_Self-Admin | nAcc |
| GABBR1 | Wang | mouse | Nic_Self-Admin | Amyg_Hipp_nAcc_PFC_VTA | Chen | mouse | Acute_Nic | VTA |
| ITPR2 | Wang | mouse | Nic_Self-Admin | Amyg_Hipp_nAcc_PFC_VTA | Chen | mouse | Acute_Nic | VTA |
| KCNK1 | Wang | mouse | Nic_Self-Admin | Amyg_Hipp_nAcc_PFC_VTA | Polesskaya | rat | Chronic_Nic_Treatment | PFC_vStriatum_Hipp |
| KRT5 | Kily | fish | Nic_CPP | Whole_Brain | Polesskaya | rat | Chronic_Nic_Treatment | PFC_vStriatum_Hipp |
| MOBP | Wang | mouse | Nic_Self-Admin | Amyg_Hipp_nAcc_PFC_VTA | Chen | mouse | Acute_Nic | VTA |
| PPM1B | Kily | fish | Nic_CPP | Whole_Brain | Chen | mouse | Acute_Nic | VTA |
| PRKACB | Chen | mouse | Acute_Nic | VTA | Polesskaya | rat | Chronic_Nic_Treatment | PFC_vStriatum_Hipp |
| PSEN1 | Wang | mouse | Nic_Self-Admin | Amyg_Hipp_nAcc_PFC_VTA | Polesskaya | rat | Chronic_Nic_Treatment | PFC_vStriatum_Hipp |
| PSMB5 | Wang | mouse | Nic_Self-Admin | Amyg_Hipp_nAcc_PFC_VTA | Chen | mouse | Acute_Nic | VTA |
| PSMD7 | Wang | mouse | Nic_Self-Admin | Amyg_Hipp_nAcc_PFC_VTA | Sharp | rat | Chronic_Nic_Self-Admin | nAcc |
| RAD21 | Sharp | rat | Chronic_Nic_Self-Admin | nAcc | Chen | mouse | Acute_Nic | VTA |

| | | | | | | | | |
|---|---|---|---|---|---|---|---|---|
| RPS28 | Wang | mouse | Nic_Self-Admin | Amyg_Hipp_nAcc_PFC_VTA | Sharp | rat | Chronic_Nic_Self-Admin | nAcc |
| SST | Wang | mouse | Nic_Self-Admin | Amyg_Hipp_nAcc_PFC_VTA | Chen | mouse | Acute_Nic | VTA |
| SYT4 | Wang | mouse | Nic_Self-Admin | Amyg_Hipp_nAcc_PFC_VTA | Chen | mouse | Acute_Nic | VTA |
| TFDP1 | Wang | mouse | Nic_Self-Admin | Amyg_Hipp_nAcc_PFC_VTA | Kily | fish | Nic_CPP | Whole_Brain |
| UCHL1 | Wang | mouse | Nic_Self-Admin | Amyg_Hipp_nAcc_PFC_VTA | Polesskaya | rat | Chronic_Nic_Treatment | PFC_vStriatum_Hipp |
| YWHAG | Wang | mouse | Nic_Self-Admin | Amyg_Hipp_nAcc_PFC_VTA | Chen | mouse | Acute_Nic | VTA |

Table showing the two studies (study 1 and study 2) in GeneWeaver for which a gene was observed to be differentially expressed across two brain tissues (brain region 1/2) for varying smoking/nicotine exposure experimental paradigms.

61

**Table S3. List of SNPs from MLMA Analysis That Reach Genome-wide Significance**

| Chr | SNP | bp | A1 | A2 | Freq | b | se | p | qvalue |
|---|---|---|---|---|---|---|---|---|---|
| 15 | rs72738786 | 78828086 | T | G | 0.336253 | 0.982504 | 0.050458 | 1.90E-84 | 1.90E-78 |
| 15 | rs8192482 | 78886198 | T | C | 0.332725 | 0.987857 | 0.050697 | 1.46E-84 | 1.90E-78 |
| 15 | rs4887067 | 78886947 | A | G | 0.332728 | 0.987736 | 0.050698 | 1.53E-84 | 1.90E-78 |
| 15 | rs55676755 | 78898932 | G | C | 0.33346 | 0.986251 | 0.050671 | 2.23E-84 | 1.90E-78 |
| 15 | rs147144681 | 78900908 | T | C | 0.333394 | 0.986231 | 0.050676 | 2.33E-84 | 1.90E-78 |
| 15 | rs146009840 | 78906177 | T | A | 0.333046 | 0.986519 | 0.050698 | 2.45E-84 | 1.90E-78 |
| 15 | rs72740955 | 78849779 | T | C | 0.335669 | 0.983358 | 0.050626 | 4.85E-84 | 3.22E-78 |
| 15 | rs8031948 | 78816057 | T | G | 0.336476 | 0.979386 | 0.050463 | 6.60E-84 | 3.36E-78 |
| 15 | rs11633958 | 78862064 | T | C | 0.331731 | 0.986843 | 0.050859 | 7.21E-84 | 3.36E-78 |
| 15 | rs113931022 | 78901113 | T | C | 0.333531 | 0.9834 | 0.050668 | 6.53E-84 | 3.36E-78 |
| 15 | rs34684276 | 78813155 | A | G | 0.335919 | 0.980237 | 0.050537 | 8.30E-84 | 3.51E-78 |
| 15 | rs111704647 | 78900650 | T | C | 0.333355 | 0.982324 | 0.050678 | 1.06E-83 | 4.11E-78 |
| 15 | rs2036527 | 78851615 | A | G | 0.335873 | 0.979389 | 0.050613 | 2.02E-83 | 6.73E-78 |
| 15 | rs7172118 | 78862453 | A | C | 0.331989 | 0.983948 | 0.050839 | 1.88E-83 | 6.73E-78 |
| 15 | rs138544659 | 78900701 | G | T | 0.333041 | 0.980841 | 0.050698 | 2.17E-83 | 6.74E-78 |
| 15 | rs17486195 | 78865197 | G | A | 0.332463 | 0.98016 | 0.05073 | 3.58E-83 | 9.38E-78 |
| 15 | rs140330585 | 78866445 | A | G | 0.332367 | 0.980137 | 0.050723 | 3.41E-83 | 9.38E-78 |
| 15 | rs56390833 | 78877381 | A | C | 0.332351 | 0.980016 | 0.050725 | 3.62E-83 | 9.38E-78 |
| 15 | rs8042849 | 78817929 | C | T | 0.344214 | 0.964855 | 0.050149 | 1.72E-82 | 4.21E-77 |
| 15 | rs112878080 | 78900647 | G | A | 0.332248 | 0.975031 | 0.050755 | 3.01E-82 | 7.02E-77 |
| 15 | rs8039449 | 78914534 | T | C | 0.36557 | 0.892218 | 0.049237 | 2.18E-73 | 4.84E-68 |
| 15 | rs17484524 | 78772676 | G | A | 0.327241 | 0.855307 | 0.050225 | 4.97E-65 | 1.05E-59 |
| 15 | rs72738732 | 78752188 | G | C | 0.327206 | 0.854901 | 0.050253 | 6.70E-65 | 1.36E-59 |
| 15 | rs17484235 | 78761414 | G | C | 0.327277 | 0.854048 | 0.050228 | 7.76E-65 | 1.51E-59 |
| 15 | rs2089162 | 78739763 | G | A | 0.327022 | 0.85403 | 0.050257 | 9.23E-65 | 1.72E-59 |
| 15 | rs17483548 | 78730313 | A | G | 0.32671 | 0.854108 | 0.050292 | 1.10E-64 | 1.97E-59 |
| 15 | rs17405217 | 78731149 | T | C | 0.326723 | 0.853555 | 0.050293 | 1.33E-64 | 2.29E-59 |
| 15 | rs4887056 | 78734585 | G | A | 0.32995 | 0.847055 | 0.050162 | 5.67E-64 | 8.97E-59 |
| 15 | rs56219465 | 78742579 | G | A | 0.330694 | 0.845923 | 0.050098 | 5.78E-64 | 8.97E-59 |
| 15 | rs72738736 | 78765122 | T | G | 0.330625 | 0.845819 | 0.050091 | 5.74E-64 | 8.97E-59 |

| | | | | | | | | | |
|---|---|---|---|---|---|---|---|---|---|
| 15 | rs17483929 | 78742376 | A | G | 0.330699 | 0.845658 | 0.050098 | 6.30E-64 | 9.47E-59 |
| 15 | rs7181486 | 78741618 | C | T | 0.330656 | 0.844752 | 0.050099 | 8.62E-64 | 1.25E-58 |
| 15 | rs17483686 | 78733390 | T | A | 0.330273 | 0.845135 | 0.050137 | 9.39E-64 | 1.33E-58 |
| 15 | rs55983731 | 78735269 | T | C | 0.330296 | 0.844721 | 0.050137 | 1.08E-63 | 1.48E-58 |
| 15 | rs72738718 | 78735438 | C | G | 0.330296 | 0.84443 | 0.050136 | 1.18E-63 | 1.57E-58 |
| 15 | rs2938670 | 78740688 | G | T | 0.331175 | 0.84292 | 0.05007 | 1.35E-63 | 1.68E-58 |
| 15 | rs2656052 | 78740932 | C | A | 0.331147 | 0.842921 | 0.050072 | 1.37E-63 | 1.68E-58 |
| 15 | rs2568494 | 78740964 | A | G | 0.331147 | 0.842921 | 0.050072 | 1.37E-63 | 1.68E-58 |
| 15 | rs1504550 | 78766250 | G | A | 0.325754 | 0.846061 | 0.050289 | 1.62E-63 | 1.94E-58 |
| 15 | rs17483721 | 78733731 | C | T | 0.330465 | 0.843673 | 0.050157 | 1.72E-63 | 2.01E-58 |
| 15 | rs2656065 | 78750549 | A | G | 0.331536 | 0.840572 | 0.050061 | 2.84E-63 | 3.23E-58 |
| 15 | rs2009746 | 78754102 | G | A | 0.331771 | 0.837775 | 0.05005 | 6.85E-63 | 7.60E-58 |
| 15 | rs3825845 | 78910258 | T | C | 0.218626 | -0.85659 | 0.056945 | 3.88E-51 | 4.21E-46 |
| 15 | rs8043009 | 78908154 | C | G | 0.223402 | -0.84393 | 0.056696 | 4.11E-50 | 4.35E-45 |
| 15 | rs8042059 | 78907859 | C | A | 0.223277 | -0.84095 | 0.056709 | 9.49E-50 | 9.60E-45 |
| 15 | rs8042374 | 78908032 | G | A | 0.223277 | -0.84095 | 0.056709 | 9.49E-50 | 9.60E-45 |
| 15 | rs7177514 | 78907406 | G | C | 0.223261 | -0.84056 | 0.056709 | 1.05E-49 | 1.04E-44 |
| 15 | rs7171869 | 78900909 | A | G | 0.223173 | -0.83999 | 0.056745 | 1.40E-49 | 1.36E-44 |
| 15 | rs11637630 | 78899719 | G | A | 0.223159 | -0.83861 | 0.056735 | 1.94E-49 | 1.84E-44 |
| 15 | rs7183604 | 78899213 | T | C | 0.223156 | -0.83754 | 0.056733 | 2.54E-49 | 2.36E-44 |
| 15 | rs189218934 | 78903987 | T | C | 0.222971 | -0.83778 | 0.056754 | 2.59E-49 | 2.36E-44 |
| 15 | rs8042494 | 78908010 | T | C | 0.223156 | -0.8364 | 0.056719 | 3.24E-49 | 2.90E-44 |
| 15 | rs7170068 | 78912943 | A | G | 0.212196 | -0.84545 | 0.057426 | 4.62E-49 | 4.06E-44 |
| 15 | rs13329271 | 78914230 | C | A | 0.212194 | -0.84399 | 0.057458 | 7.61E-49 | 6.56E-44 |
| 15 | rs4887069 | 78909070 | G | A | 0.226632 | -0.82996 | 0.056533 | 8.52E-49 | 7.21E-44 |
| 15 | rs6495309 | 78915245 | T | C | 0.211199 | -0.84316 | 0.057612 | 1.68E-48 | 1.40E-43 |
| 15 | rs7359276 | 78892661 | C | T | 0.225044 | -0.82206 | 0.05653 | 6.55E-48 | 5.35E-43 |
| 15 | rs3743078 | 78894759 | C | G | 0.2253 | -0.8212 | 0.05652 | 7.91E-48 | 6.35E-43 |
| 15 | rs576982 | 78870803 | T | C | 0.223043 | -0.81646 | 0.0567 | 5.21E-47 | 4.11E-42 |
| 15 | rs637137 | 78873976 | A | T | 0.222994 | -0.81616 | 0.056708 | 5.78E-47 | 4.49E-42 |
| 15 | rs28669908 | 78910267 | A | C | 0.208149 | -0.82903 | 0.057944 | 1.96E-46 | 1.50E-41 |
| 15 | rs2456020 | 78868398 | T | C | 0.22454 | -0.80092 | 0.056544 | 1.52E-45 | 1.14E-40 |

| | | | | | | | | | |
|---|---|---|---|---|---|---|---|---|---|
| 15 | rs28681284 | 78908565 | T | C | 0.212697 | -0.80492 | 0.057667 | 2.81E-44 | 2.08E-39 |
| 15 | rs1711731 | 78861918 | A | G | 0.213178 | -0.80042 | 0.057659 | 8.14E-44 | 5.93E-39 |
| 15 | rs1700006 | 78875623 | G | A | 0.213109 | -0.79714 | 0.057625 | 1.61E-43 | 1.15E-38 |
| 15 | rs8192477 | 78910463 | C | G | 0.276382 | -0.70284 | 0.05144 | 1.68E-42 | 1.19E-37 |
| 15 | rs518425 | 78883813 | G | A | 0.282299 | -0.69669 | 0.051216 | 3.85E-42 | 2.67E-37 |
| 19 | rs12461964 | 41341229 | G | A | 0.494453 | 0.609891 | 0.044853 | 4.13E-42 | 2.83E-37 |
| 15 | rs951984 | 78720915 | A | T | 0.279489 | 0.694951 | 0.051334 | 9.34E-42 | 6.30E-37 |
| 15 | rs564585 | 78886227 | G | A | 0.275827 | -0.69324 | 0.051674 | 4.89E-41 | 3.26E-36 |
| 15 | rs905739 | 78845110 | G | A | 0.215137 | -0.76258 | 0.057404 | 2.85E-40 | 1.87E-35 |
| 15 | rs113352275 | 78840567 | T | C | 0.205916 | -0.76256 | 0.058224 | 3.42E-39 | 2.21E-34 |
| 15 | rs684513 | 78858400 | G | C | 0.207261 | -0.76032 | 0.058227 | 5.73E-39 | 3.66E-34 |
| 15 | rs11072763 | 78724256 | A | G | 0.217739 | -0.70965 | 0.057029 | 1.51E-35 | 9.52E-31 |
| 15 | rs2938671 | 78732754 | A | G | 0.21396 | -0.7142 | 0.057439 | 1.71E-35 | 1.06E-30 |
| 15 | rs11072766 | 78771546 | T | C | 0.214452 | -0.71258 | 0.057345 | 1.88E-35 | 1.15E-30 |
| 15 | rs2656073 | 78742276 | T | G | 0.213978 | -0.7122 | 0.057392 | 2.32E-35 | 1.38E-30 |
| 15 | rs2958719 | 78743029 | G | A | 0.213982 | -0.71206 | 0.057392 | 2.40E-35 | 1.38E-30 |
| 15 | rs2656072 | 78744292 | A | G | 0.213982 | -0.71206 | 0.057392 | 2.40E-35 | 1.38E-30 |
| 15 | rs2656069 | 78745707 | C | T | 0.214008 | -0.71202 | 0.05739 | 2.40E-35 | 1.38E-30 |
| 15 | rs35031105 | 78772806 | T | C | 0.214738 | -0.71118 | 0.057326 | 2.43E-35 | 1.38E-30 |
| 15 | rs28511883 | 78783683 | T | C | 0.21471 | -0.71132 | 0.057336 | 2.42E-35 | 1.38E-30 |
| 15 | rs2036533 | 78781687 | A | G | 0.214657 | -0.71112 | 0.057339 | 2.55E-35 | 1.43E-30 |
| 15 | rs12101809 | 78779801 | T | C | 0.214594 | -0.71115 | 0.057351 | 2.61E-35 | 1.45E-30 |
| 15 | rs28602670 | 78768167 | G | C | 0.214385 | -0.711 | 0.057346 | 2.67E-35 | 1.46E-30 |
| 15 | rs2568493 | 78740233 | G | A | 0.214108 | -0.71128 | 0.057378 | 2.74E-35 | 1.48E-30 |
| 15 | rs2568499 | 78722359 | T | C | 0.213305 | -0.7132 | 0.057545 | 2.82E-35 | 1.51E-30 |
| 15 | rs2568485 | 78752114 | C | T | 0.214342 | -0.71072 | 0.057356 | 2.91E-35 | 1.54E-30 |
| 15 | rs2568490 | 78738370 | T | C | 0.214083 | -0.71091 | 0.057382 | 2.99E-35 | 1.55E-30 |
| 15 | rs2915695 | 78739471 | T | C | 0.214063 | -0.7109 | 0.057378 | 2.97E-35 | 1.55E-30 |
| 15 | rs924840 | 78731808 | T | A | 0.213887 | -0.7116 | 0.057467 | 3.24E-35 | 1.64E-30 |
| 15 | rs10851906 | 78774676 | G | A | 0.214609 | -0.71007 | 0.057343 | 3.23E-35 | 1.64E-30 |
| 15 | rs2568488 | 78736593 | T | A | 0.214034 | -0.71059 | 0.057397 | 3.34E-35 | 1.67E-30 |
| 15 | rs2036529 | 78726272 | A | T | 0.213572 | -0.71135 | 0.057501 | 3.74E-35 | 1.80E-30 |

| | | | | | | | | | |
|---|---|---|---|---|---|---|---|---|---|
| 15 | rs2656070 | 78730252 | A | G | 0.213936 | -0.7111 | 0.057478 | 3.72E-35 | 1.80E-30 |
| 15 | rs2656071 | 78745343 | T | A | 0.213951 | -0.71014 | 0.057392 | 3.63E-35 | 1.80E-30 |
| 15 | rs2568483 | 78752343 | G | A | 0.214086 | -0.7101 | 0.057398 | 3.73E-35 | 1.80E-30 |
| 15 | rs2036528 | 78726271 | G | C | 0.213565 | -0.71102 | 0.057503 | 4.04E-35 | 1.92E-30 |
| 15 | rs2656056 | 78722519 | T | C | 0.21326 | -0.71152 | 0.057547 | 4.09E-35 | 1.92E-30 |
| 15 | rs7174348 | 78792439 | A | G | 0.214698 | -0.70874 | 0.057329 | 4.17E-35 | 1.94E-30 |
| 15 | rs28480606 | 78762313 | G | A | 0.21408 | -0.70886 | 0.057386 | 4.71E-35 | 2.17E-30 |
| 15 | rs7174190 | 78763617 | T | C | 0.214079 | -0.70862 | 0.057385 | 4.96E-35 | 2.27E-30 |
| 15 | rs8042260 | 78774374 | A | G | 0.378731 | -0.58689 | 0.047539 | 5.15E-35 | 2.33E-30 |
| 15 | rs12594711 | 78793921 | C | T | 0.379182 | -0.58616 | 0.047517 | 5.81E-35 | 2.60E-30 |
| 15 | rs16969894 | 78776456 | T | C | 0.214941 | -0.70688 | 0.05733 | 6.24E-35 | 2.77E-30 |
| 15 | rs12899351 | 78792398 | T | C | 0.378855 | -0.58578 | 0.047537 | 6.85E-35 | 3.01E-30 |
| 15 | rs12910910 | 78767850 | C | T | 0.378693 | -0.58568 | 0.047537 | 7.02E-35 | 3.06E-30 |
| 15 | rs4887059 | 78782095 | C | T | 0.379148 | -0.58525 | 0.047517 | 7.37E-35 | 3.18E-30 |
| 15 | rs965604 | 78789223 | G | A | 0.379144 | -0.58495 | 0.047517 | 7.97E-35 | 3.40E-30 |
| 15 | rs8043227 | 78768871 | C | G | 0.37863 | -0.5851 | 0.047539 | 8.22E-35 | 3.42E-30 |
| 15 | rs12916801 | 78769130 | A | G | 0.37863 | -0.5851 | 0.047539 | 8.22E-35 | 3.42E-30 |
| 15 | rs4362358 | 78796104 | C | T | 0.378892 | -0.58502 | 0.047531 | 8.16E-35 | 3.42E-30 |
| 15 | rs36146269 | 78779510 | T | A | 0.379138 | -0.58477 | 0.047517 | 8.36E-35 | 3.45E-30 |
| 15 | rs1504549 | 78766629 | C | T | 0.379088 | -0.58393 | 0.047517 | 1.04E-34 | 4.24E-30 |
| 15 | rs2938674 | 78757913 | A | C | 0.213938 | -0.70516 | 0.057403 | 1.10E-34 | 4.45E-30 |
| 15 | rs2568497 | 78721397 | G | T | 0.213188 | -0.70675 | 0.05758 | 1.24E-34 | 5.00E-30 |
| 15 | rs11637656 | 78751961 | C | T | 0.378297 | -0.58398 | 0.047606 | 1.36E-34 | 5.42E-30 |
| 15 | rs12593229 | 78765290 | T | G | 0.378287 | -0.58319 | 0.04755 | 1.40E-34 | 5.53E-30 |
| 15 | rs12592111 | 78767346 | G | A | 0.381977 | -0.58179 | 0.047452 | 1.48E-34 | 5.77E-30 |
| 15 | rs1062980 | 78792527 | C | T | 0.378384 | -0.58253 | 0.047533 | 1.57E-34 | 6.11E-30 |
| 15 | rs2656055 | 78720194 | C | A | 0.213309 | -0.70502 | 0.057598 | 1.89E-34 | 7.29E-30 |
| 15 | rs12904234 | 78779384 | C | T | 0.378796 | -0.58088 | 0.047527 | 2.37E-34 | 9.03E-30 |
| 15 | rs12903295 | 78778972 | A | G | 0.378754 | -0.58084 | 0.047527 | 2.40E-34 | 9.07E-30 |
| 15 | rs4299116 | 78766194 | T | A | 0.378972 | -0.58035 | 0.047532 | 2.76E-34 | 1.03E-29 |
| 15 | rs4887057 | 78760918 | A | G | 0.378098 | -0.57991 | 0.047555 | 3.33E-34 | 1.24E-29 |
| 15 | rs2869045 | 78718899 | T | C | 0.202761 | -0.70768 | 0.058673 | 1.69E-33 | 6.25E-29 |

| | | | | | | | | | |
|---|---|---|---|---|---|---|---|---|---|
| 15 | rs12441998 | 78929372 | G | A | 0.193172 | -0.71051 | 0.059074 | 2.55E-33 | 9.35E-29 |
| 15 | rs2869032 | 78714561 | C | T | 0.203156 | -0.70324 | 0.058596 | 3.49E-33 | 1.27E-28 |
| 15 | rs11636605 | 78928878 | A | G | 0.190594 | -0.71227 | 0.059483 | 4.83E-33 | 1.75E-28 |
| 19 | rs3865454 | 41342459 | T | G | 0.336231 | -0.56715 | 0.047461 | 6.51E-33 | 2.33E-28 |
| 19 | rs11667314 | 41340983 | T | C | 0.336189 | -0.566 | 0.04746 | 8.70E-33 | 3.09E-28 |
| 19 | rs7251570 | 41341750 | A | G | 0.336246 | -0.56486 | 0.04745 | 1.12E-32 | 3.96E-28 |
| 15 | rs11072768 | 78929478 | T | G | 0.191239 | -0.70478 | 0.059307 | 1.44E-32 | 5.04E-28 |
| 15 | rs11638830 | 78948319 | C | G | 0.41029 | 0.588827 | 0.049606 | 1.69E-32 | 5.89E-28 |
| 19 | rs11083569 | 41340321 | C | G | 0.336519 | -0.56268 | 0.047525 | 2.43E-32 | 8.39E-28 |
| 15 | rs11639372 | 78966655 | T | C | 0.410865 | 0.586707 | 0.049611 | 2.86E-32 | 9.78E-28 |
| 15 | rs6495314 | 78960529 | C | A | 0.411276 | 0.586057 | 0.049572 | 2.99E-32 | 1.02E-27 |
| 19 | rs12459249 | 41339896 | T | C | 0.334108 | -0.56107 | 0.047555 | 3.98E-32 | 1.34E-27 |
| 15 | rs12899135 | 78954379 | G | A | 0.410784 | 0.584219 | 0.049548 | 4.35E-32 | 1.46E-27 |
| 15 | rs11072785 | 78968229 | T | C | 0.410579 | 0.58437 | 0.049638 | 5.40E-32 | 1.80E-27 |
| 15 | rs12902602 | 78967401 | G | A | 0.410674 | 0.584074 | 0.049617 | 5.46E-32 | 1.80E-27 |
| 15 | rs1021071 | 78968179 | C | G | 0.410727 | 0.584083 | 0.049621 | 5.51E-32 | 1.81E-27 |
| 15 | rs4886579 | 78969256 | T | C | 0.409459 | 0.581543 | 0.049707 | 1.28E-31 | 4.18E-27 |
| 15 | rs922692 | 78984214 | A | C | 0.410384 | 0.58275 | 0.049894 | 1.62E-31 | 5.24E-27 |
| 15 | rs11638372 | 78983559 | T | C | 0.410435 | 0.581153 | 0.049897 | 2.38E-31 | 7.63E-27 |
| 15 | rs4886580 | 78969385 | G | T | 0.409997 | 0.577831 | 0.0497 | 3.03E-31 | 9.65E-27 |
| 15 | rs11072790 | 78992025 | T | C | 0.415045 | 0.570723 | 0.049704 | 1.62E-30 | 5.12E-26 |
| 15 | rs12050570 | 79012888 | T | C | 0.413594 | 0.572419 | 0.049942 | 2.05E-30 | 6.46E-26 |
| 15 | rs11634543 | 79005504 | T | C | 0.4139 | 0.569193 | 0.049719 | 2.40E-30 | 7.51E-26 |
| 15 | rs12050571 | 79013055 | G | A | 0.413593 | 0.57073 | 0.049955 | 3.14E-30 | 9.76E-26 |
| 15 | rs12910627 | 78994933 | C | G | 0.411853 | 0.565428 | 0.049821 | 7.49E-30 | 2.31E-25 |
| 15 | rs34225855 | 79022136 | G | C | 0.413388 | 0.563687 | 0.049936 | 1.50E-29 | 4.59E-25 |
| 15 | rs11629637 | 79019024 | T | C | 0.41332 | 0.563558 | 0.049938 | 1.55E-29 | 4.70E-25 |
| 15 | rs35583595 | 79020152 | G | C | 0.413332 | 0.563588 | 0.049939 | 1.55E-29 | 4.70E-25 |
| 15 | rs34563625 | 79020278 | C | G | 0.413338 | 0.563507 | 0.049938 | 1.57E-29 | 4.72E-25 |
| 15 | rs3829786 | 79019546 | A | C | 0.413345 | 0.56332 | 0.049939 | 1.64E-29 | 4.91E-25 |
| 15 | rs2869554 | 79021037 | C | T | 0.413309 | 0.562993 | 0.04994 | 1.78E-29 | 5.27E-25 |
| 15 | rs11072791 | 78997076 | A | C | 0.413017 | 0.56092 | 0.049821 | 2.10E-29 | 6.19E-25 |

| | | | | | | | | | |
|---|---|---|---|---|---|---|---|---|---|
| 15 | rs11072792 | 78999911 | G | A | 0.420772 | 0.534994 | 0.049423 | 2.62E-27 | 7.69E-23 |
| 15 | rs4887096 | 79053284 | C | T | 0.425375 | 0.537745 | 0.049757 | 3.17E-27 | 9.24E-23 |
| 15 | rs12916326 | 79054011 | G | A | 0.425475 | 0.537288 | 0.049762 | 3.55E-27 | 1.03E-22 |
| 15 | rs1809419 | 79053814 | A | G | 0.424283 | 0.53426 | 0.049768 | 6.97E-27 | 2.00E-22 |
| 15 | rs12916648 | 79054129 | T | C | 0.424325 | 0.534076 | 0.049764 | 7.18E-27 | 2.05E-22 |
| 15 | rs56354501 | 79033520 | G | A | 0.417789 | 0.532469 | 0.04995 | 1.57E-26 | 4.44E-22 |
| 15 | rs8031513 | 79034276 | C | T | 0.419937 | 0.528985 | 0.049922 | 3.10E-26 | 8.76E-22 |
| 15 | rs12906653 | 79052580 | A | G | 0.420542 | 0.527238 | 0.049863 | 3.94E-26 | 1.11E-21 |
| 15 | rs55834964 | 79049766 | A | C | 0.422422 | 0.526017 | 0.049805 | 4.49E-26 | 1.25E-21 |
| 15 | rs56195905 | 79040317 | C | A | 0.420124 | 0.524137 | 0.049928 | 8.84E-26 | 2.45E-21 |
| 19 | rs72549444 | 41351843 | A | G | 0.022834 | -1.55265 | 0.148264 | 1.16E-25 | 3.19E-21 |
| 19 | rs4803380 | 41351169 | T | C | 0.022814 | -1.55079 | 0.148372 | 1.43E-25 | 3.93E-21 |
| 19 | rs73931391 | 41407874 | G | A | 0.0227 | -1.53156 | 0.14859 | 6.53E-25 | 1.78E-20 |
| 15 | rs6495267 | 79057093 | A | G | 0.429655 | 0.511327 | 0.0497 | 7.96E-25 | 2.15E-20 |
| 15 | rs1809420 | 79056769 | C | T | 0.429661 | 0.511232 | 0.0497 | 8.11E-25 | 2.18E-20 |
| 15 | rs11633351 | 79056815 | T | C | 0.429688 | 0.511149 | 0.049698 | 8.22E-25 | 2.20E-20 |
| 15 | rs1807006 | 79062340 | G | C | 0.430458 | 0.510157 | 0.049627 | 8.69E-25 | 2.31E-20 |
| 15 | rs1807007 | 79062102 | G | T | 0.430456 | 0.510105 | 0.049628 | 8.80E-25 | 2.33E-20 |
| 15 | rs4887097 | 79057949 | C | T | 0.429782 | 0.51048 | 0.049682 | 9.13E-25 | 2.39E-20 |
| 15 | rs4887098 | 79057950 | C | G | 0.429782 | 0.51048 | 0.049682 | 9.13E-25 | 2.39E-20 |
| 19 | rs112292570 | 41345586 | T | A | 0.022631 | -1.52943 | 0.148873 | 9.29E-25 | 2.42E-20 |
| 15 | rs11635870 | 79064143 | G | C | 0.430427 | 0.509684 | 0.049632 | 9.70E-25 | 2.51E-20 |
| 15 | rs11635931 | 79064080 | A | G | 0.43043 | 0.509629 | 0.049631 | 9.79E-25 | 2.52E-20 |
| 15 | rs1809409 | 79063474 | T | A | 0.430443 | 0.509521 | 0.04963 | 1.00E-24 | 2.56E-20 |
| 19 | rs117540499 | 41406448 | A | G | 0.02263 | -1.52644 | 0.14882 | 1.10E-24 | 2.80E-20 |
| 15 | rs1810165 | 79059449 | G | A | 0.431113 | 0.508103 | 0.049644 | 1.38E-24 | 3.50E-20 |
| 19 | rs186051747 | 41342042 | A | G | 0.02254 | -1.52375 | 0.149096 | 1.61E-24 | 4.06E-20 |
| 19 | rs144002017 | 41335269 | G | T | 0.022637 | -1.51333 | 0.148788 | 2.67E-24 | 6.68E-20 |
| 19 | rs187430990 | 41343605 | T | C | 0.022624 | -1.51136 | 0.148893 | 3.29E-24 | 8.20E-20 |
| 19 | rs143303864 | 41336272 | A | G | 0.022603 | -1.51032 | 0.148886 | 3.52E-24 | 8.71E-20 |
| 15 | rs1809423 | 79059670 | C | T | 0.431506 | 0.503697 | 0.049657 | 3.54E-24 | 8.72E-20 |
| 15 | rs1809424 | 79059703 | C | T | 0.431498 | 0.503646 | 0.049656 | 3.57E-24 | 8.76E-20 |

| | | | | | | | | | |
|---|---|---|---|---|---|---|---|---|---|
| 15 | rs3894351 | 79059523 | A | C | 0.431504 | 0.503518 | 0.049656 | 3.67E-24 | 8.85E-20 |
| 15 | rs3894352 | 79059526 | G | T | 0.431504 | 0.503518 | 0.049656 | 3.67E-24 | 8.85E-20 |
| 15 | rs4887099 | 79059547 | C | G | 0.431504 | 0.503518 | 0.049656 | 3.67E-24 | 8.85E-20 |
| 19 | rs56267346 | 41353338 | G | A | 0.161362 | -0.61691 | 0.061121 | 5.93E-24 | 1.42E-19 |
| 19 | rs28399462 | 41350615 | A | G | 0.022835 | -1.49755 | 0.148389 | 5.99E-24 | 1.43E-19 |
| 19 | rs187795779 | 41343511 | C | T | 0.022207 | -1.50577 | 0.150299 | 1.26E-23 | 3.00E-19 |
| 19 | rs1496402 | 41366134 | T | A | 0.348346 | -0.46326 | 0.046329 | 1.53E-23 | 3.63E-19 |
| 19 | rs8102900 | 41366928 | G | A | 0.348811 | -0.46313 | 0.046332 | 1.59E-23 | 3.73E-19 |
| 19 | rs12610432 | 41367769 | T | C | 0.266531 | -0.49698 | 0.049733 | 1.64E-23 | 3.83E-19 |
| 19 | rs150944685 | 41348419 | A | G | 0.0223 | -1.48502 | 0.150143 | 4.57E-23 | 1.06E-18 |
| 15 | rs12594247 | 78946633 | T | C | 0.151755 | -0.63212 | 0.064878 | 1.97E-22 | 4.57E-18 |
| 15 | rs11072774 | 78952697 | T | C | 0.165824 | -0.61066 | 0.0629 | 2.78E-22 | 6.40E-18 |
| 19 | rs10404667 | 41362898 | T | C | 0.26383 | -0.48168 | 0.049947 | 5.22E-22 | 1.20E-17 |
| 19 | rs2644914 | 41358011 | T | C | 0.264638 | -0.4817 | 0.050068 | 6.53E-22 | 1.49E-17 |
| 15 | rs12148319 | 78956192 | G | A | 0.152376 | -0.61583 | 0.064798 | 2.02E-21 | 4.60E-17 |
| 15 | rs7167806 | 79092366 | G | C | 0.449383 | 0.464591 | 0.048917 | 2.15E-21 | 4.86E-17 |
| 15 | rs7173267 | 79092750 | C | G | 0.449361 | 0.464483 | 0.048913 | 2.18E-21 | 4.90E-17 |
| 15 | rs4887113 | 79095287 | T | C | 0.448957 | 0.464527 | 0.048935 | 2.25E-21 | 5.04E-17 |
| 15 | rs11638321 | 79092183 | C | T | 0.449386 | 0.46381 | 0.04891 | 2.47E-21 | 5.51E-17 |
| 15 | rs28610385 | 79090606 | A | C | 0.449371 | 0.46366 | 0.048911 | 2.55E-21 | 5.63E-17 |
| 15 | rs11634450 | 79093201 | G | A | 0.449371 | 0.46369 | 0.048913 | 2.54E-21 | 5.63E-17 |
| 15 | rs11856536 | 79094325 | G | A | 0.449333 | 0.463559 | 0.048914 | 2.62E-21 | 5.75E-17 |
| 15 | rs7163204 | 78953740 | A | G | 0.151781 | -0.61477 | 0.064913 | 2.78E-21 | 6.08E-17 |
| 15 | rs7164665 | 78953919 | C | T | 0.152111 | -0.61395 | 0.06486 | 2.92E-21 | 6.34E-17 |
| 15 | rs7177699 | 79089734 | C | T | 0.449504 | 0.462959 | 0.048911 | 2.93E-21 | 6.34E-17 |
| 15 | rs12595350 | 78953464 | A | G | 0.151535 | -0.61342 | 0.064954 | 3.59E-21 | 7.74E-17 |
| 15 | rs11639044 | 79083814 | T | C | 0.439615 | 0.461001 | 0.049249 | 7.93E-21 | 1.70E-16 |
| 15 | rs12050525 | 79075746 | C | T | 0.43948 | 0.462113 | 0.049392 | 8.28E-21 | 1.77E-16 |
| 15 | rs1825087 | 79077114 | G | A | 0.439668 | 0.460413 | 0.049281 | 9.41E-21 | 2.00E-16 |
| 15 | rs11632102 | 79086057 | A | G | 0.439167 | 0.459363 | 0.049286 | 1.16E-20 | 2.46E-16 |
| 15 | rs2277545 | 79083591 | C | T | 0.440085 | 0.458202 | 0.049225 | 1.30E-20 | 2.74E-16 |
| 15 | rs2277546 | 79083376 | A | C | 0.440091 | 0.458041 | 0.049223 | 1.34E-20 | 2.80E-16 |

| | | | | | | | | | |
|---|---|---|---|---|---|---|---|---|---|
| 15 | rs12903203 | 79084933 | C | T | 0.439881 | 0.458019 | 0.049247 | 1.40E-20 | 2.92E-16 |
| 15 | rs2904220 | 79076704 | T | G | 0.439619 | 0.45853 | 0.049354 | 1.53E-20 | 3.19E-16 |
| 15 | rs11858210 | 79078358 | A | G | 0.439356 | 0.456889 | 0.04919 | 1.57E-20 | 3.24E-16 |
| 15 | rs11631955 | 79085915 | G | A | 0.439638 | 0.457522 | 0.049273 | 1.61E-20 | 3.32E-16 |
| 15 | rs4886592 | 79082547 | C | T | 0.440113 | 0.456658 | 0.049199 | 1.66E-20 | 3.42E-16 |
| 15 | rs12899147 | 79079512 | G | A | 0.439847 | 0.456545 | 0.049197 | 1.70E-20 | 3.47E-16 |
| 15 | rs2869861 | 79076744 | G | A | 0.43889 | 0.456508 | 0.049398 | 2.43E-20 | 4.95E-16 |
| 15 | rs11072803 | 79077878 | A | G | 0.439776 | 0.454064 | 0.04919 | 2.68E-20 | 5.43E-16 |
| 19 | rs8192725 | 41354712 | A | G | 0.258313 | -0.44959 | 0.051307 | 1.91E-18 | 3.85E-14 |
| 19 | rs28399433 | 41356379 | C | A | 0.060983 | -0.76462 | 0.091133 | 4.85E-17 | 9.75E-13 |
| 19 | rs8192726 | 41354496 | A | C | 0.060178 | -0.77025 | 0.092587 | 8.86E-17 | 1.77E-12 |
| 19 | rs12986371 | 41343698 | G | A | 0.274177 | -0.41885 | 0.050512 | 1.11E-16 | 2.21E-12 |
| 19 | rs76112798 | 41343700 | T | C | 0.061287 | -0.75311 | 0.091867 | 2.45E-16 | 4.85E-12 |
| 19 | rs56007948 | 41340323 | T | C | 0.266285 | 0.409692 | 0.050427 | 4.49E-16 | 8.87E-12 |
| 19 | rs4343391 | 41344368 | G | C | 0.272518 | -0.40633 | 0.050669 | 1.06E-15 | 2.09E-11 |
| 19 | rs11668399 | 41343871 | C | G | 0.27155 | -0.40709 | 0.050777 | 1.08E-15 | 2.12E-11 |
| 15 | rs74634068 | 78965209 | A | G | 0.049778 | -0.81759 | 0.102518 | 1.52E-15 | 2.97E-11 |
| 19 | rs11083571 | 41346099 | C | T | 0.272007 | -0.40241 | 0.050732 | 2.15E-15 | 4.18E-11 |
| 15 | rs12324203 | 78961421 | A | G | 0.04985 | -0.81136 | 0.10244 | 2.37E-15 | 4.58E-11 |
| 19 | rs11878604 | 41333284 | C | T | 0.069787 | -0.68229 | 0.08663 | 3.38E-15 | 6.51E-11 |
| 19 | rs11879413 | 41337923 | T | C | 0.068578 | -0.68692 | 0.087254 | 3.47E-15 | 6.65E-11 |
| 15 | rs77163824 | 78972145 | G | A | 0.049734 | -0.80402 | 0.102605 | 4.65E-15 | 8.87E-11 |
| 19 | rs111272925 | 41370132 | T | C | 0.059332 | -0.72184 | 0.092159 | 4.78E-15 | 9.08E-11 |
| 19 | rs2258314 | 41338835 | T | C | 0.068429 | -0.68239 | 0.087364 | 5.68E-15 | 1.08E-10 |
| 15 | rs8043105 | 78973356 | T | C | 0.049751 | -0.801 | 0.102606 | 5.88E-15 | 1.11E-10 |
| 19 | rs10408411 | 41336950 | T | C | 0.261412 | -0.39898 | 0.051193 | 6.51E-15 | 1.22E-10 |
| 19 | rs2644899 | 41302949 | G | T | 0.288812 | -0.38397 | 0.049414 | 7.83E-15 | 1.46E-10 |
| 19 | rs7255471 | 41335066 | T | C | 0.294186 | 0.379406 | 0.048894 | 8.51E-15 | 1.58E-10 |
| 15 | rs144425814 | 78962803 | T | C | 0.05079 | -0.78215 | 0.10156 | 1.35E-14 | 2.50E-10 |
| 19 | rs11881918 | 41334199 | A | G | 0.068019 | -0.67466 | 0.087674 | 1.41E-14 | 2.61E-10 |
| 19 | rs2644891 | 41334371 | T | C | 0.329552 | -0.36466 | 0.047971 | 2.93E-14 | 5.38E-10 |
| 19 | rs12985907 | 41343544 | A | G | 0.230649 | -0.40192 | 0.053221 | 4.29E-14 | 7.86E-10 |

| | | | | | | | | | |
|---|---|---|---|---|---|---|---|---|---|
| 15 | rs11857532 | 78968268 | G | T | 0.461239 | 0.358677 | 0.047641 | 5.12E-14 | 9.36E-10 |
| 19 | rs17726276 | 41291119 | G | A | 0.496969 | -0.33578 | 0.044801 | 6.63E-14 | 1.21E-09 |
| 19 | rs2644916 | 41309211 | T | C | 0.250682 | -0.38548 | 0.051549 | 7.55E-14 | 1.37E-09 |
| 19 | rs11668644 | 41316746 | G | C | 0.249216 | -0.38557 | 0.051743 | 9.22E-14 | 1.66E-09 |
| 15 | rs7169223 | 78992116 | G | A | 0.35324 | -0.36895 | 0.05007 | 1.72E-13 | 3.10E-09 |
| 15 | rs76912891 | 78964464 | A | G | 0.067798 | -0.65015 | 0.088722 | 2.34E-13 | 4.18E-09 |
| 15 | rs4887064 | 78842847 | G | C | 0.440946 | -0.35319 | 0.048516 | 3.34E-13 | 5.96E-09 |
| 15 | rs28438420 | 78836288 | A | T | 0.447857 | -0.35216 | 0.048469 | 3.71E-13 | 6.59E-09 |
| 15 | rs1979908 | 78843564 | C | T | 0.441159 | -0.35038 | 0.048517 | 5.13E-13 | 9.09E-09 |
| 15 | rs12901241 | 79002662 | G | A | 0.356907 | -0.3604 | 0.049909 | 5.15E-13 | 9.09E-09 |
| 15 | rs1979907 | 78842239 | T | C | 0.441122 | -0.34956 | 0.048513 | 5.78E-13 | 1.01E-08 |
| 15 | rs1979906 | 78842289 | C | T | 0.441122 | -0.34956 | 0.048513 | 5.78E-13 | 1.01E-08 |
| 15 | rs11858230 | 78835552 | A | G | 0.439781 | -0.35028 | 0.048617 | 5.81E-13 | 1.01E-08 |
| 15 | rs1979905 | 78842374 | A | C | 0.441139 | -0.34946 | 0.048513 | 5.87E-13 | 1.02E-08 |
| 15 | rs7164030 | 78844661 | G | A | 0.441031 | -0.35008 | 0.048598 | 5.87E-13 | 1.02E-08 |
| 15 | rs905740 | 78844386 | T | C | 0.441173 | -0.34933 | 0.048523 | 6.05E-13 | 1.04E-08 |
| 15 | rs880395 | 78844356 | A | G | 0.441169 | -0.34907 | 0.048522 | 6.29E-13 | 1.08E-08 |
| 15 | rs4243083 | 78833830 | C | G | 0.43968 | -0.34956 | 0.048633 | 6.59E-13 | 1.13E-08 |
| 15 | rs1504548 | 78843246 | T | G | 0.441209 | -0.34863 | 0.048511 | 6.64E-13 | 1.13E-08 |
| 15 | rs12907966 | 78843051 | T | C | 0.441035 | -0.34849 | 0.048512 | 6.79E-13 | 1.15E-08 |
| 15 | rs35704045 | 79003292 | C | G | 0.356764 | -0.35846 | 0.049919 | 6.93E-13 | 1.17E-08 |
| 15 | rs2292117 | 78834689 | A | G | 0.439628 | -0.34892 | 0.048627 | 7.20E-13 | 1.22E-08 |
| 15 | rs3813572 | 78832588 | C | T | 0.439469 | -0.34914 | 0.04868 | 7.38E-13 | 1.24E-08 |
| 15 | rs1504546 | 78824235 | T | C | 0.439476 | -0.34888 | 0.048678 | 7.66E-13 | 1.26E-08 |
| 15 | rs11630349 | 78824608 | T | C | 0.439475 | -0.34885 | 0.048678 | 7.69E-13 | 1.26E-08 |
| 15 | rs12906284 | 78825105 | T | C | 0.439476 | -0.34888 | 0.048678 | 7.66E-13 | 1.26E-08 |
| 15 | rs12906846 | 78825295 | A | G | 0.439476 | -0.34888 | 0.048678 | 7.66E-13 | 1.26E-08 |
| 15 | rs56007453 | 78826239 | C | T | 0.439476 | -0.34891 | 0.048678 | 7.63E-13 | 1.26E-08 |
| 15 | rs17407257 | 78826249 | G | A | 0.439477 | -0.3489 | 0.048678 | 7.64E-13 | 1.26E-08 |
| 15 | rs12916999 | 78826912 | T | C | 0.439476 | -0.34887 | 0.048678 | 7.67E-13 | 1.26E-08 |
| 15 | rs1847530 | 78824031 | A | G | 0.439414 | -0.34868 | 0.048673 | 7.84E-13 | 1.28E-08 |
| 15 | rs8025429 | 78836362 | G | A | 0.439802 | -0.34817 | 0.048607 | 7.90E-13 | 1.29E-08 |

| | | | | | | | | | |
|---|---|---|---|---|---|---|---|---|---|
| 15 | rs12916483 | 78832397 | A | G | 0.439474 | -0.34825 | 0.04868 | 8.45E-13 | 1.37E-08 |
| 15 | rs4886571 | 78833758 | G | A | 0.439693 | -0.34764 | 0.048647 | 8.92E-13 | 1.44E-08 |
| 15 | rs28392948 | 78836866 | T | C | 0.439836 | -0.3471 | 0.048585 | 9.06E-13 | 1.46E-08 |
| 15 | rs2869544 | 78839400 | G | A | 0.439918 | -0.34582 | 0.048552 | 1.06E-12 | 1.70E-08 |
| 15 | rs4887063 | 78839715 | C | T | 0.439913 | -0.34577 | 0.048552 | 1.07E-12 | 1.71E-08 |
| 15 | rs55690619 | 78833612 | A | G | 0.439515 | -0.34624 | 0.04867 | 1.13E-12 | 1.80E-08 |
| 15 | rs7181405 | 78948152 | G | A | 0.373541 | -0.34925 | 0.049169 | 1.22E-12 | 1.94E-08 |
| 15 | rs4886572 | 78837452 | A | G | 0.439869 | -0.34457 | 0.048558 | 1.28E-12 | 2.03E-08 |
| 15 | rs77434238 | 78836726 | T | C | 0.440336 | -0.34405 | 0.048642 | 1.52E-12 | 2.39E-08 |
| 15 | rs76712448 | 78836719 | A | G | 0.440083 | -0.34269 | 0.048645 | 1.86E-12 | 2.90E-08 |
| 15 | rs75754317 | 78836721 | G | C | 0.440083 | -0.34269 | 0.048645 | 1.86E-12 | 2.90E-08 |
| 15 | rs78406067 | 78836723 | T | C | 0.440083 | -0.34269 | 0.048645 | 1.86E-12 | 2.90E-08 |
| 15 | rs3934898 | 79055278 | G | A | 0.332246 | -0.35751 | 0.05083 | 2.02E-12 | 3.14E-08 |
| 15 | rs62010555 | 79007555 | A | G | 0.348546 | -0.35135 | 0.050229 | 2.65E-12 | 4.12E-08 |
| 19 | rs11083581 | 41388949 | G | T | 0.272974 | -0.35732 | 0.051292 | 3.25E-12 | 5.03E-08 |
| 15 | rs35701817 | 79004968 | G | A | 0.348638 | -0.3496 | 0.050228 | 3.40E-12 | 5.24E-08 |
| 15 | rs922691 | 78963994 | G | A | 0.356978 | -0.34498 | 0.049761 | 4.12E-12 | 6.34E-08 |
| 15 | rs58526289 | 79012418 | C | T | 0.348303 | -0.34876 | 0.050334 | 4.24E-12 | 6.50E-08 |
| 15 | rs2929154 | 79048551 | G | A | 0.333295 | -0.35232 | 0.050894 | 4.43E-12 | 6.76E-08 |
| 15 | rs34115022 | 79052972 | G | C | 0.332088 | -0.35178 | 0.050821 | 4.45E-12 | 6.78E-08 |
| 15 | rs62010556 | 79012311 | A | G | 0.348324 | -0.34767 | 0.050328 | 4.91E-12 | 7.45E-08 |
| 15 | rs1045130 | 79051721 | A | G | 0.333043 | -0.35094 | 0.050863 | 5.21E-12 | 7.88E-08 |
| 19 | rs145365541 | 41432307 | A | G | 0.018448 | -1.12011 | 0.162497 | 5.46E-12 | 8.23E-08 |
| 15 | rs34550457 | 79012476 | A | G | 0.348294 | -0.3454 | 0.050346 | 6.87E-12 | 1.03E-07 |
| 15 | rs58946838 | 78961449 | C | A | 0.356535 | -0.34078 | 0.04978 | 7.61E-12 | 1.14E-07 |
| 19 | rs3875145 | 41285516 | C | T | 0.338051 | -0.32492 | 0.04757 | 8.48E-12 | 1.27E-07 |
| 15 | rs12905273 | 79002755 | G | A | 0.346757 | -0.34255 | 0.05016 | 8.55E-12 | 1.27E-07 |
| 19 | rs35061187 | 41316898 | T | C | 0.350471 | 0.3167 | 0.046489 | 9.60E-12 | 1.42E-07 |
| 19 | rs66889044 | 41338712 | T | C | 0.360375 | 0.315692 | 0.046428 | 1.05E-11 | 1.55E-07 |
| 15 | rs12910237 | 78956338 | T | C | 0.359214 | -0.33704 | 0.049691 | 1.18E-11 | 1.74E-07 |
| 19 | rs4803356 | 41207206 | G | C | 0.073295 | -0.57794 | 0.085608 | 1.47E-11 | 2.16E-07 |
| 19 | rs28602288 | 41388740 | C | T | 0.349817 | -0.3228 | 0.048111 | 1.95E-11 | 2.86E-07 |

| | | | | | | | | | |
|---|---|---|---|---|---|---|---|---|---|
| 15 | rs12901331 | 79021097 | C | T | 0.348309 | -0.33737 | 0.050414 | 2.20E-11 | 3.21E-07 |
| 15 | rs1964562 | 79046491 | G | C | 0.337428 | -0.33991 | 0.050815 | 2.24E-11 | 3.27E-07 |
| 15 | rs34422348 | 79020472 | G | A | 0.348498 | -0.33473 | 0.050414 | 3.15E-11 | 4.56E-07 |
| 15 | rs4887062 | 78837801 | G | A | 0.445688 | -0.32048 | 0.048442 | 3.70E-11 | 5.35E-07 |
| 8 | rs6987704 | 42547623 | T | C | 0.204265 | -0.38998 | 0.05931 | 4.85E-11 | 7.00E-07 |
| 19 | rs2644900 | 41303000 | G | A | 0.422249 | -0.2958 | 0.045292 | 6.54E-11 | 9.40E-07 |
| 15 | rs34275594 | 78981282 | C | A | 0.344953 | -0.32858 | 0.050348 | 6.75E-11 | 9.67E-07 |
| 19 | rs6508949 | 41371110 | A | C | 0.140722 | -0.41312 | 0.063412 | 7.28E-11 | 1.04E-06 |
| 15 | rs61630816 | 78965866 | C | T | 0.344372 | -0.32663 | 0.050184 | 7.58E-11 | 1.08E-06 |
| 15 | rs34019568 | 79044233 | T | C | 0.338412 | -0.33056 | 0.050793 | 7.62E-11 | 1.08E-06 |
| 15 | rs7168915 | 79128889 | G | A | 0.463374 | 0.317893 | 0.048853 | 7.66E-11 | 1.08E-06 |
| 15 | rs34694149 | 78965917 | T | C | 0.34437 | -0.32653 | 0.050184 | 7.68E-11 | 1.08E-06 |
| 8 | rs6984252 | 42644651 | G | C | 0.205566 | -0.38519 | 0.059211 | 7.75E-11 | 1.09E-06 |
| 8 | rs4567031 | 42646055 | A | G | 0.205239 | -0.38426 | 0.059276 | 9.02E-11 | 1.27E-06 |
| 15 | rs59414039 | 79043766 | C | T | 0.338391 | -0.32878 | 0.050796 | 9.63E-11 | 1.35E-06 |
| 15 | rs4551997 | 79130433 | G | A | 0.496149 | 0.309636 | 0.047867 | 9.89E-11 | 1.38E-06 |
| 19 | rs2545758 | 41297303 | A | C | 0.262016 | -0.32803 | 0.050912 | 1.17E-10 | 1.63E-06 |
| 8 | rs4509313 | 42645846 | G | C | 0.205277 | -0.38172 | 0.059263 | 1.19E-10 | 1.64E-06 |
| 8 | rs1955186 | 42549491 | G | C | 0.225671 | -0.37508 | 0.058252 | 1.20E-10 | 1.66E-06 |
| 8 | rs6474413 | 42551064 | C | T | 0.225737 | -0.37491 | 0.058245 | 1.22E-10 | 1.68E-06 |
| 8 | rs4951 | 42563557 | C | T | 0.225833 | -0.37437 | 0.058267 | 1.32E-10 | 1.81E-06 |
| 19 | rs2545757 | 41297302 | A | G | 0.2621 | -0.32698 | 0.050905 | 1.33E-10 | 1.83E-06 |
| 8 | rs10958727 | 42554763 | C | G | 0.225739 | -0.3741 | 0.058249 | 1.34E-10 | 1.83E-06 |
| 15 | rs8034804 | 79129600 | C | T | 0.495841 | 0.306864 | 0.047788 | 1.35E-10 | 1.84E-06 |
| 8 | rs6985052 | 42551319 | C | T | 0.22573 | -0.37391 | 0.058247 | 1.37E-10 | 1.86E-06 |
| 15 | rs12903613 | 79129076 | G | T | 0.49586 | 0.30668 | 0.047791 | 1.39E-10 | 1.88E-06 |
| 8 | rs13277524 | 42550057 | G | T | 0.2256 | -0.37357 | 0.058239 | 1.41E-10 | 1.91E-06 |
| 8 | rs9643853 | 42556652 | A | C | 0.225523 | -0.37346 | 0.058247 | 1.44E-10 | 1.94E-06 |
| 15 | rs12903668 | 79129320 | A | G | 0.495855 | 0.306348 | 0.047789 | 1.45E-10 | 1.95E-06 |
| 8 | rs1955185 | 42549647 | C | T | 0.225569 | -0.37311 | 0.058236 | 1.49E-10 | 1.98E-06 |
| 8 | rs9643891 | 42556597 | C | T | 0.225528 | -0.37317 | 0.058247 | 1.49E-10 | 1.98E-06 |
| 15 | rs7178007 | 78965254 | A | G | 0.346029 | -0.3211 | 0.050123 | 1.49E-10 | 1.99E-06 |

| | | | | | | | | | |
|---|---|---|---|---|---|---|---|---|---|
| 8 | rs7004381 | 42551161 | A | G | 0.225647 | -0.37306 | 0.058241 | 1.50E-10 | 1.99E-06 |
| 8 | rs9792277 | 42545827 | G | A | 0.225585 | -0.37284 | 0.058222 | 1.52E-10 | 2.01E-06 |
| 8 | rs13277254 | 42549982 | G | A | 0.225552 | -0.37248 | 0.058238 | 1.60E-10 | 2.11E-06 |
| 8 | rs11783507 | 42534395 | C | A | 0.225244 | -0.37246 | 0.058244 | 1.61E-10 | 2.11E-06 |
| 8 | rs4736835 | 42547033 | T | C | 0.225551 | -0.37228 | 0.058228 | 1.62E-10 | 2.13E-06 |
| 8 | rs1451239 | 42546542 | G | A | 0.225548 | -0.37225 | 0.058228 | 1.63E-10 | 2.13E-06 |
| 8 | rs6474415 | 42562938 | G | A | 0.225664 | -0.37227 | 0.058241 | 1.64E-10 | 2.14E-06 |
| 8 | rs7816726 | 42535437 | A | G | 0.225312 | -0.37214 | 0.058247 | 1.67E-10 | 2.17E-06 |
| 8 | rs77232073 | 42542356 | G | C | 0.225308 | -0.3722 | 0.058255 | 1.67E-10 | 2.17E-06 |
| 8 | rs4736836 | 42547176 | C | T | 0.225592 | -0.37201 | 0.058238 | 1.68E-10 | 2.18E-06 |
| 8 | rs6997909 | 42560249 | A | G | 0.225562 | -0.37176 | 0.058226 | 1.72E-10 | 2.22E-06 |
| 8 | rs13254578 | 42545846 | G | C | 0.225509 | -0.3717 | 0.058227 | 1.73E-10 | 2.22E-06 |
| 8 | rs6474414 | 42560336 | A | C | 0.225574 | -0.37164 | 0.058228 | 1.74E-10 | 2.23E-06 |
| 8 | rs4736837 | 42547183 | C | T | 0.225586 | -0.37164 | 0.058238 | 1.76E-10 | 2.25E-06 |
| 8 | rs35599391 | 42540066 | C | T | 0.2253 | -0.37149 | 0.058252 | 1.80E-10 | 2.30E-06 |
| 8 | rs4950 | 42552633 | G | A | 0.225515 | -0.3713 | 0.05825 | 1.84E-10 | 2.34E-06 |
| 8 | rs4236926 | 42578059 | T | G | 0.229218 | -0.36949 | 0.057971 | 1.85E-10 | 2.34E-06 |
| 8 | rs78258002 | 42542494 | T | G | 0.225249 | -0.37113 | 0.058252 | 1.88E-10 | 2.38E-06 |
| 8 | rs4736838 | 42547333 | T | C | 0.225525 | -0.37096 | 0.05824 | 1.90E-10 | 2.39E-06 |
| 8 | rs9792257 | 42545551 | C | T | 0.225576 | -0.37094 | 0.05824 | 1.90E-10 | 2.39E-06 |
| 8 | rs4295650 | 42537811 | G | A | 0.225246 | -0.37085 | 0.058253 | 1.94E-10 | 2.43E-06 |
| 8 | rs7842601 | 42537055 | C | T | 0.225221 | -0.37072 | 0.058252 | 1.96E-10 | 2.46E-06 |
| 8 | rs1979140 | 42530836 | T | C | 0.225315 | -0.37066 | 0.058258 | 1.99E-10 | 2.48E-06 |
| 8 | rs9692914 | 42545296 | T | C | 0.225538 | -0.37031 | 0.058251 | 2.05E-10 | 2.56E-06 |
| 8 | rs6474411 | 42541446 | A | G | 0.225265 | -0.37021 | 0.058253 | 2.08E-10 | 2.58E-06 |
| 8 | rs9693858 | 42545357 | C | T | 0.225773 | -0.3699 | 0.058261 | 2.17E-10 | 2.68E-06 |
| 8 | rs57645595 | 42579025 | T | A | 0.229163 | -0.36764 | 0.057986 | 2.29E-10 | 2.83E-06 |
| 8 | rs9693825 | 42545177 | C | T | 0.225643 | -0.36898 | 0.058234 | 2.36E-10 | 2.89E-06 |
| 15 | rs1878399 | 78912003 | G | C | 0.447717 | -0.30731 | 0.048501 | 2.35E-10 | 2.89E-06 |
| 8 | rs16891561 | 42579739 | T | C | 0.229036 | -0.36742 | 0.058018 | 2.41E-10 | 2.95E-06 |
| 8 | rs58379124 | 42579203 | T | C | 0.229133 | -0.36708 | 0.057995 | 2.46E-10 | 3.01E-06 |
| 15 | rs8030937 | 79129587 | G | T | 0.496784 | 0.302199 | 0.047798 | 2.57E-10 | 3.14E-06 |

| | | | | | | | | | |
|---|---|---|---|---|---|---|---|---|---|
| 15 | rs8024048 | 79127907 | G | A | 0.496233 | 0.30311 | 0.047949 | 2.59E-10 | 3.15E-06 |
| 19 | rs2604893 | 41293543 | A | G | 0.262426 | -0.32162 | 0.050885 | 2.60E-10 | 3.16E-06 |
| 15 | rs62011004 | 79126536 | C | T | 0.496103 | 0.302631 | 0.047982 | 2.84E-10 | 3.43E-06 |
| 15 | rs12908341 | 79126561 | T | G | 0.496098 | 0.302615 | 0.047982 | 2.85E-10 | 3.43E-06 |
| 15 | rs7169511 | 79128665 | T | C | 0.496318 | 0.302183 | 0.04791 | 2.84E-10 | 3.43E-06 |
| 15 | rs7178750 | 79126909 | G | A | 0.49609 | 0.302344 | 0.04797 | 2.92E-10 | 3.51E-06 |
| 15 | rs6495339 | 79128272 | T | C | 0.496165 | 0.302017 | 0.04794 | 2.98E-10 | 3.56E-06 |
| 8 | rs13273442 | 42544017 | A | G | 0.225665 | -0.36654 | 0.058225 | 3.07E-10 | 3.67E-06 |
| 8 | rs7459838 | 42584279 | G | A | 0.228676 | -0.36506 | 0.058064 | 3.23E-10 | 3.85E-06 |
| 15 | rs4539564 | 79128499 | G | A | 0.496235 | 0.301163 | 0.047928 | 3.31E-10 | 3.93E-06 |
| 15 | rs6495340 | 79128396 | A | G | 0.496215 | 0.300854 | 0.047934 | 3.47E-10 | 4.11E-06 |
| 15 | rs588765 | 78865425 | T | C | 0.442653 | -0.30454 | 0.048625 | 3.78E-10 | 4.46E-06 |
| 15 | rs647041 | 78880481 | T | C | 0.441206 | -0.30447 | 0.048688 | 4.01E-10 | 4.73E-06 |
| 15 | rs4366683 | 78912203 | C | T | 0.449587 | -0.30286 | 0.048455 | 4.10E-10 | 4.82E-06 |
| 15 | rs481134 | 78877563 | A | G | 0.442702 | -0.30366 | 0.048626 | 4.24E-10 | 4.98E-06 |
| 8 | rs1530848 | 42552908 | G | T | 0.227747 | -0.36213 | 0.05808 | 4.52E-10 | 5.29E-06 |
| 15 | rs555018 | 78879242 | G | A | 0.442897 | -0.30337 | 0.048688 | 4.64E-10 | 5.41E-06 |
| 19 | rs7249450 | 41300771 | T | C | 0.262853 | -0.31635 | 0.050776 | 4.66E-10 | 5.42E-06 |
| 8 | rs34456987 | 42523329 | G | A | 0.224873 | -0.36349 | 0.058391 | 4.81E-10 | 5.59E-06 |
| 15 | rs59709310 | 79061214 | C | T | 0.332202 | -0.31609 | 0.050826 | 5.00E-10 | 5.79E-06 |
| 15 | rs12901300 | 78892952 | A | G | 0.44181 | -0.30243 | 0.048651 | 5.09E-10 | 5.88E-06 |
| 15 | rs12911602 | 78891449 | C | T | 0.441857 | -0.30231 | 0.048648 | 5.16E-10 | 5.95E-06 |
| 19 | rs2604869 | 41283693 | A | G | 0.264736 | -0.31543 | 0.050834 | 5.47E-10 | 6.28E-06 |
| 19 | rs2604895 | 41292263 | T | A | 0.264326 | -0.31529 | 0.050813 | 5.47E-10 | 6.28E-06 |
| 15 | rs56276142 | 78889795 | C | T | 0.441865 | -0.30128 | 0.048643 | 5.88E-10 | 6.73E-06 |
| 19 | rs2545755 | 41293162 | T | G | 0.262201 | -0.31545 | 0.050959 | 6.00E-10 | 6.85E-06 |
| 15 | rs4352017 | 78912204 | A | C | 0.44929 | -0.29994 | 0.048464 | 6.05E-10 | 6.89E-06 |
| 15 | rs11857877 | 79140838 | C | G | 0.46202 | 0.30173 | 0.048864 | 6.62E-10 | 7.52E-06 |
| 15 | rs6495306 | 78865893 | G | A | 0.442765 | -0.29997 | 0.048624 | 6.87E-10 | 7.78E-06 |
| 19 | rs3776611 | 41307778 | T | C | 0.361086 | 0.28469 | 0.046155 | 6.91E-10 | 7.81E-06 |
| 8 | rs13280301 | 42550017 | A | G | 0.168199 | -0.38883 | 0.063239 | 7.82E-10 | 8.82E-06 |
| 19 | rs4802090 | 41315234 | G | C | 0.361929 | 0.283061 | 0.046047 | 7.89E-10 | 8.87E-06 |

| | | | | | | | | | |
|---|---|---|---|---|---|---|---|---|---|
| 15 | rs57945453 | 78862845 | T | C | 0.384078 | -0.30372 | 0.049414 | 7.93E-10 | 8.89E-06 |
| 8 | rs10958725 | 42524584 | T | G | 0.225169 | -0.35832 | 0.058342 | 8.17E-10 | 9.14E-06 |
| 8 | rs11783289 | 42574596 | C | T | 0.156257 | -0.39832 | 0.064874 | 8.26E-10 | 9.23E-06 |
| 15 | rs8037171 | 79139478 | A | C | 0.46034 | 0.299433 | 0.048786 | 8.37E-10 | 9.33E-06 |
| 15 | rs4439728 | 79138908 | G | A | 0.460339 | 0.299125 | 0.048786 | 8.71E-10 | 9.68E-06 |
| 8 | rs1530847 | 42548239 | C | T | 0.168149 | -0.38751 | 0.063243 | 8.93E-10 | 9.91E-06 |
| 15 | rs4420501 | 79138478 | T | A | 0.460309 | 0.298767 | 0.048785 | 9.12E-10 | 1.01E-05 |
| 15 | rs11637783 | 79139000 | C | T | 0.460233 | 0.298811 | 0.048791 | 9.11E-10 | 1.01E-05 |
| 15 | rs4438276 | 79138565 | G | A | 0.460203 | 0.298462 | 0.048791 | 9.52E-10 | 1.05E-05 |
| 15 | rs3743077 | 78894896 | T | C | 0.441319 | -0.29749 | 0.048651 | 9.67E-10 | 1.06E-05 |
| 19 | rs4803369 | 41315980 | A | G | 0.361308 | 0.281777 | 0.04609 | 9.74E-10 | 1.07E-05 |
| 8 | rs13263434 | 42573053 | A | G | 0.156237 | -0.3963 | 0.06486 | 9.96E-10 | 1.09E-05 |
| 15 | rs12232282 | 79136525 | C | T | 0.460192 | 0.298028 | 0.048788 | 1.00E-09 | 1.10E-05 |
| 15 | rs28580532 | 79134724 | T | C | 0.460274 | 0.297153 | 0.048745 | 1.09E-09 | 1.18E-05 |
| 15 | rs62012623 | 79062888 | A | G | 0.331529 | -0.30996 | 0.050857 | 1.10E-09 | 1.19E-05 |
| 15 | rs28694044 | 79134718 | T | A | 0.460276 | 0.29628 | 0.048745 | 1.22E-09 | 1.32E-05 |
| 15 | rs2456019 | 78868489 | T | G | 0.384365 | -0.29988 | 0.049349 | 1.23E-09 | 1.33E-05 |
| 19 | rs4803368 | 41301714 | A | G | 0.261878 | -0.30874 | 0.050837 | 1.25E-09 | 1.35E-05 |
| 15 | rs4567668 | 79140692 | C | T | 0.46029 | 0.2961 | 0.048786 | 1.28E-09 | 1.38E-05 |
| 15 | rs5029904 | 79152422 | G | C | 0.463339 | 0.29511 | 0.048671 | 1.33E-09 | 1.43E-05 |
| 15 | rs11072812 | 79140111 | T | C | 0.46027 | 0.295529 | 0.048784 | 1.38E-09 | 1.48E-05 |
| 15 | rs12911870 | 79148650 | T | C | 0.463495 | 0.294676 | 0.048661 | 1.40E-09 | 1.49E-05 |
| 15 | rs4344704 | 79141703 | A | T | 0.460804 | 0.29535 | 0.048797 | 1.42E-09 | 1.52E-05 |
| 15 | rs11632020 | 79143759 | T | C | 0.46014 | 0.295112 | 0.048781 | 1.45E-09 | 1.54E-05 |
| 15 | rs4132786 | 79141588 | A | G | 0.460143 | 0.294923 | 0.048792 | 1.50E-09 | 1.59E-05 |
| 15 | rs7173743 | 79141784 | C | T | 0.460779 | 0.294934 | 0.048794 | 1.50E-09 | 1.59E-05 |
| 15 | rs601079 | 78869579 | T | A | 0.444033 | -0.29333 | 0.048586 | 1.57E-09 | 1.65E-05 |
| 15 | rs621849 | 78872861 | G | A | 0.44405 | -0.29326 | 0.048585 | 1.58E-09 | 1.66E-05 |
| 15 | rs692780 | 78876505 | C | G | 0.38443 | -0.29753 | 0.049341 | 1.64E-09 | 1.72E-05 |
| 19 | rs2545774 | 41287674 | T | C | 0.264608 | -0.3057 | 0.050704 | 1.65E-09 | 1.73E-05 |
| 15 | rs651209 | 78874838 | G | A | 0.444053 | -0.29259 | 0.048593 | 1.73E-09 | 1.81E-05 |
| 15 | rs77681598 | 78906103 | C | A | 0.444767 | -0.29243 | 0.048644 | 1.84E-09 | 1.92E-05 |

| | | | | | | | | | |
|---|---|---|---|---|---|---|---|---|---|
| 15 | rs12903249 | 79151860 | T | C | 0.463542 | 0.292442 | 0.048649 | 1.84E-09 | 1.92E-05 |
| 8 | rs6474418 | 42631284 | C | G | 0.223153 | -0.34564 | 0.057524 | 1.87E-09 | 1.95E-05 |
| 15 | rs12903542 | 79151999 | T | C | 0.463515 | 0.292152 | 0.048649 | 1.91E-09 | 1.98E-05 |
| 15 | rs139978901 | 78902026 | A | C | 0.443181 | -0.29156 | 0.048614 | 2.01E-09 | 2.08E-05 |
| 15 | rs62010327 | 78892784 | A | G | 0.385223 | -0.29662 | 0.049464 | 2.01E-09 | 2.08E-05 |
| 8 | rs10107450 | 42629895 | T | C | 0.222999 | -0.34457 | 0.057493 | 2.06E-09 | 2.12E-05 |
| 15 | rs2869546 | 78907345 | C | T | 0.39029 | -0.29537 | 0.049308 | 2.10E-09 | 2.15E-05 |
| 8 | rs10110473 | 42634630 | T | A | 0.18646 | -0.36865 | 0.061577 | 2.14E-09 | 2.20E-05 |
| 15 | rs2067808 | 78911780 | A | G | 0.390367 | -0.29459 | 0.049234 | 2.18E-09 | 2.24E-05 |
| 15 | rs8040544 | 78904261 | A | G | 0.440838 | -0.29127 | 0.048737 | 2.28E-09 | 2.33E-05 |
| 15 | rs146172900 | 78903850 | A | C | 0.44312 | -0.29055 | 0.048632 | 2.31E-09 | 2.35E-05 |
| 15 | rs7182583 | 78899210 | C | G | 0.384449 | -0.2942 | 0.049374 | 2.54E-09 | 2.58E-05 |
| 15 | rs190259891 | 78902906 | A | G | 0.443149 | -0.28976 | 0.048629 | 2.54E-09 | 2.58E-05 |
| 15 | rs77403874 | 78903165 | C | T | 0.443131 | -0.2898 | 0.048631 | 2.54E-09 | 2.58E-05 |
| 15 | rs183822442 | 78902775 | G | A | 0.443142 | -0.28968 | 0.04863 | 2.57E-09 | 2.60E-05 |
| 15 | rs3743057 | 79089007 | T | C | 0.254299 | -0.31652 | 0.053311 | 2.90E-09 | 2.92E-05 |
| 15 | rs61012457 | 78865694 | G | C | 0.384298 | -0.29277 | 0.04935 | 2.98E-09 | 3.00E-05 |
| 19 | rs11547373 | 41306362 | A | G | 0.356349 | 0.27515 | 0.046382 | 2.99E-09 | 3.00E-05 |
| 15 | rs1994017 | 79080306 | T | C | 0.257015 | -0.3146 | 0.053095 | 3.12E-09 | 3.12E-05 |
| 15 | rs4390557 | 78717905 | C | T | 0.467601 | -0.2804 | 0.047532 | 3.65E-09 | 3.65E-05 |
| 15 | rs1976007 | 79085023 | T | C | 0.256915 | -0.31248 | 0.053093 | 3.97E-09 | 3.96E-05 |
| 15 | rs471889 | 78870235 | T | C | 0.38596 | -0.28926 | 0.049282 | 4.37E-09 | 4.35E-05 |
| 15 | rs495956 | 78869930 | C | T | 0.385957 | -0.28912 | 0.049283 | 4.45E-09 | 4.42E-05 |
| 15 | rs12905740 | 79082364 | T | C | 0.256753 | -0.31142 | 0.053114 | 4.54E-09 | 4.50E-05 |
| 8 | rs4392883 | 42541155 | T | C | 0.189398 | -0.36076 | 0.061585 | 4.69E-09 | 4.63E-05 |
| 15 | rs111725959 | 78899346 | A | G | 0.384096 | -0.28901 | 0.049378 | 4.83E-09 | 4.76E-05 |
| 15 | rs495090 | 78870003 | A | G | 0.385885 | -0.28831 | 0.049285 | 4.92E-09 | 4.84E-05 |
| 15 | rs660652 | 78887832 | A | G | 0.383439 | -0.28885 | 0.049379 | 4.92E-09 | 4.84E-05 |
| 15 | rs12901971 | 78942159 | T | G | 0.214978 | -0.33678 | 0.057653 | 5.17E-09 | 5.07E-05 |
| 15 | rs10851911 | 79077666 | T | C | 0.257333 | -0.31028 | 0.053125 | 5.20E-09 | 5.09E-05 |
| 15 | rs472054 | 78887994 | A | G | 0.383704 | -0.28816 | 0.049367 | 5.31E-09 | 5.18E-05 |
| 15 | rs2904228 | 79086099 | A | G | 0.255127 | -0.31007 | 0.053253 | 5.80E-09 | 5.65E-05 |

| | | | | | | | | | |
|---|---|---|---|---|---|---|---|---|---|
| 15 | rs12902294 | 78942339 | T | C | 0.214991 | -0.33543 | 0.057645 | 5.92E-09 | 5.76E-05 |
| 15 | rs11072810 | 79132206 | T | C | 0.493513 | 0.277508 | 0.047768 | 6.27E-09 | 6.08E-05 |
| 15 | rs615470 | 78885988 | T | C | 0.384474 | -0.28687 | 0.049385 | 6.29E-09 | 6.09E-05 |
| 15 | rs62010328 | 78894971 | T | C | 0.378296 | -0.28778 | 0.049562 | 6.38E-09 | 6.16E-05 |
| 15 | rs11072811 | 79132330 | A | C | 0.493169 | 0.276974 | 0.047769 | 6.70E-09 | 6.45E-05 |
| 15 | rs8035783 | 79135481 | G | T | 0.390569 | -0.28116 | 0.048489 | 6.69E-09 | 6.45E-05 |
| 15 | rs584135 | 78883628 | A | G | 0.38426 | -0.28635 | 0.04939 | 6.72E-09 | 6.46E-05 |
| 15 | rs514743 | 78884227 | T | A | 0.384072 | -0.28629 | 0.04939 | 6.77E-09 | 6.48E-05 |
| 15 | rs4533253 | 79138841 | T | C | 0.392018 | -0.28048 | 0.048436 | 7.01E-09 | 6.70E-05 |
| 15 | rs11632963 | 79132644 | G | A | 0.493603 | 0.276617 | 0.047772 | 7.03E-09 | 6.71E-05 |
| 8 | rs1072003 | 42620001 | G | C | 0.186116 | -0.35186 | 0.060772 | 7.04E-09 | 6.71E-05 |
| 15 | rs8037017 | 79139441 | A | C | 0.392052 | -0.27995 | 0.048438 | 7.49E-09 | 7.12E-05 |
| 15 | rs3743074 | 78909480 | G | A | 0.383774 | -0.28524 | 0.049369 | 7.58E-09 | 7.19E-05 |
| 15 | rs7403442 | 79136123 | G | A | 0.391351 | -0.27967 | 0.048447 | 7.80E-09 | 7.38E-05 |
| 15 | rs17408276 | 78881618 | C | T | 0.383987 | -0.28508 | 0.049395 | 7.86E-09 | 7.43E-05 |
| 15 | rs3743073 | 78909539 | G | T | 0.383863 | -0.28489 | 0.049372 | 7.91E-09 | 7.46E-05 |
| 15 | rs7403393 | 79135802 | G | C | 0.391368 | -0.27953 | 0.048447 | 7.94E-09 | 7.47E-05 |
| 15 | rs4887074 | 78952110 | G | C | 0.22937 | -0.32773 | 0.05681 | 7.98E-09 | 7.50E-05 |
| 15 | rs7403438 | 79136048 | T | C | 0.391348 | -0.27943 | 0.048447 | 8.03E-09 | 7.52E-05 |
| 19 | rs8100418 | 41414363 | T | A | 0.372534 | 0.269336 | 0.046731 | 8.24E-09 | 7.71E-05 |
| 8 | rs2304297 | 42608199 | C | G | 0.22917 | -0.32929 | 0.057142 | 8.28E-09 | 7.73E-05 |
| 15 | rs4608303 | 79136713 | C | T | 0.391339 | -0.27909 | 0.048446 | 8.38E-09 | 7.80E-05 |
| 15 | rs12912524 | 78952756 | G | A | 0.22948 | -0.32666 | 0.056789 | 8.81E-09 | 8.19E-05 |
| 15 | rs2568498 | 78721932 | T | A | 0.45875 | -0.27401 | 0.047641 | 8.85E-09 | 8.21E-05 |
| 15 | rs28439027 | 79135478 | C | A | 0.390191 | -0.27889 | 0.048499 | 8.91E-09 | 8.25E-05 |
| 15 | rs8027870 | 79134045 | G | A | 0.391502 | -0.27781 | 0.048358 | 9.20E-09 | 8.50E-05 |
| 15 | rs7164529 | 79145798 | A | G | 0.390811 | -0.27829 | 0.048476 | 9.42E-09 | 8.68E-05 |
| 19 | rs28417358 | 41428105 | A | G | 0.372779 | 0.267471 | 0.046592 | 9.43E-09 | 8.68E-05 |
| 15 | rs11856903 | 79140017 | A | G | 0.391815 | -0.27756 | 0.048443 | 1.01E-08 | 9.24E-05 |
| 15 | rs7166723 | 79133848 | T | C | 0.391523 | -0.27683 | 0.048359 | 1.04E-08 | 9.51E-05 |
| 15 | rs4341710 | 79140460 | C | T | 0.392071 | -0.27722 | 0.048436 | 1.04E-08 | 9.54E-05 |
| 15 | rs7175271 | 79144509 | A | G | 0.39172 | -0.27727 | 0.048457 | 1.05E-08 | 9.62E-05 |

| | | | | | | | | | |
|---|---|---|---|---|---|---|---|---|---|
| 15 | rs8029173 | 79134568 | A | G | 0.391495 | -0.27681 | 0.04838 | 1.06E-08 | 9.62E-05 |
| 15 | rs2656058 | 78723411 | T | G | 0.458052 | -0.27234 | 0.047613 | 1.07E-08 | 9.70E-05 |
| 15 | rs7173955 | 79143857 | G | A | 0.391438 | -0.27709 | 0.048453 | 1.07E-08 | 9.72E-05 |
| 19 | rs3843043 | 41433931 | C | T | 0.261911 | -0.29589 | 0.051738 | 1.07E-08 | 9.72E-05 |
| 19 | rs3844444 | 41434047 | A | G | 0.261929 | -0.29577 | 0.051739 | 1.09E-08 | 9.83E-05 |
| 15 | rs8028410 | 79134267 | T | C | 0.391512 | -0.27619 | 0.04836 | 1.12E-08 | 0.000101 |
| 15 | rs7495518 | 78900574 | T | C | 0.377351 | -0.2831 | 0.049593 | 1.14E-08 | 0.000103 |
| 19 | rs4359558 | 41422518 | A | G | 0.371542 | 0.266287 | 0.046672 | 1.16E-08 | 0.000104 |
| 15 | rs4401016 | 79134791 | A | G | 0.391278 | -0.27617 | 0.048412 | 1.17E-08 | 0.000105 |
| 19 | rs4609955 | 41433613 | C | T | 0.3726 | 0.265677 | 0.046619 | 1.21E-08 | 0.000108 |
| 19 | rs7255901 | 41435736 | C | T | 0.372933 | 0.265436 | 0.046596 | 1.22E-08 | 0.000109 |
| 19 | rs12151139 | 41433543 | T | C | 0.372599 | 0.265524 | 0.046618 | 1.23E-08 | 0.00011 |
| 15 | rs12910090 | 78743300 | C | A | 0.45437 | -0.2712 | 0.047654 | 1.26E-08 | 0.000112 |
| 19 | rs12150973 | 41434524 | C | T | 0.372941 | 0.265162 | 0.046596 | 1.27E-08 | 0.000112 |
| 15 | rs8041818 | 79139831 | T | C | 0.391443 | -0.27556 | 0.04845 | 1.29E-08 | 0.000114 |
| 15 | rs11639224 | 78753371 | G | A | 0.453939 | -0.27083 | 0.047667 | 1.33E-08 | 0.000118 |
| 15 | rs12915669 | 78950101 | A | C | 0.228432 | -0.32307 | 0.056874 | 1.34E-08 | 0.000118 |
| 19 | rs56129017 | 41416948 | T | C | 0.372814 | 0.264515 | 0.046566 | 1.34E-08 | 0.000118 |
| 19 | rs2161245 | 41421517 | A | G | 0.372853 | 0.264512 | 0.046573 | 1.35E-08 | 0.000119 |
| 19 | rs4001944 | 41425875 | A | C | 0.37297 | 0.26453 | 0.046577 | 1.35E-08 | 0.000119 |
| 19 | rs76935404 | 41419294 | T | C | 0.372789 | 0.264145 | 0.046563 | 1.40E-08 | 0.000123 |
| 19 | rs3933911 | 41428583 | G | C | 0.261568 | -0.29356 | 0.051748 | 1.40E-08 | 0.000123 |
| 19 | rs7249124 | 41437619 | T | C | 0.261561 | -0.29353 | 0.051738 | 1.40E-08 | 0.000123 |
| 19 | rs12459565 | 41427539 | G | C | 0.261573 | -0.29348 | 0.051749 | 1.42E-08 | 0.000124 |
| 15 | rs11636545 | 79140348 | T | C | 0.391177 | -0.27472 | 0.048454 | 1.43E-08 | 0.000124 |
| 19 | rs11083590 | 41428626 | C | T | 0.26157 | -0.29341 | 0.051748 | 1.43E-08 | 0.000124 |
| 19 | rs73034465 | 41419075 | G | A | 0.372811 | 0.2639 | 0.046564 | 1.45E-08 | 0.000125 |
| 19 | rs4468739 | 41424762 | C | T | 0.372901 | 0.263968 | 0.046576 | 1.45E-08 | 0.000125 |
| 19 | rs1820025 | 41436546 | G | C | 0.261695 | -0.29293 | 0.051736 | 1.50E-08 | 0.000129 |
| 15 | rs2568500 | 78726928 | T | C | 0.454654 | -0.26994 | 0.047686 | 1.51E-08 | 0.00013 |
| 15 | rs12909921 | 78743260 | G | A | 0.454131 | -0.26958 | 0.047659 | 1.54E-08 | 0.000133 |
| 15 | rs10519198 | 78742754 | A | C | 0.454124 | -0.26939 | 0.047659 | 1.58E-08 | 0.000136 |

| | | | | | | | | | |
|---|---|---|---|---|---|---|---|---|---|
| 19 | rs4090553 | 41426757 | T | C | 0.261702 | -0.29234 | 0.051745 | 1.61E-08 | 0.000138 |
| 19 | rs4001943 | 41425900 | T | C | 0.262261 | -0.29219 | 0.051724 | 1.61E-08 | 0.000138 |
| 8 | rs6990603 | 42523039 | T | G | 0.188779 | -0.34847 | 0.06175 | 1.67E-08 | 0.000142 |
| 15 | rs7402877 | 79132814 | C | T | 0.391577 | -0.27341 | 0.048447 | 1.67E-08 | 0.000142 |
| 19 | rs4001945 | 41425721 | T | C | 0.259497 | -0.29326 | 0.051971 | 1.67E-08 | 0.000142 |
| 15 | rs6495341 | 79133026 | T | C | 0.39157 | -0.27283 | 0.048436 | 1.77E-08 | 0.000151 |
| 15 | rs12899131 | 78726885 | G | A | 0.45403 | -0.26846 | 0.047694 | 1.81E-08 | 0.000154 |
| 15 | rs2036532 | 78727386 | A | G | 0.454008 | -0.26842 | 0.047697 | 1.83E-08 | 0.000155 |
| 8 | rs11995030 | 42626418 | G | A | 0.201411 | -0.33707 | 0.059902 | 1.83E-08 | 0.000155 |
| 8 | rs10109040 | 42625313 | G | C | 0.201427 | -0.33665 | 0.059901 | 1.91E-08 | 0.000161 |
| 20 | rs2273500 | 61986949 | C | T | 0.14741 | 0.350165 | 0.06245 | 2.06E-08 | 0.000173 |
| 8 | rs10108797 | 42625074 | G | C | 0.201343 | -0.33569 | 0.059905 | 2.10E-08 | 0.000176 |
| 15 | rs954144 | 78730591 | T | C | 0.454113 | -0.26705 | 0.047695 | 2.15E-08 | 0.000181 |
| 15 | rs11072765 | 78734292 | G | A | 0.453906 | -0.26683 | 0.047695 | 2.21E-08 | 0.000185 |
| 15 | rs1847529 | 78735070 | C | A | 0.454281 | -0.26635 | 0.047688 | 2.33E-08 | 0.000195 |
| 15 | rs8041628 | 78735355 | G | C | 0.454245 | -0.26632 | 0.047688 | 2.34E-08 | 0.000196 |
| 8 | rs7837296 | 42526894 | A | C | 0.189119 | -0.34417 | 0.061673 | 2.40E-08 | 0.0002 |
| 19 | rs3852872 | 41416143 | C | T | 0.260774 | -0.29011 | 0.05202 | 2.45E-08 | 0.000204 |
| 8 | rs36057318 | 42525284 | T | G | 0.188962 | -0.344 | 0.061695 | 2.46E-08 | 0.000205 |
| 19 | rs10420231 | 41420030 | G | A | 0.260892 | -0.28964 | 0.051975 | 2.51E-08 | 0.000208 |
| 15 | rs7181033 | 79152414 | T | C | 0.384878 | -0.27034 | 0.048568 | 2.60E-08 | 0.000215 |
| 19 | rs191883259 | 41275725 | T | C | 0.025132 | -0.8066 | 0.145017 | 2.67E-08 | 0.00022 |
| 19 | rs142449117 | 41287573 | G | C | 0.025032 | -0.8085 | 0.145355 | 2.66E-08 | 0.00022 |
| 19 | rs3852873 | 41416260 | T | G | 0.260687 | -0.28933 | 0.052023 | 2.67E-08 | 0.00022 |
| 19 | rs10419450 | 41419925 | A | G | 0.260875 | -0.28886 | 0.051976 | 2.74E-08 | 0.000225 |
| 8 | rs4737068 | 42626424 | T | C | 0.20112 | -0.33258 | 0.059949 | 2.90E-08 | 0.000237 |
| 15 | rs8043123 | 78973393 | T | C | 0.229805 | -0.316 | 0.056981 | 2.93E-08 | 0.00024 |
| 8 | rs4737069 | 42626560 | G | A | 0.201042 | -0.33206 | 0.059952 | 3.05E-08 | 0.000249 |
| 15 | rs8032552 | 78971136 | C | T | 0.230489 | -0.3152 | 0.056922 | 3.07E-08 | 0.00025 |
| 19 | rs4500872 | 41422371 | C | T | 0.261069 | -0.28741 | 0.051909 | 3.08E-08 | 0.000251 |
| 8 | rs7845663 | 42608563 | A | G | 0.186084 | -0.33636 | 0.060787 | 3.14E-08 | 0.000255 |
| 8 | rs28501554 | 42622160 | G | T | 0.201251 | -0.33152 | 0.059912 | 3.14E-08 | 0.000255 |

| | | | | | | | | | |
|---|---|---|---|---|---|---|---|---|---|
| 8 | rs2117225 | 42627113 | A | G | 0.200934 | -0.33071 | 0.05995 | 3.46E-08 | 0.00028 |
| 15 | rs56227704 | 79136372 | G | A | 0.427077 | 0.269792 | 0.04892 | 3.49E-08 | 0.000282 |
| 15 | rs7180375 | 79157379 | C | T | 0.384888 | -0.26766 | 0.048562 | 3.55E-08 | 0.000287 |
| 8 | rs5005909 | 42528667 | G | A | 0.189378 | -0.33866 | 0.061751 | 4.15E-08 | 0.000335 |
| 19 | rs139707534 | 41273971 | C | T | 0.025166 | -0.79419 | 0.144949 | 4.28E-08 | 0.000344 |
| 19 | rs144374087 | 41271825 | C | G | 0.025065 | -0.79482 | 0.145237 | 4.44E-08 | 0.000354 |
| 19 | rs200795716 | 41273444 | G | A | 0.025066 | -0.7947 | 0.145237 | 4.46E-08 | 0.000354 |
| 19 | rs144243973 | 41273622 | A | G | 0.025066 | -0.7947 | 0.145237 | 4.46E-08 | 0.000354 |
| 19 | rs116972110 | 41276055 | T | G | 0.025066 | -0.7947 | 0.145237 | 4.46E-08 | 0.000354 |
| 19 | rs146641468 | 41278493 | C | A | 0.025066 | -0.7947 | 0.145237 | 4.46E-08 | 0.000354 |
| 19 | rs141788519 | 41279142 | T | C | 0.025066 | -0.7947 | 0.145237 | 4.46E-08 | 0.000354 |
| 19 | rs146221490 | 41280048 | C | G | 0.025066 | -0.7947 | 0.145237 | 4.46E-08 | 0.000354 |
| 19 | rs2233149 | 41280550 | G | T | 0.025066 | -0.7947 | 0.145237 | 4.46E-08 | 0.000354 |
| 19 | rs141715458 | 41284397 | C | T | 0.025084 | -0.79362 | 0.14521 | 4.62E-08 | 0.000365 |
| 19 | rs146247868 | 41284546 | A | G | 0.025084 | -0.79362 | 0.14521 | 4.62E-08 | 0.000365 |
| 19 | rs144761778 | 41266842 | C | T | 0.02506 | -0.79349 | 0.14525 | 4.68E-08 | 0.00037 |
| 8 | rs10087172 | 42616868 | C | T | 0.201424 | -0.32703 | 0.059891 | 4.75E-08 | 0.000375 |
| 8 | rs10110332 | 42613186 | A | C | 0.201266 | -0.32667 | 0.059899 | 4.94E-08 | 0.000388 |
| 8 | rs892413 | 42614378 | A | C | 0.201149 | -0.32661 | 0.0599 | 4.96E-08 | 0.00039 |
| 19 | rs12971445 | 41484602 | T | G | 0.333295 | -0.26326 | 0.048287 | 4.98E-08 | 0.000391 |
| 15 | rs6495346 | 79157755 | C | T | 0.385407 | -0.26428 | 0.048547 | 5.22E-08 | 0.000408 |
| 19 | rs111390768 | 41484257 | G | C | 0.333361 | -0.26108 | 0.048296 | 6.45E-08 | 0.000504 |
| 15 | rs28661610 | 78985317 | A | G | 0.221711 | -0.31159 | 0.057901 | 7.39E-08 | 0.000577 |
| 8 | rs2217732 | 42618446 | G | A | 0.207567 | -0.31982 | 0.059436 | 7.41E-08 | 0.000577 |
| 8 | rs7824614 | 42607123 | G | A | 0.206817 | -0.32019 | 0.059583 | 7.71E-08 | 0.0006 |
| 8 | rs7828366 | 42607107 | C | T | 0.20683 | -0.31928 | 0.059584 | 8.39E-08 | 0.000651 |
| 15 | rs11629639 | 79019086 | A | G | 0.237073 | -0.30621 | 0.057176 | 8.53E-08 | 0.000661 |
| 8 | rs7824155 | 42606851 | G | A | 0.206989 | -0.3162 | 0.059539 | 1.09E-07 | 0.000844 |
| 8 | rs10808962 | 42606649 | C | T | 0.206973 | -0.31561 | 0.059549 | 1.16E-07 | 0.000894 |
| 15 | rs28455815 | 79114453 | C | T | 0.427979 | 0.260498 | 0.049266 | 1.24E-07 | 0.000955 |
| 15 | rs899997 | 79019578 | G | T | 0.23727 | -0.30162 | 0.057145 | 1.30E-07 | 0.001004 |
| 19 | rs113557544 | 41294910 | T | C | 0.026169 | -0.74686 | 0.141632 | 1.34E-07 | 0.00103 |

| | | | | | | | | | |
|---|---|---|---|---|---|---|---|---|---|
| 19 | rs7251781 | 41301069 | G | A | 0.026266 | -0.74363 | 0.141296 | 1.42E-07 | 0.001088 |
| 19 | rs7257854 | 41492677 | C | T | 0.335953 | -0.25367 | 0.048207 | 1.42E-07 | 0.00109 |
| 19 | rs149755483 | 41299866 | C | A | 0.026232 | -0.74391 | 0.14142 | 1.44E-07 | 0.0011 |
| 19 | rs892216 | 41489851 | C | T | 0.33627 | -0.25335 | 0.048208 | 1.48E-07 | 0.001128 |
| 15 | rs11633170 | 79004642 | C | T | 0.237045 | -0.30037 | 0.057201 | 1.51E-07 | 0.001152 |
| 15 | rs11629824 | 79005524 | T | G | 0.237107 | -0.30011 | 0.057193 | 1.54E-07 | 0.001174 |
| 15 | rs11072793 | 79006442 | G | A | 0.23715 | -0.30009 | 0.057193 | 1.55E-07 | 0.001174 |
| 15 | rs1809412 | 79021908 | C | T | 0.237175 | -0.29983 | 0.05715 | 1.55E-07 | 0.001177 |
| 8 | rs78145231 | 42646264 | T | C | 0.140422 | -0.35273 | 0.067265 | 1.57E-07 | 0.00119 |
| 8 | rs9298628 | 42605991 | T | C | 0.206625 | -0.31226 | 0.05958 | 1.60E-07 | 0.001207 |
| 19 | rs149397079 | 41302140 | C | A | 0.026227 | -0.74095 | 0.141431 | 1.62E-07 | 0.001219 |
| 19 | rs45600135 | 41302501 | A | G | 0.026227 | -0.74079 | 0.141431 | 1.62E-07 | 0.001224 |
| 8 | rs1960346 | 42643045 | T | C | 0.23366 | -0.29252 | 0.055875 | 1.65E-07 | 0.001239 |
| 15 | rs11634628 | 79005579 | G | A | 0.237142 | -0.29921 | 0.057191 | 1.68E-07 | 0.001262 |
| 8 | rs7004108 | 42608710 | G | A | 0.207069 | -0.31138 | 0.059521 | 1.68E-07 | 0.001262 |
| 15 | rs9888691 | 79010046 | T | C | 0.23716 | -0.29899 | 0.057186 | 1.71E-07 | 0.001281 |
| 19 | rs55721612 | 41486375 | C | T | 0.336524 | -0.25198 | 0.048213 | 1.73E-07 | 0.001292 |
| 19 | rs113336186 | 41296479 | T | G | 0.026122 | -0.74049 | 0.141757 | 1.75E-07 | 0.001309 |
| 15 | rs4887078 | 79011073 | C | T | 0.237195 | -0.29849 | 0.057188 | 1.80E-07 | 0.001338 |
| 15 | rs7497235 | 79014517 | G | A | 0.236807 | -0.29856 | 0.057222 | 1.81E-07 | 0.001349 |
| 8 | rs7008077 | 42644888 | G | A | 0.140673 | -0.35033 | 0.067196 | 1.85E-07 | 0.001376 |
| 15 | rs11072794 | 79006582 | T | C | 0.237149 | -0.29808 | 0.057191 | 1.87E-07 | 0.001385 |
| 15 | rs8039034 | 79017861 | C | T | 0.237063 | -0.29808 | 0.057212 | 1.89E-07 | 0.001398 |
| 15 | rs11635680 | 79016547 | G | A | 0.236856 | -0.29814 | 0.05724 | 1.90E-07 | 0.001407 |
| 8 | rs28826992 | 42641577 | C | T | 0.233845 | -0.29061 | 0.055855 | 1.96E-07 | 0.001448 |
| 15 | rs75810806 | 79015329 | T | C | 0.236895 | -0.29783 | 0.057247 | 1.97E-07 | 0.001449 |
| 15 | rs12899940 | 79001699 | C | T | 0.236265 | -0.2977 | 0.057251 | 1.99E-07 | 0.001468 |
| 8 | rs28784665 | 42641195 | A | G | 0.233739 | -0.29001 | 0.055857 | 2.08E-07 | 0.001528 |
| 15 | rs7497171 | 79014509 | C | A | 0.236717 | -0.2969 | 0.057263 | 2.16E-07 | 0.001586 |
| 8 | rs9298629 | 42606186 | T | G | 0.206642 | -0.30875 | 0.059572 | 2.19E-07 | 0.0016 |
| 15 | rs12916717 | 79014948 | T | C | 0.236824 | -0.29656 | 0.057251 | 2.22E-07 | 0.001622 |
| 20 | rs6011779 | 61984317 | C | T | 0.194248 | 0.288923 | 0.055882 | 2.34E-07 | 0.001707 |

| | | | | | | | | | |
|---|---|---|---|---|---|---|---|---|---|
| 8 | rs7812298 | 42608579 | T | C | 0.206833 | -0.30757 | 0.059523 | 2.38E-07 | 0.001732 |
| 8 | rs56395510 | 42642160 | G | A | 0.233844 | -0.2883 | 0.055889 | 2.49E-07 | 0.001811 |
| 8 | rs7843218 | 42640451 | C | T | 0.140825 | -0.34601 | 0.067112 | 2.53E-07 | 0.001835 |
| 8 | rs11786036 | 42642808 | C | T | 0.140765 | -0.34548 | 0.067157 | 2.68E-07 | 0.001947 |
| 15 | rs117756166 | 78939395 | C | T | 0.050493 | 0.517711 | 0.100782 | 2.79E-07 | 0.002019 |
| 19 | rs150437622 | 41013214 | A | C | 0.029993 | -0.65442 | 0.127393 | 2.79E-07 | 0.002019 |
| 8 | rs11990422 | 42582955 | A | G | 0.10622 | -0.38813 | 0.075603 | 2.84E-07 | 0.00205 |
| 15 | rs28505515 | 79114477 | C | G | 0.466244 | 0.244706 | 0.047787 | 3.04E-07 | 0.002193 |
| 15 | rs1809414 | 79028120 | G | T | 0.236729 | -0.29316 | 0.05726 | 3.06E-07 | 0.002202 |
| 15 | rs11072799 | 79027814 | G | T | 0.236726 | -0.29304 | 0.05726 | 3.09E-07 | 0.002223 |
| 8 | rs11776376 | 42649863 | A | G | 0.1398 | -0.34515 | 0.067468 | 3.13E-07 | 0.002243 |
| 8 | rs71521595 | 42573091 | C | T | 0.106317 | -0.38646 | 0.075563 | 3.15E-07 | 0.002254 |
| 8 | rs112866926 | 42633597 | T | C | 0.140637 | -0.34317 | 0.06715 | 3.21E-07 | 0.002295 |
| 19 | rs2303728 | 41114065 | T | G | 0.195997 | -0.28786 | 0.056328 | 3.21E-07 | 0.002295 |
| 15 | rs11638020 | 79107724 | A | G | 0.405029 | 0.253276 | 0.049604 | 3.29E-07 | 0.002347 |
| 19 | rs4803361 | 41241975 | G | T | 0.285913 | -0.25365 | 0.049714 | 3.36E-07 | 0.002391 |
| 15 | rs12900519 | 78949127 | C | T | 0.062706 | 0.466018 | 0.091468 | 3.49E-07 | 0.002481 |
| 15 | rs11856441 | 79030526 | T | G | 0.236354 | -0.29178 | 0.057285 | 3.52E-07 | 0.002496 |
| 15 | rs56683682 | 79112524 | T | G | 0.406037 | 0.251538 | 0.049512 | 3.77E-07 | 0.00267 |
| 8 | rs7828365 | 42629314 | T | C | 0.1406 | -0.34056 | 0.067134 | 3.92E-07 | 0.002775 |
| 15 | rs66925868 | 79115406 | G | A | 0.405366 | 0.250895 | 0.049508 | 4.02E-07 | 0.002844 |
| 19 | rs2607420 | 41244887 | G | A | 0.286648 | -0.25169 | 0.049702 | 4.11E-07 | 0.002898 |
| 8 | rs2196129 | 42618406 | C | G | 0.140483 | -0.33919 | 0.067119 | 4.34E-07 | 0.003056 |
| 15 | rs3971829 | 79109052 | A | G | 0.405835 | 0.250013 | 0.049489 | 4.37E-07 | 0.003077 |
| 15 | rs12593043 | 78953346 | T | A | 0.062619 | 0.461881 | 0.09148 | 4.44E-07 | 0.00312 |
| 15 | rs4887075 | 78953131 | C | T | 0.062621 | 0.461739 | 0.091481 | 4.48E-07 | 0.003141 |
| 15 | rs12898292 | 79112603 | A | G | 0.407335 | 0.24945 | 0.049539 | 4.77E-07 | 0.003338 |
| 8 | rs13261190 | 42578309 | G | A | 0.106057 | -0.38038 | 0.075673 | 4.99E-07 | 0.003485 |
| 15 | rs12898712 | 79112622 | G | T | 0.407117 | 0.249054 | 0.049544 | 4.99E-07 | 0.003485 |
| 15 | rs56076311 | 79115120 | G | A | 0.408985 | 0.248597 | 0.049469 | 5.03E-07 | 0.003504 |
| 15 | rs12899452 | 79112550 | T | A | 0.406552 | 0.24816 | 0.049511 | 5.38E-07 | 0.003746 |
| 15 | rs755998 | 79032469 | C | T | 0.238478 | -0.28579 | 0.057055 | 5.47E-07 | 0.003803 |

| | | | | | | | | | |
|---|---|---|---|---|---|---|---|---|---|
| 15 | rs3890715 | 79107862 | C | A | 0.406106 | 0.248241 | 0.049575 | 5.52E-07 | 0.003828 |
| 15 | rs12898452 | 79112628 | C | A | 0.407117 | 0.247929 | 0.049542 | 5.60E-07 | 0.003884 |
| 15 | rs7403608 | 79115570 | G | A | 0.405598 | 0.247654 | 0.049497 | 5.63E-07 | 0.003897 |
| 15 | rs7167242 | 79113541 | G | A | 0.405823 | 0.247622 | 0.049496 | 5.65E-07 | 0.003902 |
| 15 | rs7167056 | 79113484 | G | T | 0.405884 | 0.2475 | 0.049493 | 5.71E-07 | 0.00394 |
| 15 | rs34372090 | 79110735 | T | C | 0.405407 | 0.247063 | 0.049471 | 5.91E-07 | 0.004065 |
| 15 | rs55870734 | 79117917 | T | A | 0.405572 | 0.247613 | 0.049578 | 5.90E-07 | 0.004065 |
| 15 | rs28437201 | 57049810 | T | C | 0.163424 | 0.341376 | 0.068376 | 5.96E-07 | 0.004091 |
| 19 | rs3852875 | 41449922 | T | C | 0.262229 | -0.25461 | 0.051038 | 6.08E-07 | 0.004169 |
| 15 | rs7178051 | 79118296 | T | C | 0.405509 | 0.247347 | 0.049587 | 6.09E-07 | 0.004174 |
| 15 | rs7166340 | 79113293 | C | T | 0.405879 | 0.246793 | 0.049492 | 6.15E-07 | 0.004198 |
| 15 | rs7167853 | 79113318 | T | C | 0.405879 | 0.246793 | 0.049492 | 6.15E-07 | 0.004198 |
| 15 | rs12909317 | 79114419 | T | G | 0.405627 | 0.246748 | 0.049494 | 6.18E-07 | 0.004216 |
| 15 | rs7402376 | 79113570 | G | A | 0.405621 | 0.246677 | 0.04949 | 6.22E-07 | 0.004233 |
| 15 | rs4887092 | 79043675 | T | C | 0.237922 | -0.28433 | 0.057057 | 6.25E-07 | 0.004244 |
| 15 | rs6495333 | 79113215 | G | A | 0.405881 | 0.246632 | 0.049491 | 6.25E-07 | 0.004244 |
| 15 | rs12915192 | 79115585 | G | A | 0.40602 | 0.246622 | 0.049499 | 6.28E-07 | 0.004259 |
| 15 | rs7166770 | 79113568 | A | G | 0.405843 | 0.246572 | 0.049495 | 6.30E-07 | 0.004266 |
| 15 | rs2622811 | 79034480 | T | A | 0.239259 | -0.28431 | 0.057076 | 6.32E-07 | 0.00427 |
| 9 | rs56116178 | 1.36E+08 | G | A | 0.104395 | 0.35648 | 0.071577 | 6.35E-07 | 0.004284 |
| 15 | rs9806363 | 79110302 | G | A | 0.405568 | 0.246189 | 0.049465 | 6.46E-07 | 0.004352 |
| 8 | rs76874443 | 42609433 | T | C | 0.140022 | -0.33456 | 0.067257 | 6.55E-07 | 0.004407 |
| 15 | rs2017091 | 79032744 | G | A | 0.238485 | -0.28339 | 0.057052 | 6.79E-07 | 0.004562 |
| 15 | rs12913779 | 79032355 | T | C | 0.238473 | -0.2835 | 0.057082 | 6.82E-07 | 0.004569 |
| 15 | rs12442476 | 79117856 | G | A | 0.405863 | 0.246208 | 0.049574 | 6.82E-07 | 0.004569 |
| 15 | rs11072807 | 79111517 | A | G | 0.405605 | 0.245622 | 0.049468 | 6.86E-07 | 0.004589 |
| 15 | rs7403593 | 79111962 | G | A | 0.405553 | 0.245587 | 0.049466 | 6.88E-07 | 0.004589 |
| 15 | rs7403728 | 79112016 | G | T | 0.405553 | 0.245587 | 0.049466 | 6.88E-07 | 0.004589 |
| 8 | rs57308096 | 42537736 | A | G | 0.122354 | -0.35422 | 0.071369 | 6.94E-07 | 0.004616 |
| 15 | rs8027833 | 79112837 | T | C | 0.405949 | 0.245593 | 0.049487 | 6.95E-07 | 0.004616 |
| 15 | rs8026598 | 79112938 | A | G | 0.405964 | 0.245596 | 0.049485 | 6.94E-07 | 0.004616 |
| 8 | rs16891620 | 42625663 | A | C | 0.140316 | -0.33346 | 0.0672 | 6.97E-07 | 0.004623 |

| | | | | | | | | | |
|---|---|---|---|---|---|---|---|---|---|
| 19 | rs12982314 | 41385774 | A | G | 0.255299 | -0.25497 | 0.051385 | 6.98E-07 | 0.004623 |
| 15 | rs3878226 | 79034412 | T | C | 0.239078 | -0.28284 | 0.05703 | 7.07E-07 | 0.004662 |
| 15 | rs7165902 | 79115872 | T | G | 0.405911 | 0.245496 | 0.0495 | 7.07E-07 | 0.004662 |
| 15 | rs7165075 | 79115965 | G | A | 0.40591 | 0.245493 | 0.0495 | 7.07E-07 | 0.004662 |
| 8 | rs13277840 | 42530999 | T | G | 0.106529 | -0.37423 | 0.075471 | 7.10E-07 | 0.004676 |
| 15 | rs12906847 | 79031357 | A | G | 0.238254 | -0.2835 | 0.057177 | 7.11E-07 | 0.004679 |
| 15 | rs4387568 | 79109514 | G | C | 0.405673 | 0.245293 | 0.049475 | 7.12E-07 | 0.004679 |
| 15 | rs12907207 | 79031290 | G | A | 0.237828 | -0.28334 | 0.057187 | 7.25E-07 | 0.004754 |
| 15 | rs12898392 | 79112432 | G | A | 0.405798 | 0.244959 | 0.049476 | 7.38E-07 | 0.004834 |
| 19 | rs4079366 | 41384675 | T | C | 0.255447 | -0.25398 | 0.051348 | 7.57E-07 | 0.004949 |
| 15 | rs12906977 | 79031329 | C | A | 0.238347 | -0.28271 | 0.057172 | 7.62E-07 | 0.004962 |
| 15 | rs9920493 | 79113846 | A | C | 0.405961 | 0.24474 | 0.049493 | 7.62E-07 | 0.004962 |
| 15 | rs8041064 | 79115388 | A | G | 0.405987 | 0.244772 | 0.049497 | 7.61E-07 | 0.004962 |
| 19 | rs2302988 | 41383344 | T | C | 0.255507 | -0.25408 | 0.051418 | 7.76E-07 | 0.005045 |
| 15 | rs12907432 | 79031473 | G | C | 0.237734 | -0.28247 | 0.057188 | 7.84E-07 | 0.005085 |
| 15 | rs12907536 | 79031485 | G | A | 0.237734 | -0.28247 | 0.057188 | 7.84E-07 | 0.005085 |
| 19 | rs10402703 | 41384386 | C | G | 0.255946 | -0.25346 | 0.051319 | 7.85E-07 | 0.005087 |
| 15 | rs1383635 | 79029511 | T | C | 0.238186 | -0.28205 | 0.057134 | 7.95E-07 | 0.005139 |
| 15 | rs2219939 | 79029723 | G | A | 0.238189 | -0.28203 | 0.057132 | 7.96E-07 | 0.005139 |
| 19 | rs2909906 | 41384084 | T | C | 0.25591 | -0.25341 | 0.051338 | 7.97E-07 | 0.005142 |
| 15 | rs12901228 | 79030049 | G | T | 0.238195 | -0.28201 | 0.057143 | 8.01E-07 | 0.005159 |
| 15 | rs28513584 | 79109856 | G | A | 0.406007 | 0.24403 | 0.049459 | 8.06E-07 | 0.005181 |
| 15 | rs1383634 | 79029396 | T | C | 0.238169 | -0.28173 | 0.057135 | 8.18E-07 | 0.005242 |
| 15 | rs1825086 | 79029492 | A | G | 0.238169 | -0.28173 | 0.057135 | 8.18E-07 | 0.005242 |
| 15 | rs1904920 | 79029587 | C | T | 0.238169 | -0.28173 | 0.057135 | 8.18E-07 | 0.005242 |
| 19 | rs2545781 | 41385768 | T | C | 0.255443 | -0.25322 | 0.051376 | 8.28E-07 | 0.005296 |
| 19 | rs10404300 | 41384385 | T | G | 0.255949 | -0.25287 | 0.05132 | 8.33E-07 | 0.005323 |
| 15 | rs7403714 | 79118650 | T | C | 0.405347 | 0.243723 | 0.049608 | 8.97E-07 | 0.005722 |
| 15 | rs6495335 | 79117133 | G | T | 0.405868 | 0.242832 | 0.049548 | 9.54E-07 | 0.006077 |
| 15 | rs112723723 | 79031139 | G | C | 0.237957 | -0.28021 | 0.057276 | 9.97E-07 | 0.006342 |
| 15 | rs7173137 | 57026346 | A | C | 0.163537 | 0.334324 | 0.068381 | 1.01E-06 | 0.006435 |
| 7 | rs1635759 | 74339397 | G | A | 0.188994 | 0.273723 | 0.05602 | 1.03E-06 | 0.006524 |

| | | | | | | | | | |
|---|---|---|---|---|---|---|---|---|---|
| 15 | rs71402966 | 79031059 | G | A | 0.237977 | -0.2798 | 0.057276 | 1.03E-06 | 0.006549 |
| 15 | rs11071290 | 57051751 | G | T | 0.16475 | 0.332789 | 0.068173 | 1.05E-06 | 0.00666 |
| 15 | rs12900336 | 78948850 | G | A | 0.063516 | 0.443525 | 0.090938 | 1.08E-06 | 0.006798 |
| 15 | rs58021857 | 79119229 | A | C | 0.405469 | 0.241906 | 0.049613 | 1.08E-06 | 0.006837 |
| 4 | rs4282179 | 1.05E+08 | T | C | 0.350472 | 0.232045 | 0.047621 | 1.10E-06 | 0.006936 |
| 4 | rs4440219 | 1.05E+08 | T | C | 0.348944 | 0.232798 | 0.047784 | 1.11E-06 | 0.006957 |
| 19 | rs2604909 | 41297615 | T | C | 0.207757 | 0.26906 | 0.055253 | 1.12E-06 | 0.007029 |
| 15 | rs8030090 | 79034260 | A | G | 0.240794 | -0.27656 | 0.056849 | 1.15E-06 | 0.007188 |
| 15 | rs6495312 | 78959322 | T | C | 0.062728 | 0.444681 | 0.091471 | 1.17E-06 | 0.007304 |
| 4 | rs4475153 | 1.05E+08 | T | C | 0.350431 | 0.231228 | 0.047626 | 1.20E-06 | 0.007533 |
| 20 | rs45577732 | 61983934 | G | C | 0.081321 | 0.394064 | 0.081175 | 1.21E-06 | 0.007545 |
| 15 | rs12904489 | 79033925 | T | A | 0.240825 | -0.27584 | 0.05684 | 1.22E-06 | 0.007597 |
| 15 | rs11633152 | 79032256 | C | T | 0.238091 | -0.27751 | 0.057192 | 1.22E-06 | 0.007613 |
| 15 | rs8041389 | 79115491 | C | T | 0.405482 | 0.240476 | 0.049588 | 1.24E-06 | 0.007707 |
| 3 | rs75741814 | 1.06E+08 | G | A | 0.135106 | -0.34419 | 0.07098 | 1.24E-06 | 0.007712 |
| 15 | rs8030335 | 79034124 | G | A | 0.240786 | -0.27548 | 0.056846 | 1.26E-06 | 0.007821 |
| 19 | rs2604910 | 41297254 | T | C | 0.207737 | 0.267748 | 0.055259 | 1.26E-06 | 0.007839 |
| 15 | rs11856432 | 57046107 | G | A | 0.161339 | 0.33277 | 0.068802 | 1.32E-06 | 0.008178 |
| 1 | rs35252813 | 32256590 | A | G | 0.034277 | 0.582827 | 0.120704 | 1.38E-06 | 0.008494 |
| 4 | rs10018248 | 1.05E+08 | T | C | 0.357365 | 0.229517 | 0.047531 | 1.37E-06 | 0.008494 |
| 19 | rs2604911 | 41296117 | G | C | 0.207909 | 0.266704 | 0.055259 | 1.39E-06 | 0.008574 |
| 3 | rs9872827 | 1.06E+08 | T | C | 0.13607 | -0.34115 | 0.070768 | 1.43E-06 | 0.008813 |
| 15 | rs12899201 | 79033534 | G | A | 0.240589 | -0.27387 | 0.056858 | 1.46E-06 | 0.008982 |
| 20 | rs144298540 | 61984931 | T | G | 0.082957 | 0.386981 | 0.080381 | 1.48E-06 | 0.009074 |
| 15 | rs4243085 | 79057723 | G | C | 0.234023 | -0.27343 | 0.057072 | 1.66E-06 | 0.010183 |
| 8 | rs34626112 | 42525628 | T | G | 0.121835 | -0.34244 | 0.071539 | 1.69E-06 | 0.010384 |
| 19 | rs62107913 | 41127036 | T | C | 0.171658 | -0.28336 | 0.059202 | 1.70E-06 | 0.010393 |
| 19 | rs2604874 | 41257816 | C | A | 0.278568 | -0.23988 | 0.050137 | 1.71E-06 | 0.010472 |
| 8 | rs11785369 | 42527386 | C | A | 0.121932 | -0.34164 | 0.071508 | 1.77E-06 | 0.010823 |
| 20 | rs45623037 | 61989658 | C | G | 0.082254 | 0.385959 | 0.080801 | 1.78E-06 | 0.010863 |
| 15 | rs4887091 | 79043580 | T | C | 0.240234 | -0.27155 | 0.056877 | 1.80E-06 | 0.010978 |
| 8 | rs6999449 | 42638261 | C | A | 0.215002 | -0.27567 | 0.057748 | 1.81E-06 | 0.011 |

| | | | | | | | | | |
|---|---|---|---|---|---|---|---|---|---|
| 4 | rs28564630 | 1.05E+08 | A | T | 0.351432 | 0.227426 | 0.047659 | 1.82E-06 | 0.011076 |
| 20 | rs45449494 | 61987930 | G | A | 0.082434 | 0.384863 | 0.080669 | 1.83E-06 | 0.011121 |
| 15 | rs12148817 | 79041128 | A | G | 0.2404 | -0.27133 | 0.056885 | 1.84E-06 | 0.011168 |
| 19 | rs7247231 | 41339837 | A | T | 0.131482 | -0.31213 | 0.065493 | 1.88E-06 | 0.011378 |
| 8 | rs7825261 | 42637683 | C | A | 0.214889 | -0.27501 | 0.057777 | 1.94E-06 | 0.011702 |
| 3 | rs17807129 | 1.06E+08 | A | G | 0.135651 | -0.3369 | 0.070887 | 2.01E-06 | 0.012115 |
| 19 | rs55740667 | 41112605 | C | G | 0.172472 | -0.27988 | 0.058945 | 2.05E-06 | 0.012372 |
| 15 | rs12910831 | 79043801 | T | C | 0.239843 | -0.26998 | 0.056907 | 2.09E-06 | 0.012594 |
| 19 | rs9676346 | 41472269 | T | C | 0.388502 | 0.221932 | 0.046955 | 2.28E-06 | 0.013729 |
| 15 | rs1114829 | 78989821 | A | G | 0.211321 | -0.27825 | 0.058921 | 2.33E-06 | 0.013983 |
| 11 | rs483129 | 1.33E+08 | A | T | 0.012924 | 0.901588 | 0.191006 | 2.36E-06 | 0.014122 |
| 15 | rs2017297 | 79033278 | T | C | 0.240772 | -0.26803 | 0.056869 | 2.44E-06 | 0.0146 |
| 3 | rs75847184 | 1.06E+08 | A | G | 0.135833 | -0.33365 | 0.070822 | 2.46E-06 | 0.014728 |
| 15 | rs12908510 | 78997972 | T | C | 0.227983 | -0.27292 | 0.057956 | 2.49E-06 | 0.014855 |
| 19 | rs7254893 | 41252736 | C | G | 0.281716 | -0.23481 | 0.049883 | 2.51E-06 | 0.014974 |
| 19 | rs4802079 | 41114866 | A | C | 0.172992 | -0.27666 | 0.058863 | 2.60E-06 | 0.015482 |
| 15 | rs11632672 | 79044330 | A | G | 0.239849 | -0.2673 | 0.056901 | 2.63E-06 | 0.015656 |
| 15 | rs12905116 | 79042734 | T | A | 0.239587 | -0.26734 | 0.056914 | 2.64E-06 | 0.015659 |
| 15 | rs12909668 | 79043949 | A | G | 0.239882 | -0.26715 | 0.056898 | 2.66E-06 | 0.015803 |
| 15 | rs7182993 | 79045014 | G | A | 0.240059 | -0.26678 | 0.056901 | 2.75E-06 | 0.016302 |
| 15 | rs7170552 | 78994066 | C | A | 0.211064 | -0.27633 | 0.05897 | 2.79E-06 | 0.016495 |
| 15 | rs7182567 | 79045054 | A | G | 0.239976 | -0.26642 | 0.056887 | 2.82E-06 | 0.016679 |
| 15 | rs7170563 | 78994094 | G | A | 0.211062 | -0.27613 | 0.05897 | 2.83E-06 | 0.016721 |
| 19 | rs73047967 | 41113001 | A | G | 0.1694 | -0.2779 | 0.059368 | 2.85E-06 | 0.016828 |
| 15 | rs7182694 | 79045030 | C | T | 0.239969 | -0.26614 | 0.056887 | 2.89E-06 | 0.017022 |
| 19 | rs113892385 | 41126668 | A | G | 0.171178 | -0.27735 | 0.059357 | 2.97E-06 | 0.017492 |
| 14 | rs28786188 | 49386067 | T | G | 0.07245 | 0.400635 | 0.085807 | 3.03E-06 | 0.017774 |
| 19 | rs10425150 | 41387620 | A | G | 0.497143 | -0.21727 | 0.046546 | 3.04E-06 | 0.017842 |
| 15 | rs12917502 | 79045521 | T | C | 0.239931 | -0.26528 | 0.056864 | 3.08E-06 | 0.018063 |
| 8 | rs11782372 | 42532957 | G | C | 0.143715 | -0.3186 | 0.068317 | 3.11E-06 | 0.01818 |
| 19 | rs4802099 | 41454470 | A | G | 0.364498 | -0.22147 | 0.047513 | 3.14E-06 | 0.018359 |
| 7 | rs35005436 | 74134911 | C | T | 0.15622 | 0.284353 | 0.061022 | 3.16E-06 | 0.018466 |

| | | | | | | | | | |
|---|---|---|---|---|---|---|---|---|---|
| 15 | rs72625802 | 78992811 | A | G | 0.210957 | -0.27465 | 0.058969 | 3.20E-06 | 0.018654 |
| 16 | rs447735 | 89734349 | C | T | 0.420643 | 0.226414 | 0.048621 | 3.21E-06 | 0.018707 |
| 15 | rs6495317 | 78991716 | C | A | 0.211175 | -0.27418 | 0.058949 | 3.30E-06 | 0.019196 |
| 20 | rs4809543 | 61986950 | A | G | 0.079336 | 0.381337 | 0.082147 | 3.45E-06 | 0.020023 |
| 19 | rs2246761 | 41116001 | A | G | 0.169744 | -0.27537 | 0.059329 | 3.46E-06 | 0.020071 |
| 15 | rs2622821 | 79046693 | A | G | 0.239917 | -0.26396 | 0.05688 | 3.47E-06 | 0.020113 |
| 19 | rs7245595 | 41295859 | T | G | 0.342186 | 0.21696 | 0.046805 | 3.56E-06 | 0.020608 |
| 15 | rs12906835 | 79046989 | A | G | 0.239465 | -0.26351 | 0.056919 | 3.67E-06 | 0.021181 |
| 19 | rs41512448 | 41133844 | C | T | 0.171642 | -0.27348 | 0.059101 | 3.71E-06 | 0.021383 |
| 19 | rs10425169 | 41387647 | A | G | 0.497251 | -0.2153 | 0.046557 | 3.76E-06 | 0.021646 |
| 19 | rs11673582 | 41461318 | A | G | 0.364877 | -0.21946 | 0.047474 | 3.79E-06 | 0.021792 |
| 15 | rs16970006 | 78970259 | C | T | 0.067672 | -0.41116 | 0.089039 | 3.88E-06 | 0.022307 |
| 15 | rs7163236 | 78991292 | A | C | 0.210904 | -0.27195 | 0.058938 | 3.95E-06 | 0.022634 |
| 19 | rs12978065 | 41469279 | A | G | 0.364258 | -0.21935 | 0.047537 | 3.94E-06 | 0.022634 |
| 19 | rs35858034 | 41299864 | C | A | 0.342562 | 0.215355 | 0.046719 | 4.03E-06 | 0.02311 |
| 19 | rs10425176 | 41387656 | A | T | 0.497266 | -0.21443 | 0.046557 | 4.11E-06 | 0.023515 |
| 15 | rs12907739 | 78997980 | G | A | 0.229405 | -0.26612 | 0.057836 | 4.20E-06 | 0.023992 |
| 15 | rs6493886 | 57027079 | G | A | 0.165347 | 0.313325 | 0.068103 | 4.21E-06 | 0.024025 |
| 7 | rs35275911 | 74108249 | C | G | 0.193071 | 0.259018 | 0.056304 | 4.22E-06 | 0.024046 |
| 19 | rs4803419 | 41512792 | T | C | 0.296239 | -0.22688 | 0.049364 | 4.30E-06 | 0.0245 |
| 1 | rs10798888 | 32199099 | T | G | 0.174491 | 0.267265 | 0.058166 | 4.33E-06 | 0.024627 |
| 7 | rs68008267 | 74087465 | T | C | 0.192953 | 0.258626 | 0.05632 | 4.39E-06 | 0.024905 |
| 15 | rs2128171 | 56999901 | G | A | 0.165627 | 0.31002 | 0.067513 | 4.39E-06 | 0.024905 |
| 15 | rs2046213 | 56918975 | A | G | 0.201146 | 0.281273 | 0.061271 | 4.42E-06 | 0.025003 |
| 19 | rs2278243 | 41132022 | G | C | 0.172063 | -0.27069 | 0.058963 | 4.42E-06 | 0.025003 |
| 15 | rs7164095 | 57002275 | A | C | 0.164745 | 0.310323 | 0.067644 | 4.48E-06 | 0.025341 |
| 7 | rs67755137 | 74108135 | A | G | 0.193071 | 0.258187 | 0.056303 | 4.53E-06 | 0.025543 |
| 19 | rs4105143 | 41393533 | T | C | 0.499073 | -0.2136 | 0.046583 | 4.53E-06 | 0.025563 |
| 19 | rs7258590 | 41408581 | C | T | 0.497813 | -0.21375 | 0.046621 | 4.54E-06 | 0.025569 |
| 19 | rs10416968 | 41407637 | G | A | 0.497047 | -0.21301 | 0.046483 | 4.59E-06 | 0.025831 |
| 15 | rs11639335 | 79142307 | G | A | 0.411181 | 0.224211 | 0.048937 | 4.61E-06 | 0.025916 |
| 1 | rs12037019 | 32213655 | C | T | 0.042294 | 0.49868 | 0.108875 | 4.64E-06 | 0.026007 |

| | | | | | | | | | |
|---|---|---|---|---|---|---|---|---|---|
| 19 | rs12151282 | 41298419 | C | T | 0.342562 | 0.214022 | 0.046728 | 4.65E-06 | 0.026007 |
| 19 | rs7252852 | 41403325 | T | C | 0.497097 | -0.21291 | 0.046482 | 4.64E-06 | 0.026007 |
| 15 | rs2002854 | 79053014 | A | G | 0.240843 | -0.25904 | 0.056573 | 4.67E-06 | 0.026128 |
| 19 | rs2369006 | 41120083 | T | A | 0.305964 | -0.22208 | 0.048571 | 4.82E-06 | 0.026923 |
| 19 | rs2316319 | 41463215 | C | G | 0.364652 | -0.2172 | 0.047513 | 4.85E-06 | 0.027037 |
| 16 | rs6499240 | 69686912 | A | G | 0.404128 | -0.23439 | 0.051285 | 4.87E-06 | 0.027124 |
| 19 | rs57755489 | 41476740 | A | G | 0.364264 | -0.21722 | 0.047533 | 4.88E-06 | 0.027159 |
| 19 | rs59300404 | 41476707 | T | C | 0.364265 | -0.21714 | 0.047533 | 4.92E-06 | 0.027338 |
| 19 | rs882808 | 41479051 | A | G | 0.364396 | -0.21701 | 0.04751 | 4.93E-06 | 0.027374 |
| 4 | rs1974941 | 1.3E+08 | A | G | 0.244953 | 0.232608 | 0.050931 | 4.94E-06 | 0.027409 |
| 15 | rs12907511 | 78998061 | C | G | 0.228983 | -0.26372 | 0.057753 | 4.96E-06 | 0.027491 |
| 19 | rs1104854 | 41479309 | T | C | 0.364423 | -0.21691 | 0.047506 | 4.97E-06 | 0.0275 |
| 19 | rs67421541 | 41394420 | C | T | 0.498707 | -0.21257 | 0.046579 | 5.03E-06 | 0.027776 |
| 15 | rs12907178 | 78997922 | A | G | 0.22898 | -0.26353 | 0.057754 | 5.04E-06 | 0.02783 |
| 1 | rs12037400 | 32215178 | C | A | 0.042338 | 0.496596 | 0.108856 | 5.07E-06 | 0.027928 |
| 19 | rs7343061 | 41405348 | C | A | 0.497056 | -0.2119 | 0.046488 | 5.16E-06 | 0.028405 |
| 19 | rs11083582 | 41395851 | A | C | 0.498707 | -0.2123 | 0.046581 | 5.17E-06 | 0.028427 |
| 19 | rs28649715 | 41125850 | G | C | 0.305517 | -0.22185 | 0.048682 | 5.19E-06 | 0.028478 |
| 19 | rs3909342 | 41396690 | G | A | 0.498717 | -0.21216 | 0.046581 | 5.25E-06 | 0.028775 |
| 19 | rs10418318 | 41395036 | G | A | 0.498905 | -0.21198 | 0.046581 | 5.35E-06 | 0.029293 |
| 3 | rs12485274 | 1.06E+08 | A | G | 0.153677 | -0.30539 | 0.067138 | 5.40E-06 | 0.029549 |
| 7 | rs73137144 | 74073590 | G | A | 0.191721 | 0.257012 | 0.056507 | 5.41E-06 | 0.029557 |
| 19 | rs10421888 | 41392048 | C | T | 0.498678 | -0.21176 | 0.046572 | 5.45E-06 | 0.029735 |
| 1 | rs12035217 | 32212514 | C | T | 0.042623 | 0.492943 | 0.108463 | 5.50E-06 | 0.029987 |
| 19 | rs11665969 | 41391667 | G | A | 0.49893 | -0.21161 | 0.046572 | 5.53E-06 | 0.030112 |
| 15 | rs7179781 | 56931765 | A | T | 0.200148 | 0.278866 | 0.06139 | 5.56E-06 | 0.030241 |
| 16 | rs1437134 | 69730426 | A | G | 0.405943 | -0.23271 | 0.051252 | 5.61E-06 | 0.030493 |
| 16 | rs397891 | 89753031 | C | G | 0.417033 | 0.220568 | 0.048611 | 5.69E-06 | 0.03091 |
| 15 | rs2869552 | 78993600 | C | T | 0.210157 | -0.26784 | 0.059042 | 5.72E-06 | 0.031011 |
| 15 | rs4491476 | 79157405 | A | G | 0.41486 | 0.221211 | 0.048773 | 5.75E-06 | 0.031121 |
| 21 | rs2226697 | 22561893 | G | A | 0.448858 | -0.20538 | 0.045305 | 5.81E-06 | 0.031398 |
| 19 | rs7254188 | 41407343 | A | G | 0.443896 | 0.209954 | 0.04635 | 5.91E-06 | 0.031905 |

| | | | | | | | | | |
|---|---|---|---|---|---|---|---|---|---|
| 15 | rs11633519 | 78999552 | A | G | 0.228975 | -0.26155 | 0.057753 | 5.93E-06 | 0.031979 |
| 19 | rs3852870 | 41397370 | A | C | 0.498727 | -0.21096 | 0.046583 | 5.93E-06 | 0.031979 |
| 19 | rs55815263 | 41104977 | T | C | 0.168764 | -0.26943 | 0.059523 | 6.00E-06 | 0.03228 |
| 19 | rs10410750 | 41393937 | G | A | 0.498693 | -0.2107 | 0.046573 | 6.07E-06 | 0.032615 |
| 1 | rs35384162 | 32236033 | A | C | 0.035462 | 0.536719 | 0.118684 | 6.12E-06 | 0.032796 |
| 19 | rs28503746 | 41393135 | A | G | 0.498688 | -0.2106 | 0.046572 | 6.13E-06 | 0.032796 |
| 19 | rs3909341 | 41393326 | T | C | 0.498688 | -0.2106 | 0.046572 | 6.13E-06 | 0.032796 |
| 19 | rs4105142 | 41393570 | C | T | 0.498688 | -0.2106 | 0.046572 | 6.13E-06 | 0.032796 |
| 8 | rs1376442 | 42509597 | A | G | 0.134626 | -0.30578 | 0.067661 | 6.21E-06 | 0.033143 |
| 15 | rs10518871 | 57018224 | A | G | 0.165831 | 0.30505 | 0.067498 | 6.20E-06 | 0.033143 |
| 9 | rs3025360 | 1.36E+08 | A | G | 0.117662 | 0.307544 | 0.068057 | 6.22E-06 | 0.033145 |
| 21 | rs2826729 | 22562660 | A | G | 0.448637 | -0.20473 | 0.045306 | 6.22E-06 | 0.033145 |
| 19 | rs11667025 | 41480217 | C | G | 0.364414 | -0.21438 | 0.047494 | 6.37E-06 | 0.033889 |
| 20 | rs4809542 | 61986787 | G | C | 0.072018 | 0.387708 | 0.085903 | 6.38E-06 | 0.033927 |
| 1 | rs2297600 | 32207581 | G | T | 0.17409 | 0.262761 | 0.058354 | 6.70E-06 | 0.035595 |
| 16 | rs244415 | 69666683 | A | G | 0.40597 | -0.23059 | 0.051214 | 6.72E-06 | 0.035619 |
| 19 | rs11083580 | 41388836 | A | G | 0.498943 | 0.209605 | 0.046596 | 6.85E-06 | 0.036288 |
| 15 | rs7178196 | 57078278 | A | G | 0.165352 | 0.30645 | 0.068139 | 6.88E-06 | 0.0364 |
| 21 | rs2826724 | 22557624 | C | T | 0.448413 | -0.2033 | 0.045239 | 6.99E-06 | 0.036948 |
| 21 | rs8127329 | 22564279 | G | C | 0.450829 | -0.20345 | 0.045283 | 7.03E-06 | 0.037124 |
| 4 | rs13116922 | 1.3E+08 | T | C | 0.240967 | 0.230127 | 0.051233 | 7.06E-06 | 0.037245 |
| 15 | rs16977020 | 56919277 | C | A | 0.203269 | 0.274429 | 0.06111 | 7.10E-06 | 0.037383 |
| 9 | rs112270518 | 1.36E+08 | A | G | 0.117683 | 0.305566 | 0.068069 | 7.15E-06 | 0.037641 |
| 3 | rs6440994 | 1.55E+08 | T | C | 0.240162 | -0.2545 | 0.056716 | 7.22E-06 | 0.037934 |
| 15 | rs62012588 | 79032942 | A | G | 0.213705 | -0.2631 | 0.058647 | 7.25E-06 | 0.038069 |
| 21 | rs2826725 | 22559727 | G | T | 0.450009 | -0.20335 | 0.045341 | 7.30E-06 | 0.038263 |
| 16 | rs669696 | 69626136 | A | C | 0.407167 | -0.22941 | 0.051156 | 7.31E-06 | 0.038274 |
| 19 | rs10406188 | 41396915 | G | A | 0.442047 | 0.208167 | 0.046427 | 7.33E-06 | 0.038369 |
| 1 | rs113891242 | 32227413 | C | T | 0.036006 | 0.527585 | 0.117784 | 7.49E-06 | 0.039145 |
| 16 | rs862320 | 69651866 | T | C | 0.405776 | -0.22923 | 0.051201 | 7.57E-06 | 0.039499 |
| 7 | rs34997674 | 74120370 | A | G | 0.193013 | 0.252076 | 0.056314 | 7.60E-06 | 0.039617 |
| 1 | rs12033812 | 32206555 | C | A | 0.036452 | 0.523879 | 0.117102 | 7.69E-06 | 0.040047 |

| | | | | | | | | | |
|---|---|---|---|---|---|---|---|---|---|
| 19 | rs3865457 | 41398969 | T | C | 0.442074 | 0.207567 | 0.046437 | 7.83E-06 | 0.040729 |
| 1 | rs12739999 | 32207990 | A | G | 0.173589 | 0.261023 | 0.058448 | 7.97E-06 | 0.041438 |
| 1 | rs12749187 | 32205254 | G | T | 0.036785 | 0.519755 | 0.116422 | 8.03E-06 | 0.041639 |
| 15 | rs6493882 | 57016382 | A | G | 0.165927 | 0.301273 | 0.06748 | 8.02E-06 | 0.041639 |
| 19 | rs28472879 | 41395755 | A | G | 0.442144 | 0.207184 | 0.04643 | 8.11E-06 | 0.041996 |
| 19 | rs28427254 | 41388777 | A | G | 0.499233 | -0.20764 | 0.046544 | 8.15E-06 | 0.042155 |
| 19 | rs111769937 | 41172688 | T | C | 0.056344 | -0.42869 | 0.096103 | 8.17E-06 | 0.042219 |
| 21 | rs766090 | 22608321 | A | T | 0.417527 | -0.20629 | 0.046267 | 8.24E-06 | 0.042565 |
| 19 | rs10411264 | 41394336 | T | C | 0.442279 | 0.206778 | 0.046417 | 8.40E-06 | 0.043316 |
| 19 | rs8103288 | 41397499 | G | C | 0.442009 | 0.206774 | 0.046431 | 8.45E-06 | 0.043543 |
| 3 | rs12489801 | 1.82E+08 | A | T | 0.346437 | -0.20626 | 0.046321 | 8.47E-06 | 0.043606 |
| 19 | rs4803413 | 41485977 | A | G | 0.327286 | -0.21589 | 0.048501 | 8.54E-06 | 0.043895 |
| 19 | rs10414481 | 41398236 | T | C | 0.441983 | 0.206571 | 0.046433 | 8.63E-06 | 0.044333 |
| 4 | rs2391329 | 1.3E+08 | G | A | 0.243047 | 0.226659 | 0.050955 | 8.66E-06 | 0.044416 |
| 3 | rs28575712 | 1.55E+08 | T | C | 0.239413 | -0.25213 | 0.056697 | 8.71E-06 | 0.044609 |
| 1 | rs12028238 | 32228867 | A | C | 0.036095 | 0.522831 | 0.117656 | 8.84E-06 | 0.045193 |
| 21 | rs1041784 | 22615742 | T | C | 0.418313 | -0.20552 | 0.046248 | 8.84E-06 | 0.045193 |
| 3 | rs12486386 | 1.82E+08 | A | G | 0.346499 | -0.20577 | 0.04631 | 8.86E-06 | 0.045227 |
| 19 | rs4105141 | 41393600 | A | T | 0.44228 | 0.206114 | 0.046415 | 8.97E-06 | 0.045745 |
| 9 | rs117055611 | 14230974 | A | G | 0.134492 | 0.282029 | 0.06352 | 9.00E-06 | 0.045832 |
| 16 | rs2917677 | 69750849 | T | C | 0.406393 | -0.22752 | 0.051273 | 9.10E-06 | 0.046284 |
| 19 | rs5007415 | 41393760 | A | C | 0.442291 | 0.205984 | 0.046417 | 9.09E-06 | 0.046284 |
| 15 | rs6495318 | 78992137 | C | G | 0.212297 | -0.26092 | 0.058824 | 9.18E-06 | 0.046637 |
| 9 | rs118023793 | 14230975 | A | G | 0.134501 | 0.281625 | 0.063519 | 9.26E-06 | 0.046989 |
| 1 | rs2050256 | 32204683 | G | A | 0.172581 | 0.259582 | 0.05856 | 9.30E-06 | 0.047088 |
| 19 | rs4560023 | 41451893 | C | T | 0.381815 | 0.20838 | 0.047007 | 9.29E-06 | 0.047088 |
| 9 | rs4289905 | 91272656 | G | A | 0.142358 | -0.2863 | 0.064599 | 9.34E-06 | 0.047206 |
| 19 | rs10423695 | 41392141 | T | C | 0.499196 | -0.20654 | 0.046609 | 9.36E-06 | 0.04729 |
| 3 | rs6440995 | 1.55E+08 | T | A | 0.239759 | -0.25171 | 0.056807 | 9.38E-06 | 0.047333 |
| 4 | rs11097862 | 1.05E+08 | T | C | 0.360108 | 0.210758 | 0.047582 | 9.45E-06 | 0.047625 |
| 8 | rs7002242 | 42612683 | T | C | 0.127445 | -0.30951 | 0.06992 | 9.57E-06 | 0.048178 |
| 19 | rs2604861 | 41150922 | G | A | 0.376071 | -0.20532 | 0.046386 | 9.58E-06 | 0.048178 |

| | | | | | | | | | |
|---|---|---|---|---|---|---|---|---|---|
| 16 | rs244418 | 69622762 | A | G | 0.406176 | -0.22637 | 0.051145 | 9.60E-06 | 0.048243 |
| 9 | rs10961442 | 14236275 | A | G | 0.121202 | 0.293444 | 0.066322 | 9.67E-06 | 0.048506 |
| 19 | rs2260140 | 41391544 | T | C | 0.442227 | 0.205382 | 0.04643 | 9.71E-06 | 0.048677 |
| 9 | rs148428140 | 1.36E+08 | T | G | 0.117452 | 0.301393 | 0.068146 | 9.75E-06 | 0.048804 |
| 1 | rs34582656 | 32198423 | T | C | 0.036695 | 0.515136 | 0.116495 | 9.78E-06 | 0.048884 |
| 19 | rs1872122 | 41497997 | T | C | 0.326394 | -0.21474 | 0.048563 | 9.78E-06 | 0.048884 |
| 4 | rs28657372 | 1.05E+08 | G | A | 0.362471 | 0.210077 | 0.04752 | 9.83E-06 | 0.04908 |
| 16 | rs244417 | 69663519 | C | T | 0.406825 | -0.22649 | 0.051263 | 9.95E-06 | 0.049625 |
| 1 | rs6691283 | 32211454 | G | A | 0.036417 | 0.517134 | 0.11709 | 1.00E-05 | 0.049887 |
| 1 | rs71516323 | 32230280 | T | C | 0.036027 | 0.520135 | 0.11777 | 1.00E-05 | 0.049887 |
| 16 | rs7359336 | 69733460 | G | A | 0.407492 | -0.22654 | 0.051296 | 1.00E-05 | 0.049887 |
| 2 | rs7595779 | 1.48E+08 | T | C | 0.333034 | 0.214016 | 0.048466 | 1.01E-05 | 0.04996 |

**Table S4. Top Genes Identified after Correction for Multiple Testing**

| Gene Name | Chromosome | # of SNPs | ZSTAT | P-value | Q-value |
|---|---|---|---|---|---|
| IREB2 | 15 | 97 | 12.818 | 6.49E-38 | 8.94E-34 |
| CHRNA3 | 15 | 56 | 11.657 | 1.06E-31 | 7.29E-28 |
| CHRNA5 [a] | 15 | 39 | 10.747 | 3.05E-27 | 1.40E-23 |
| HYKK | 15 | 14 | 8.6195 | 3.36E-18 | 1.16E-14 |
| ADAMTS7 | 15 | 72 | 8.5805 | 4.72E-18 | 1.30E-14 |
| CHRNB4 [a] | 15 | 3 | 8.2885 | 5.73E-17 | 1.32E-13 |
| CYP2A6 | 19 | 13 | 7.7569 | 4.35E-15 | 8.57E-12 |
| RAB4B [a] | 19 | 47 | 7.1145 | 5.62E-13 | 9.67E-10 |
| PSMA4 | 15 | 17 | 6.4485 | 5.65E-11 | 8.65E-08 |
| EGLN2 | 19 | 12 | 6.0349 | 7.96E-10 | 1.10E-06 |
| CHRNA6 | 8 | 18 | 5.3396 | 4.66E-08 | 5.83E-05 |
| C19orf54 | 19 | 10 | 4.9707 | 3.34E-07 | 3.83E-04 |
| ADGRB2 | 1 | 23 | 4.9266 | 4.18E-07 | 4.43E-04 |
| LTBP4 | 19 | 62 | 4.7345 | 1.10E-06 | 1.08E-03 |
| MORF4L1 | 15 | 158 | 4.6981 | 1.31E-06 | 1.16E-03 |
| CXXC4 | 4 | 7 | 4.6925 | 1.35E-06 | 1.16E-03 |
| CHRNA4 [a] | 20 | 95 | 4.6583 | 1.59E-06 | 1.29E-03 |
| NUMBL | 19 | 48 | 4.4902 | 3.56E-06 | 2.72E-03 |
| GTF2I | 7 | 73 | 4.2133 | 1.26E-05 | 9.13E-03 |
| TOX2 | 20 | 168 | 4.0601 | 2.45E-05 | 1.69E-02 |
| APH1A | 1 | 1 | 4.0133 | 2.99E-05 | 1.96E-02 |
| SNRPA | 19 | 13 | 3.9841 | 3.39E-05 | 2.12E-02 |
| ADCK4 | 19 | 33 | 3.9631 | 3.70E-05 | 2.22E-02 |
| RAPSN | 11 | 29 | 3.9259 | 4.32E-05 | 2.47E-02 |
| CYP2A7 | 19 | 46 | 3.9172 | 4.48E-05 | 2.47E-02 |
| TIMM17A | 1 | 12 | 3.8678 | 5.49E-05 | 2.91E-02 |
| ZBTB7A | 19 | 18 | 3.8417 | 6.11E-05 | 3.12E-02 |

| | | | | | |
|---|---|---|---|---|---|
| XRCC3 | 14 | 22 | 3.8057 | 7.07E-05 | 3.48E-02 |
| SPATA33 | 16 | 38 | 3.7718 | 8.10E-05 | 3.85E-02 |
| BNC2 | 9 | 753 | 3.7538 | 8.71E-05 | 4.00E-02 |
| ADD1 | 4 | 61 | 3.7408 | 9.17E-05 | 4.08E-02 |

Notation: [a] indicates gene that was included in the transcriptionally regulated set of genes tested.

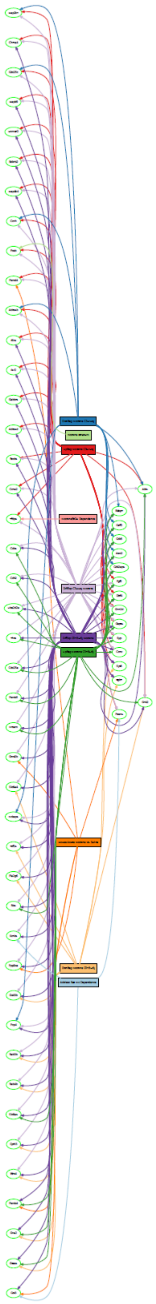

**Figure S1. Gene set Graph showing representation of genes across gene sets identified in Table 1 from five studies**. Citation: Plots of the relationship between buffer length and the calculated enrichment values explained by the buffer-length model components. Enrichment values are calculated across all 3 folds to be representative of the entire sample, not by-fold. Lines shown reflect inferred trends for buffer lengths not assessed.

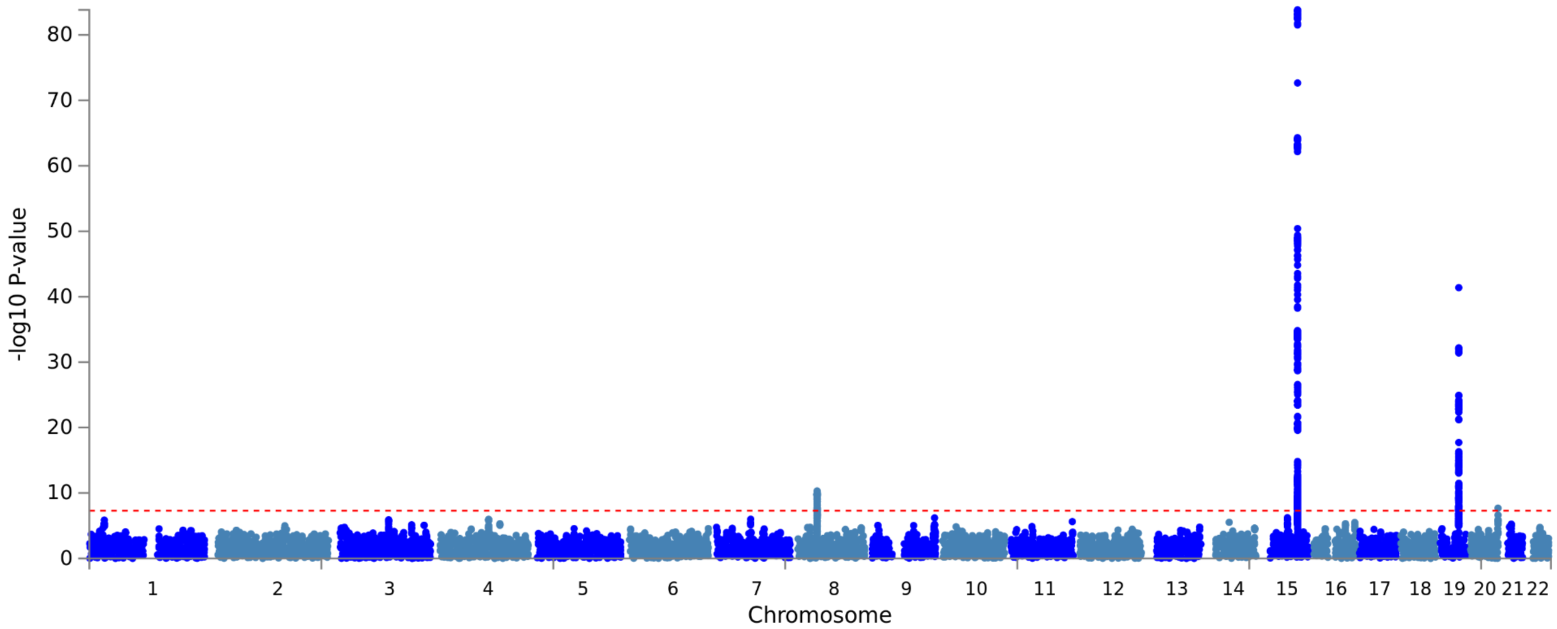

**<u>Figure S2. Manhattan plot of tobacco consumption GWAS.</u>**

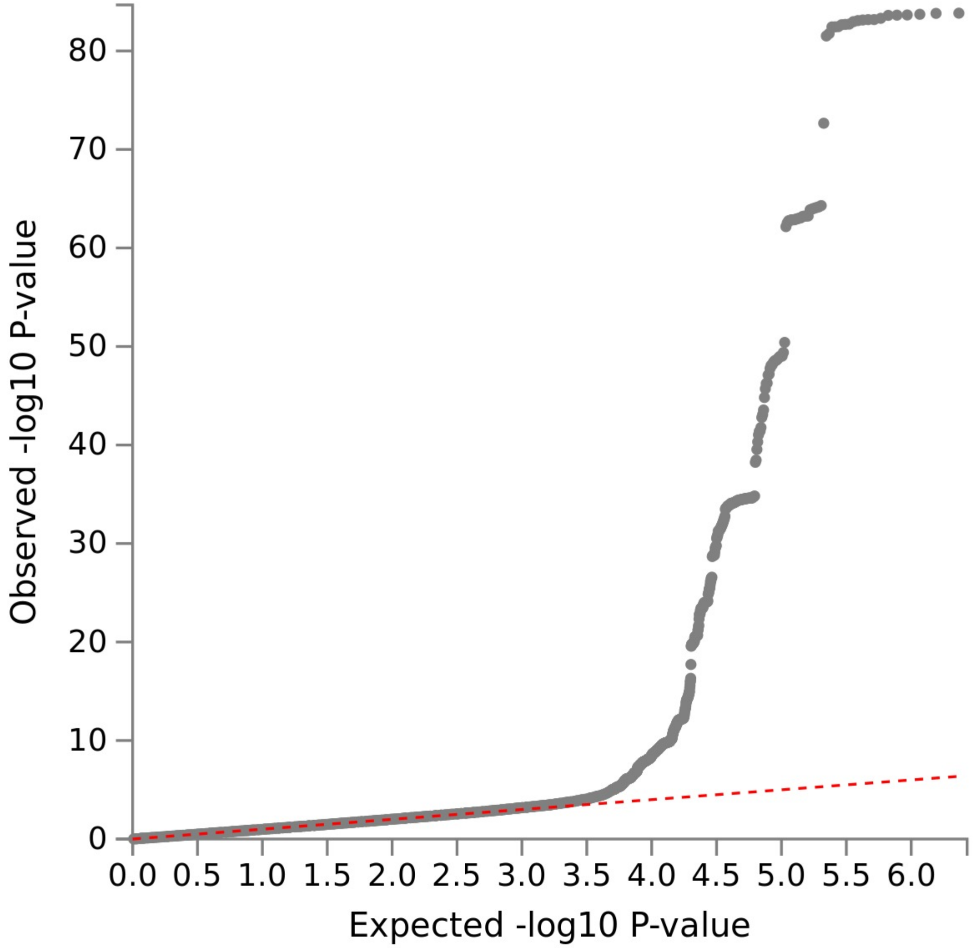

**Figure S3: GWAS-QQ plot**

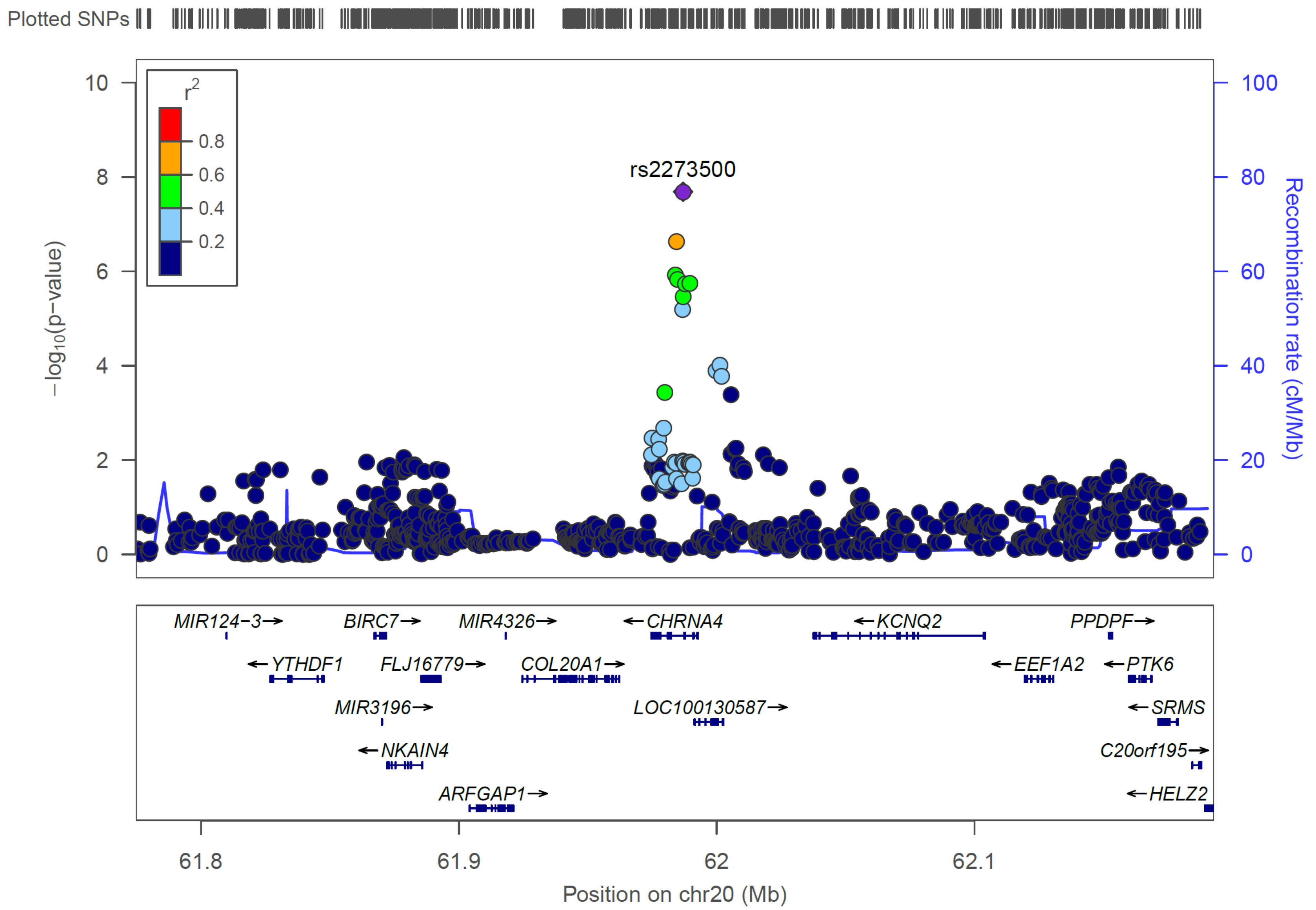

**Figure S4: GWAS regional association plot of CHRNA4 on chromosome 20**

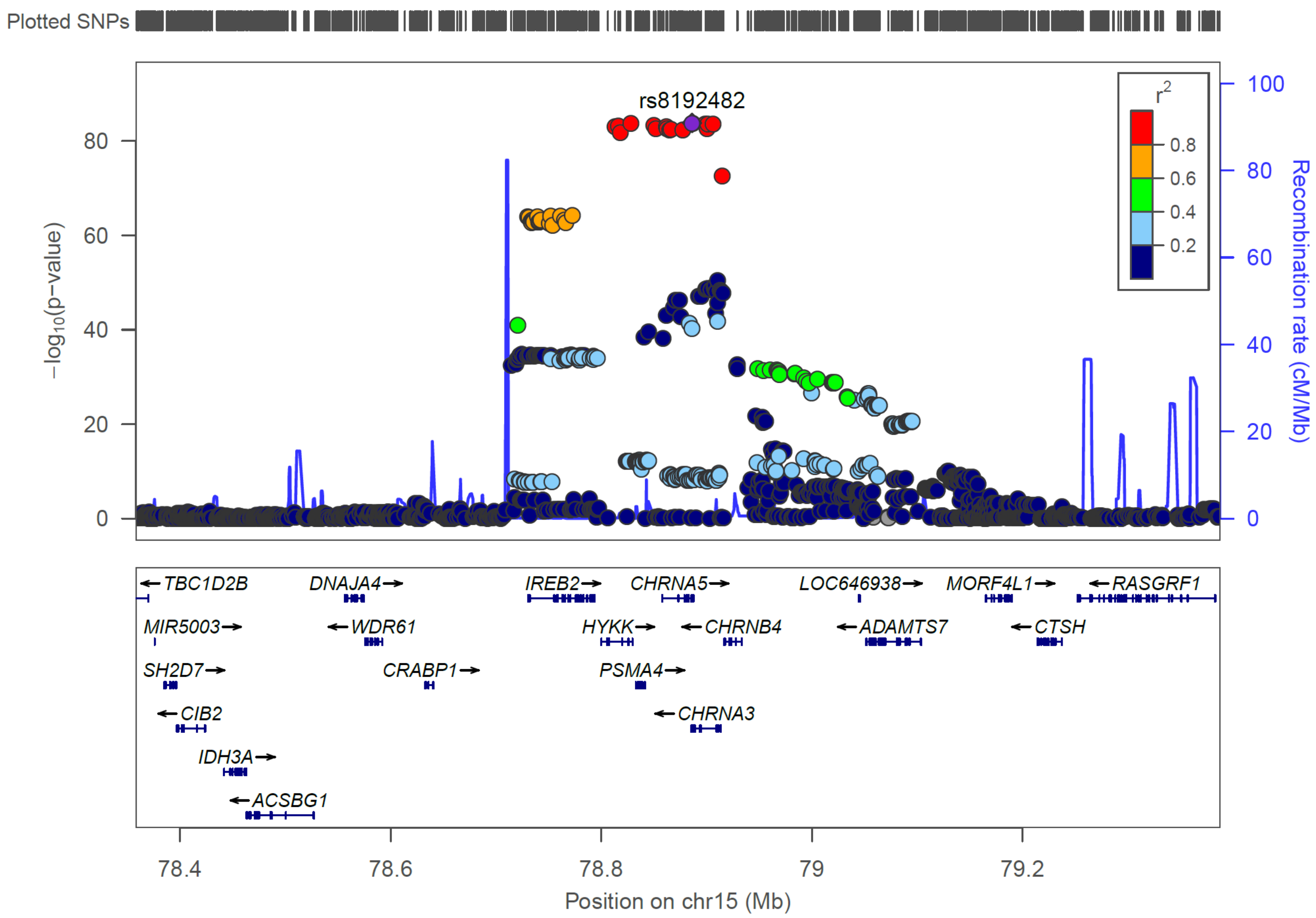

**Figure S5: GWAS regional association plot of CHRNA5 on chromosome 15**

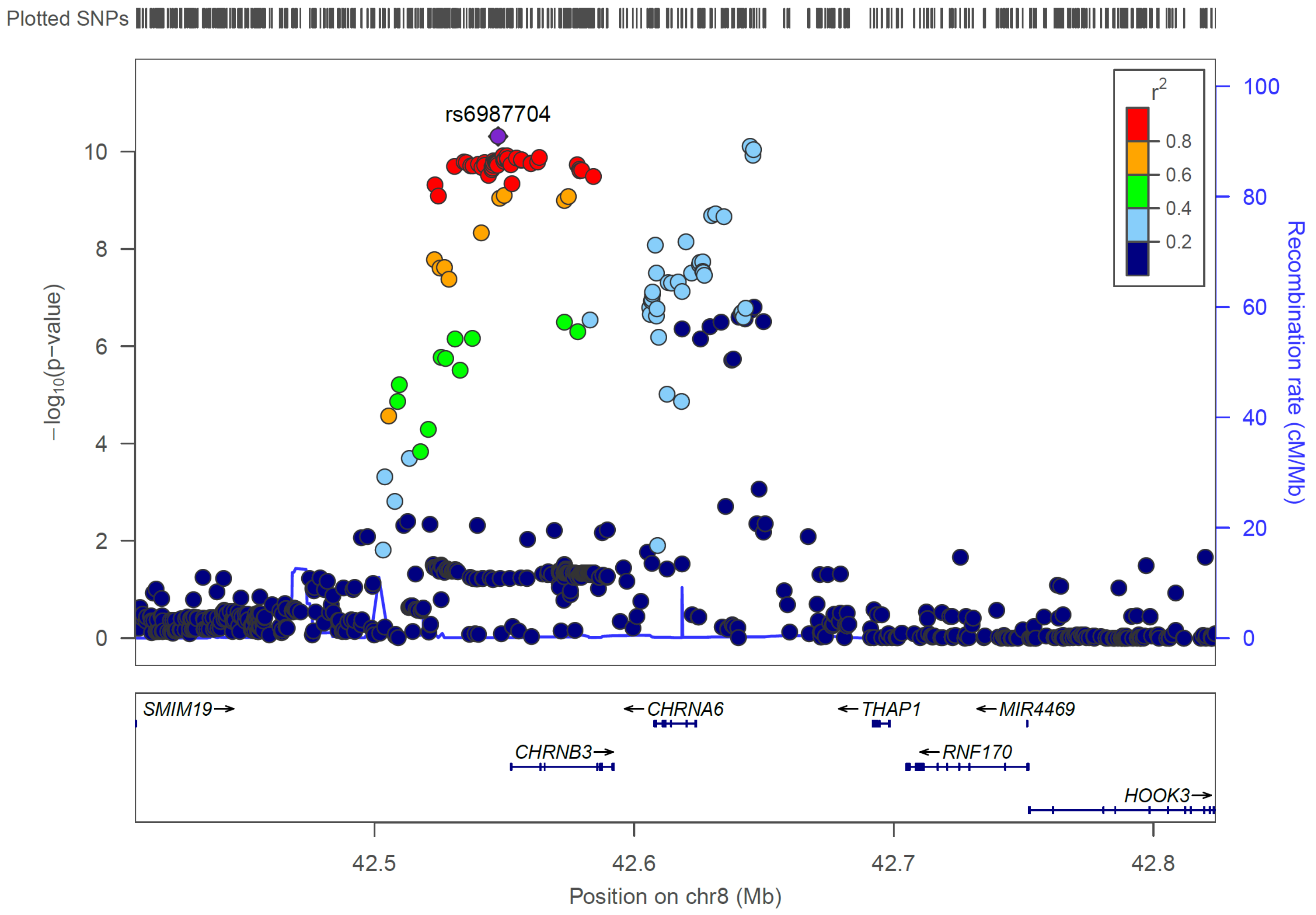

**Figure S6: GWAS regional association plot of CHRNA6 on chromosome 8**

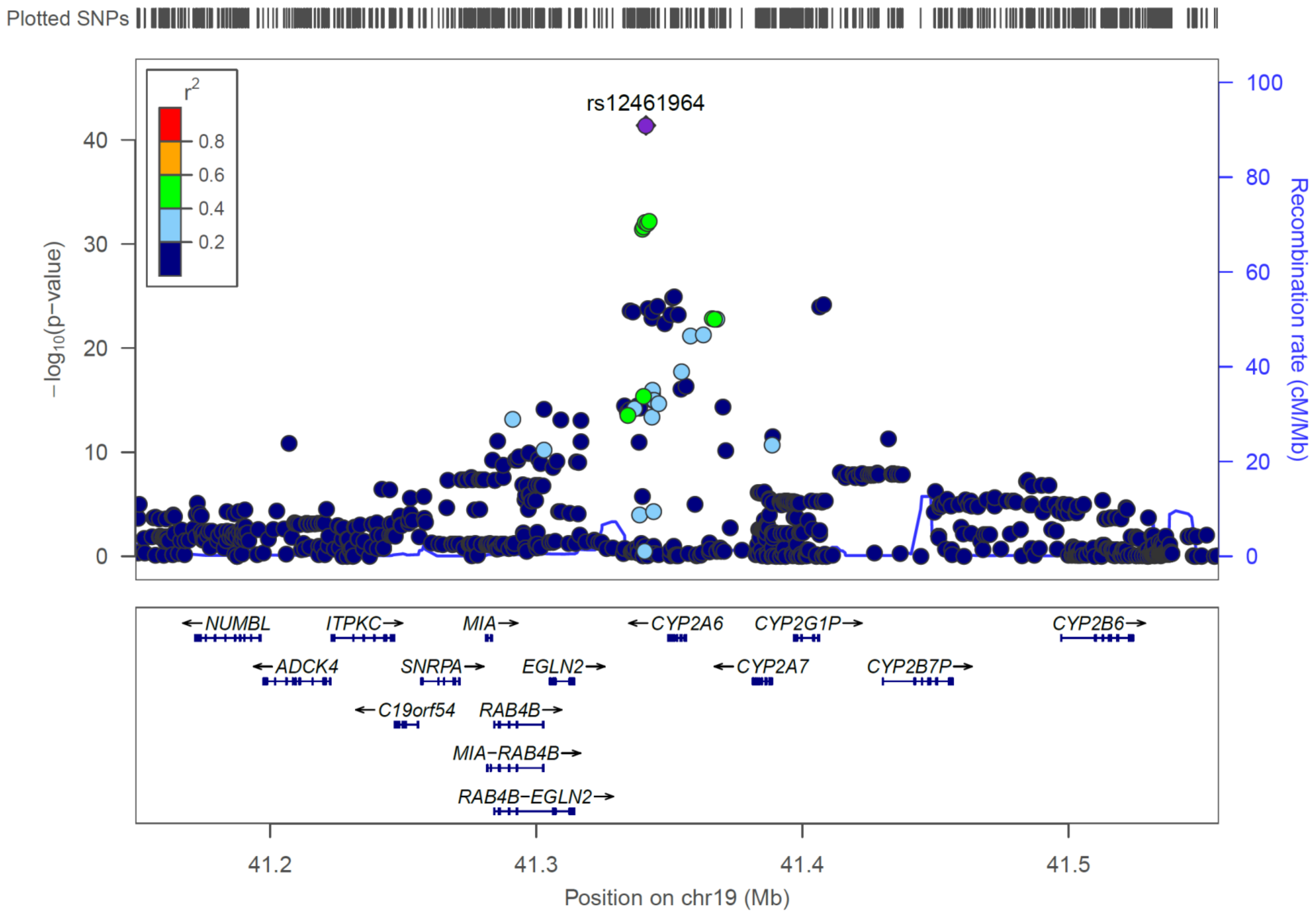

**Figure S7: GWAS regional association plot of CYP2A6 on chromosome 19**

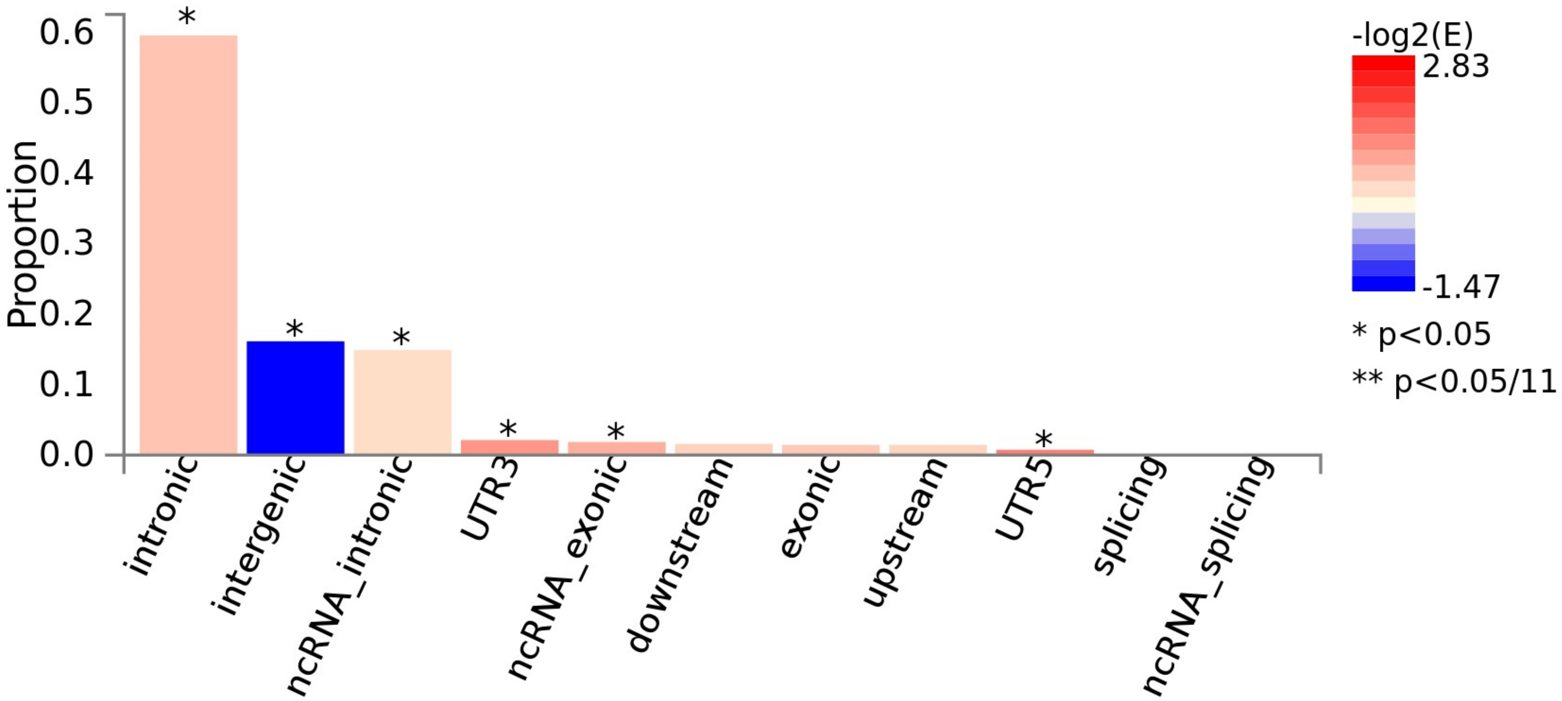

**Figure S8: GWAS SNP Annotation plot**

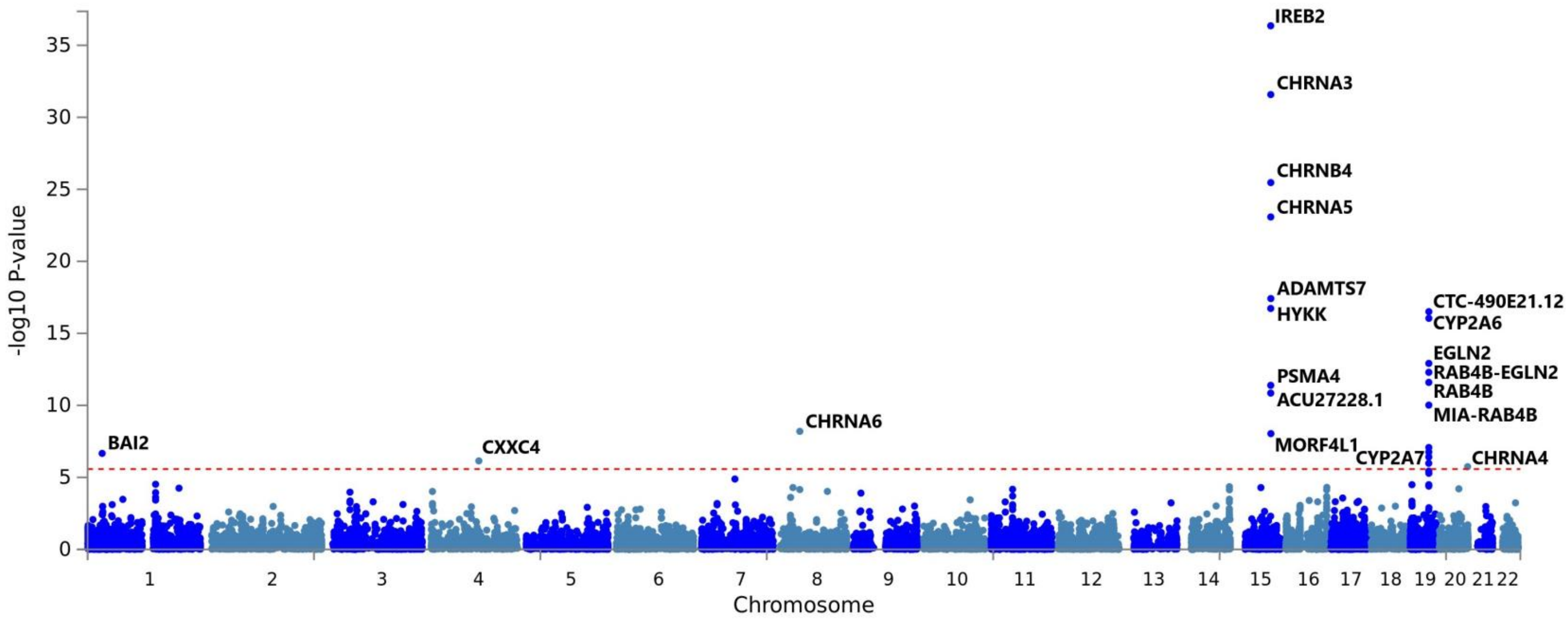

**Figure S9: Gene-based test Manhattan Plot**

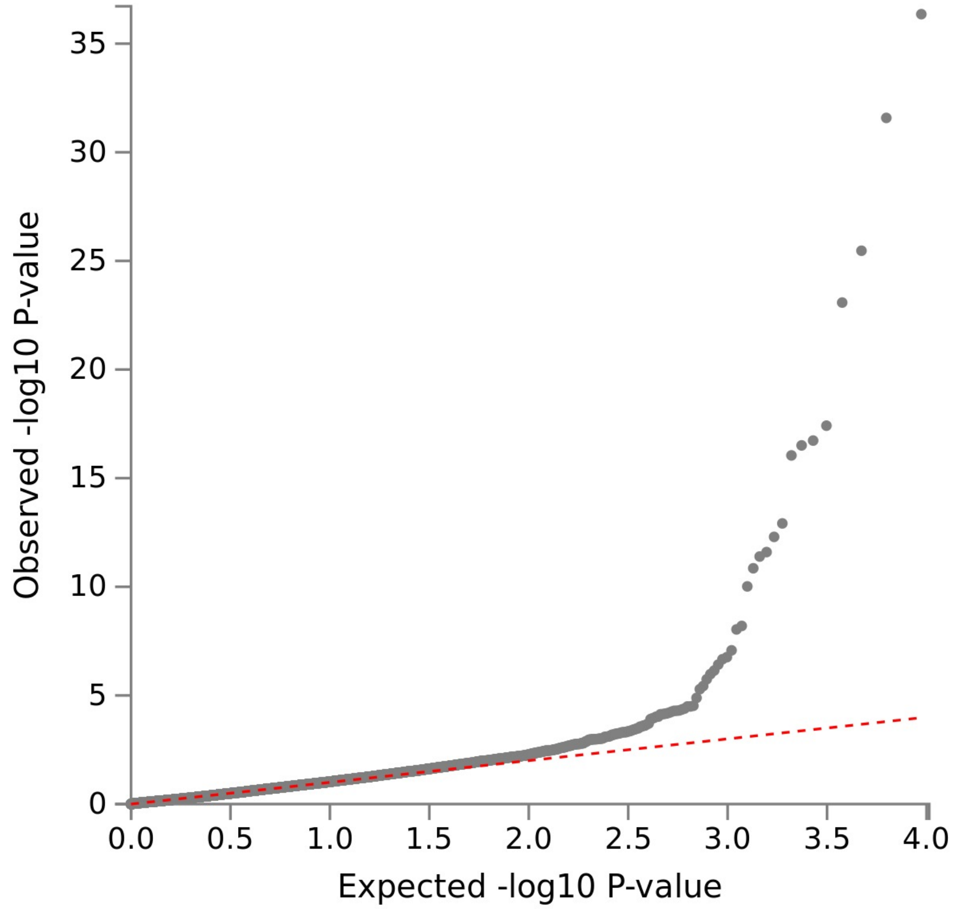

**Figure S10: Gene-based test QQ Plot**

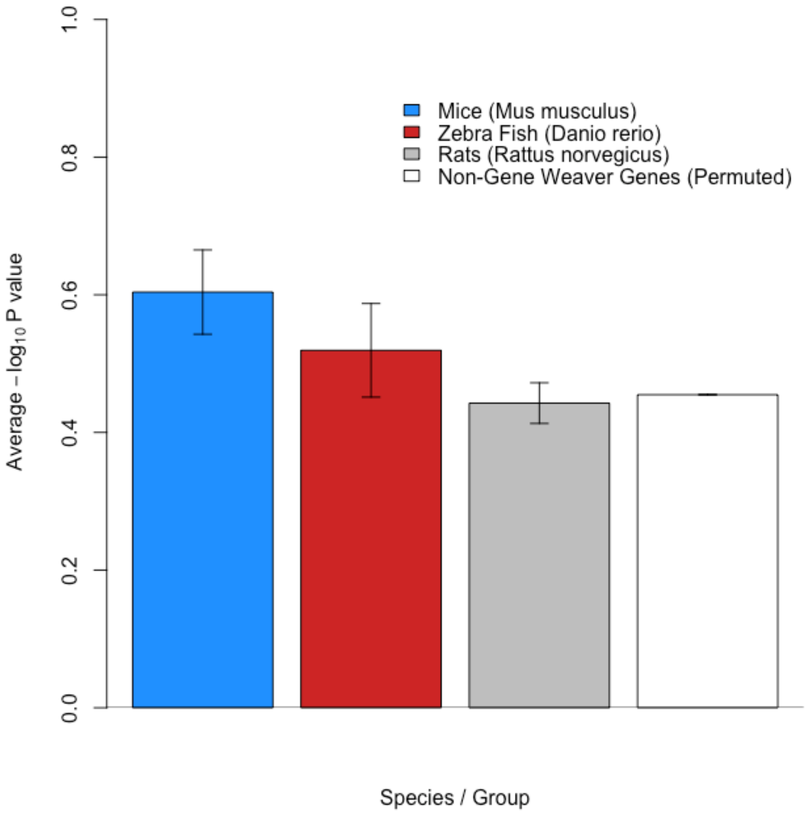

**Figure S11. Cross–Species comparison of human gene–based test on tobacco use**. Citation: Comparison of the –log10 p-value across contributing gene sets was conducted using Welch Two Sample t-tests (Mus musculus:Danio rerio- t=0.9244, df=225.54, p=0.3563; Mus musculus:Rattus norvegicus-t=2.3719, df=833.63, p=0.01792). Note we re-sampled the average –log10 p-value (10,000 permutations) of a list of 500 genes outside of our Gene Weaver list (Non-Gene Weaver Genes) to determine whether the genes from our nicotine Gene Weaver lists were more significant in our human gene-based test than we would expect by chance. Only the gene list from mice was significantly greater than this random permuted list of non-Gene Weaver genes (Mus musculus:Non-Gene Weaver Genes- t=2.4342, df=640.03, p=0.0152).
